# Phenylphenalenones and Linear Diarylheptanoid Derivatives are Biosynthesized via Parallel Routes in *Musella lasiocarpa*, the Chinese Dwarf Banana

**DOI:** 10.1101/2024.03.11.584385

**Authors:** Hui Lyu, Lukas Ernst, Yoko Nakamura, Yu Okamura, Tobias G. Köllner, Katrin Luck, Benye Liu, Yu Chen, Ludger Beerhues, Jonathan Gershenzon, Christian Paetz

**Affiliations:** NMR/Biosynthesis Group, Max Planck Institute for Chemical Ecology, Jena 07745, Germany; Technische Universität Braunschweig, Institute of Pharmaceutical Biology, Braunschweig 38106, Germany; Department of Biochemistry, Max Planck Institute for Chemical Ecology, Jena 07745, Germany; Department of Natural Product Biosynthesis, Max Planck Institute for Chemical Ecology, Jena 07745, Germany; Jiangsu Key Laboratory for the Research and Utilization of Plant Resources, Institute of Botany, Jiangsu Province and Chinese Academy of Sciences (Nanjing Botanical Garden Mem. Sun Yat-Sen), Nanjing 210014, China; Department of Insect Symbiosis, Max Planck Institute for Chemical Ecology, Jena 07745, Germany; Department of Biological Sciences, Graduate School of Science, University of Tokyo, Tokyo 113-0033, Japan

## Abstract

Phenylphenalenones (PPs) are complex polycyclic natural products that play an important role in the chemical defense system of banana and plantain (Musaceae). Although suggestions for how plants synthesize the PP scaffold were first proposed more than 50 years ago, no biosynthetic information is yet available at the enzyme level. Here, we use transcriptomic data from seeds of *Musella lasiocarpa*, the Chinese dwarf banana, to identify five biosynthetic genes involved in the formation of dihydrocurcuminoids. Characterization of the substrate specificities of the enzymes reveals two distinct dihydrocurcuminoid pathways leading to the two types of major aromatic seed metabolites, the PPs and the linear diarylheptanoid derivatives. Furthermore, through multiple rounds of feeding potential intermediates to *M. lasiocarpa* root protein extract, followed by high-resolution mass spectrometry profiling, product isolation, and NMR-based elucidation, we demonstrate the stepwise conversion of a dihydrocurcumin-type precursor to the PP 4’-hydroxylachnanthocarpone. In contrast to the commonly hypothesized Diels–Alder cyclization mechanism, we propose an unexpected two-step cyclization route to the PP scaffold.

---

Phenylphenalenones (PPs) are structurally diverse polycyclic aromatic natural products with a tricyclic phenalene-1*H*-one nucleus and an attached phenyl group (Scheme 1).^[1]^ They are constituents of plants in the monocotyledonous families Haemodoraceae, Pontederiaceae, Strelitziaceae, and Musaceae,^[2]^ including perennial herbs and trees such as kangaroo paws (*Anigozanthos* spp.), water hyacinths (*Pontederia* spp.), bird of paradise (*Strelitzia* spp.), and banana plants (*Musa* spp.). Although the ecological function of PPs is not fully understood, there is evidence that Musaceae species biosynthesize PPs as phytoalexins against pathogens such as *Fusarium oxysporum* f. sp. *cubense* (causing Panama disease),^[3]^ *Pseudocercospora fijiensis* (causing Black Sigatoka disease),^[4]^ and the nematode *Radopholus similis* (causing blackhead toppling disease).^[5]^ Notably, a previous study linked the increased accumulation of PPs to the increased viability of disease-resistant banana varieties.^[5]^ However, despite numerous breeding and hybridization efforts,^[6]^ prevention of fungal diseases in commercial banana varieties still relies on the extensive use of pesticides.^[7]^ Thus, understanding the biosynthetic formation of PPs may open new opportunities to engineer the natural defense mechanisms of banana plants.

A partial biosynthetic hypothesis for PPs first proposed in 1961^[8]^ was later validated through feeding experiments with radiotracer and stable isotope precursors, leading to the proposed biosynthetic pathway depicted in Scheme 1.^[1]^ In the initial stage of this biosynthetic scenario, a linear diarylheptanoid (DH) is formed by the successive condensation of two phenylpropanoids with a malonate unit.^[9]^ After the A-ring hydroxylation of the linear DH intermediate, a hypothetical intramolecular Diels–Alder cyclization step yields the PP scaffold.^[10]^ The remarkable structural diversity observed for PPs is primarily attributed to their varying oxygenation patterns, yet nearly all PPs feature a 1,2-dioxygenated A-ring.^[11]^ The carbonyl function at C-1 and, if present, the hydroxy group at C-6 of the PPs are derived from *p*-coumaric acid, which was the preferred substrate in studies using stable isotope labeled precursors.^[12]^ The hydroxy group at C-2 is introduced into the linear DH precursor from atmospheric O_2_.^[11]^ However, most of the intermediates of PP biosynthesis are still unknown and none of the genes involved in the pathway have been identified.

**Scheme 1.**
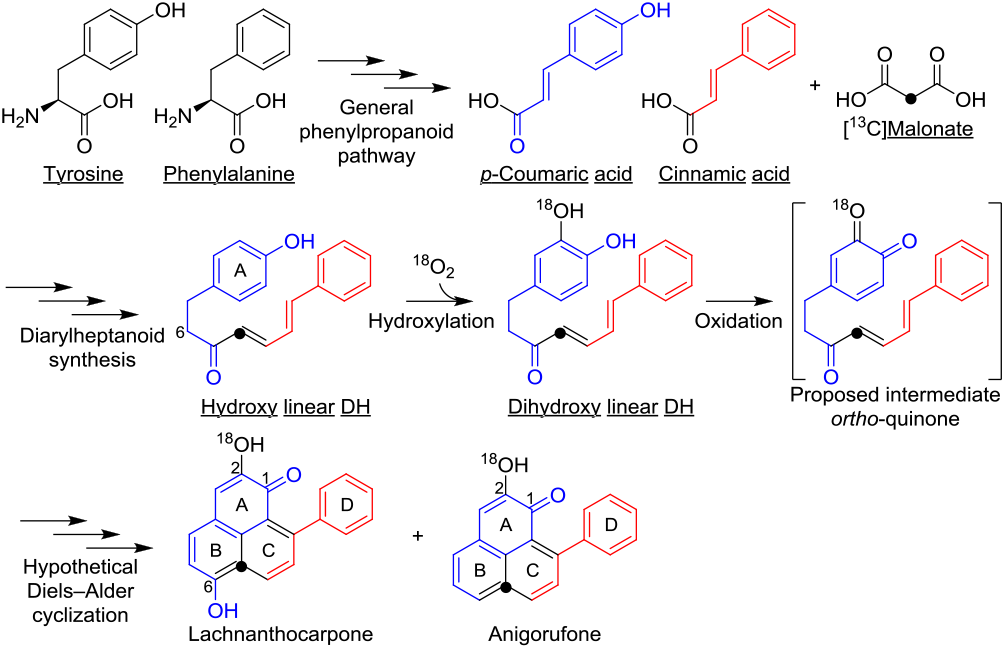
The proposed biosynthetic pathway of phenylphenalenones. The underlined biosynthetic precursors of lachnanthocarpone and anigorufone were identified by isotope labeling experiments. The indicated carbon atom (black dot) of linear DH is derived from ^13^C-labeled malonate during diarylheptanoid synthesis. The order of oxygenation events was probed by studies with isotopic oxygen (^18^O_2_).

In a recent study, we elucidated the structures of various PPs and linear DH derivatives found in the seeds of the Chinese dwarf banana (*Musella lasiocarpa*, Musaceae), which were present at later stages (brown and black seeds) but absent at early stages of development (yellow seeds) (Figure 1A).^[13]^ Based on their structural features, we hypothesized that the assembly of both compound classes likely shares the initial steps of dihydrocurcuminoid biosynthesis and involves two different starter substrates (*p*-coumaroyl- and feruloyl-CoA; Figure S1). Here, we perform a *de novo* transcriptome assembly and compare gene expression across all three stages to identify five biosynthetic genes involved in the formation of the two central dihydrocurcumin-type intermediates. We use feeding experiments with *M. lasiocarpa* protein extracts to show many of the steps involved in the conversion of the *p*-coumaroyl-derived dihydrocurcuminoid into the PP 4’-hydroxylachnanthocarpone.

**Figure 1.**
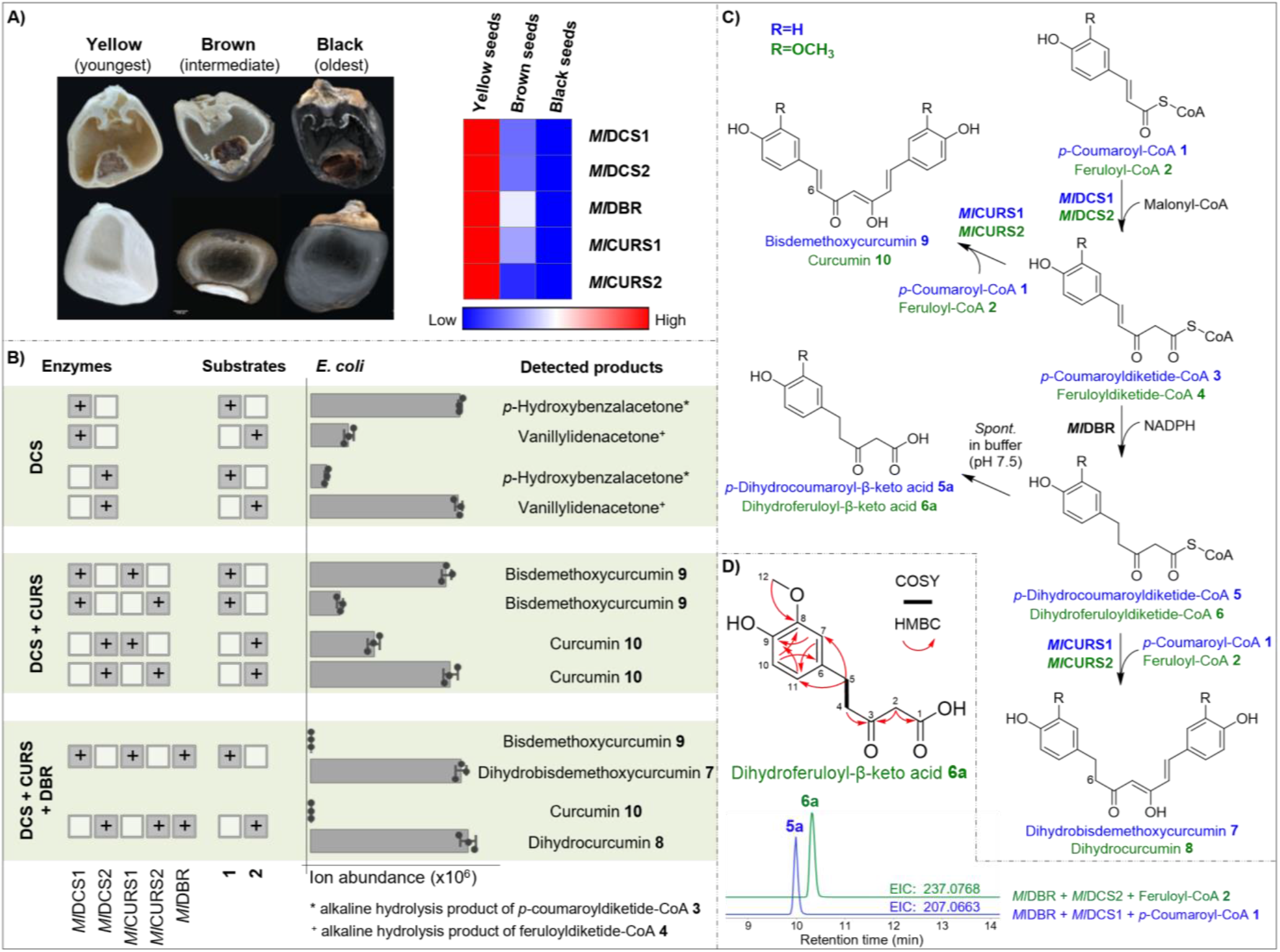
Biosynthesis of dihydrocurcuminoids in *M. lasiocarpa* seeds. (A) Left: Three developmental stages of *M. lasiocarpa* seeds were used to isolate RNA for subsequent RNA-seq analysis. Right: Expression profiles of the identified genes representing the FPKM values in the seed transcriptomes. (B) The LCMS peak area of products detected in *in vitro* assays using the indicated combinations of purified recombinant enzymes and substrates. Diketide-CoA esters were detected after alkaline hydrolysis. Data are mean ± s.e.m. (n = 3). (C) The biosynthetic pathway for the formation of dihydrocurcuminoids (dihydrobisdemethoxycurcumin **7** and dihydrocurcumin **8**) and curcuminoids (bisdemethoxycurcumin **9** and curcumin **10**). (D) Extracted ion chromatograms (EIC) of the spontaneously hydrolyzed dihydrodiketide-CoA products, *p*-dihydrocoumaroyl-β-keto acid **5a** and dihydroferuloyl-β-keto acid **6a**. Key COSY and HMBC correlations of the previously undescribed structure of **6a** are shown above. Compound names in blue, R = H; compound names in green, R = OCH_3_.

The biosynthesis of linear DH scaffolds has already been studied in several plant species.^[14,15]^ Curcuminoids, bis-α,β-unsaturated tautomeric β-diketone linear DHs (Figure S2), are well-known constituents of turmeric (*Curcuma longa*, Zingiberaceae).^[16]^ They are formed in a two-step process catalyzed by the sequential action of the type III polyketide synthases diketide-CoA synthase (DCS) and curcumin synthase (CURS) (Figure S3A).^[14]^ First, the DCS condenses feruloyl-CoA **2** and malonyl-CoA to form feruloyldiketide-CoA **4**. Subsequently, the CURS hydrolyzes feruloyldiketide-CoA **4** into feruloyl-β-keto acid and catalyzes its coupling with another molecule of feruloyl-CoA **2** to form curcumin **10**. In contrast, another curcuminoid synthase isolated from rice (*Oryza sativa*, Poaceae) catalyzed the formation of bisdemethoxycurcumin **9** by condensation of two molecules of *p*-coumaroyl-CoA **1** and one molecule of malonyl-CoA (Figure S3B).^[15]^ Since *C. longa* is phylogenetically more closely related to *M. lasiocarpa* than to *O. sativa* (Figure S4),^[17]^ a translated nucleotide search (tBLASTn) using the turmeric DCS and CURS as queries against the *M. lasiocarpa* seed transcriptome led to the identification of two putative DCS sequences (*Ml*DCS1 and *Ml*DCS2) and two CURS homologs (*Ml*CURS1 and *Ml*CURS2), which all were highly expressed in yellow seeds of *M. lasiocarpa* (Figure 1A).

The catalytic activities of *Ml*DCS1 and *Ml*DCS2 were established in assays using purified enzymes overexpressed in *Escherichia coli*. Both enzymes accepted various phenylpropanoid-CoA esters as starter substrates and used one molecule of malonyl-CoA as the extender substrate to catalyze the formation of the corresponding diketide-CoA products (Figure S5). The benzalacetone derivatives were detected after alkaline hydrolysis using liquid chromatography–mass spectrometry (LCMS). Product identities were confirmed by comparing their retention times and MS/MS fragmentation patterns with those of commercial standards. Interestingly, the substrate specificities of the two enzymes differed. While *Ml*DCS1 clearly preferred *p*-coumaroyl-CoA **1** as the starter substrate, *Ml*DCS2 favored feruloyl-CoA **2** (Figure 1B). To test the catalytic function of the enzymes in a plant environment, *Nicotiana benthamiana* leaves were transiently transformed via *Agrobacterium*-mediated infiltration. LCMS analysis of alkaline-hydrolyzed leaf extracts revealed that plants expressing *Ml*DCS1 exclusively formed *p*-coumaroyldiketide-CoA **3**, while feruloyldiketide-CoA **4** was the only product formed in plants expressing *Ml*DCS2 (Figure S5).

The enzymatic functions of *Ml*CURS1 and *Ml*CURS2 were assessed in *in situ* co-incubations with purified *Ml*DCS1 or *Ml*DCS2 in the presence of their corresponding preferred starter substrates, *p*-coumaroyl-CoA **1** and feruloyl-CoA **2**. The two CURS enzymes were able to catalyze the decarboxylative condensation of the diketide-CoA intermediates **3** or **4** with another molecule of the substrates **1** or **2**, respectively, to yield the curcuminoids bisdemethoxycurcumin **9** and curcumin **10**, respectively (Figure 1C). However, *Ml*CURS1 showed higher activity in combination with *Ml*DCS1 and starter substrate **1**, while *Ml*CURS2 performed better with *Ml*DCS2 and substrate **2** (Figure 1B). These findings were further substantiated through co-infiltration experiments with different combinations of DCS and CURS in *N. benthamiana* leaves, in which *Ml*CURS1 and *Ml*CURS2 exhibited identical activities and substrate preferences to those after *E. coli* expression (Figure S6 and Figure S7).

Given that curcuminoids **9** and **10** possess a Δ^6^ double bond, while the putative PP precursors (Scheme 1) and all isolated linear DH derivatives^13^ are saturated at C-6, reduction of the Δ^6^ double bond may be a shared step in the biosynthesis of both PPs and linear DH derivatives (Figure S1). It could occur at either the phenylpropanoid-CoA, diketide-CoA, or curcuminoid stage (Figure S8). We searched for double bond reductases (DBRs) that could act on double bonds in conjugation with carbonyl functions in the *de novo* transcriptome assembly of *M. lasiocarpa*. A total of nine homologs were annotated as NADPH-dependent 2-alkenal reductases. However, only one of them, named *Ml*DBR, showed an expression pattern similar to those of *Ml*DCS and *Ml*CURS (Figure 1A). Following overexpression in *E. coli* and affinity-purification of the recombinant protein, its enzymatic activity was initially tested against *p*-coumaroyl-CoA **1**, feruloyl-CoA **2**, bisdemethoxycurcumin **9**, and curcumin **10**. However, no conversion was detected for any of the phenylpropanoid-CoA and curcuminoid substrates using NADH and NADPH as putative cofactors.

We next tested diketide-CoAs as potential substrates by incubating *Ml*DBR with *Ml*DCS1 or *Ml*DCS2 in the presence of their respective starter substrates **1** and **2**, malonyl-CoA, and NAD(P)H. A mass peak at *m/z* 237.0768 [M – H]^−^ (calculated C_12_H_13_O_5_^-^) was detected in assays comprising *Ml*DBR, *Ml*DCS2, substrate **2**, and NADPH (Figure 1D). The product was isolated from large-scale incubations and its structure was elucidated by NMR spectroscopy (Figure S23–S26 and Table S1), revealing the previously undescribed dihydroferuloyl-β-keto acid **6a** (Figure 1D). The corresponding product, *p*-dihydrocoumaroyl-β-keto acid **5a**, was identified in co-incubations containing *Ml*DBR, *Ml*DCS1, substrate **1**, and NADPH through detection of an ion at *m/z* 207.0663 [M – H]^−^. The recorded mass was consistent with the loss of a methoxy group (30 Da) from compound **6a**, and comparison of MS/MS fragmentation patterns further supported this identification (Figure S9C). Spontaneous hydrolysis of dihydroferuloyldiketide-CoA **6** to **6a** in pH 7.5 buffer was expected and underlined by a linear correlation between the formation of **6a** and the incubation time of the assay (from 1 h to 10 h; Figure S9D).

The addition of *Ml*DBR to the previously conducted co-incubation assays of *Ml*DCS1/*Ml*CURS1 and *Ml*DCS2/*Ml*CURS2 led to the formation of the expected dihydrocurcuminoid products dihydrobisdemethoxycurcumin **7** and dihydrocurcumin **8**, respectively (Figure 1B and Figure S6). No traces of curcuminoids **9** and **10** were observed in the assays, indicating the efficiency of this reduction step. These results were consistently replicated through co-infiltration experiments in *N. benthamiana* using the same enzyme combinations (Figure S6 and Figure S7). For the confirmation of product identities, dihydrocurcumin **8** was compared to a commercially available standard, while dihydrobisdemethoxycurcumin **7** was obtained from chemical synthesis (see Supporting Information for synthetic procedures).

We therefore characterize *Ml*DBR as an unusual diketide-CoA-accepting member of the zinc-independent medium-chain dehydrogenase/reductase superfamily, containing two conserved NADPH-binding motifs AXXGXXG and GXXS (Figure S10).^[18]^ Examples of plant DBRs catalyzing similar reactions include the NADPH-dependent hydroxycinnamoyl-CoA reductase isolated from apple (*Malus domestica*, Rosaceae; Figure S11A).^[19]^ However, a recent study has raised doubts about the reproducibility of these results, as the identical enzyme was reported to act on benzalacetone and *p*-coumaroyl aldehyde substrates (Figure S11B).^[20]^ Two curcuminoid-specific reductases have recently been isolated from *Alnus sieboldiana* (Betulaceae).^[21]^ These enzymes catalyze the two-step NADPH-dependent reduction of curcumin **10** to dihydrocurcumin **8** and then further to tetrahydrocurcumin (Figure S11C). Notably, no reductase has been described that accepts diketide-CoAs as substrates.

We hypothesized that *Ml*DBR participates in two distinct enzymatic constellations in the formation of aromatic compounds in *M. lasiocarpa* seeds. One constellation (*Ml*DCS1/*Ml*DBR/*Ml*CURS1) is responsible for the production of dihydrobisdemethoxycurcumin **7**, the proposed precursor of *M. lasiocarpa* PPs, while the other constellation (*Ml*DCS2/*Ml*DBR/*Ml*CURS2) is responsible for the formation of dihydrocurcumin **8**, the precursor of various linear DH derivatives.

The PP skeleton was previously proposed to be formed from a trihydroxy linear DH like **12** via a reactive *ortho*-quinone intermediate that undergoes intramolecular Diels–Alder-like cyclization to yield PPs (Figure S12).^[10,13]^ Of the intermediates formed in the above reactions, we considered dihydrobisdemethoxycurcumin **7** a likely precursor of **12** because of its ring substitution pattern (Figure 2). The conversion of **7** to **12** could proceed via reduction and dehydration to the linear DH **11** with an α,β,γ,δ-unsaturated ketone function, followed by hydroxylation on the A ring at the C-3’’ position (Figure 2). To demonstrate this conversion, we employed protein extracts from *M. lasiocarpa*. Due to the limited amount of fresh seed material available and the difficulties in preparing cell-free homogenates from the hard seeds, we explored *M. lasiocarpa* roots as a possible alternative. The outermost root layer proved to be another rich source of aromatic metabolites, including trace amounts of dihydrobisdemethoxycurcumin **7** (Figure S13). Incubation of 2’,3’,5’,6’-deuterium-labeled **7** (*d*_4_-**7**, see Supporting Information for synthetic procedures) with NADPH in the crude root protein extracts resulted in the appearance of two products with mass peaks at *m/z* 299.1580 and *m/z* 315.1529 [M + H]^+^, matching the expected masses of labeled 4’,4’’-dihydroxy linear DH *d*_4_-**11** and 4’,3’’,4’’-trihydroxy linear DH *d*_4_-**12** (peak a, b; Figure S14B). No activity was observed in control assays lacking NADPH or using heat-inactivated proteins. To determine the identity of **11** and **12**, these compounds were isolated in greater quantities from root methanolic extracts and structural determination was carried out by NMR (Figure S45–S48 for compound **11** and Figure S49– S52 for compound **12**). The MS/MS spectra of the isolated compounds matched those obtained from the corresponding products **11** and **12** formed when **7** or *d*_4_-**7** was incubated with root protein extract. An additional peak observed on incubation with **7** and *d*_4_-**7** had the same retention time as **12**, but was found to be hydroxylated on the other aromatic ring (**12a**) (Figure S14). Since **12a** did not occur in the methanolic plant extract and could not be further converted into corresponding PP structures, we suspect it to be an artifact obtained only by feeding potential intermediates to root extracts.

**Figure 2.**
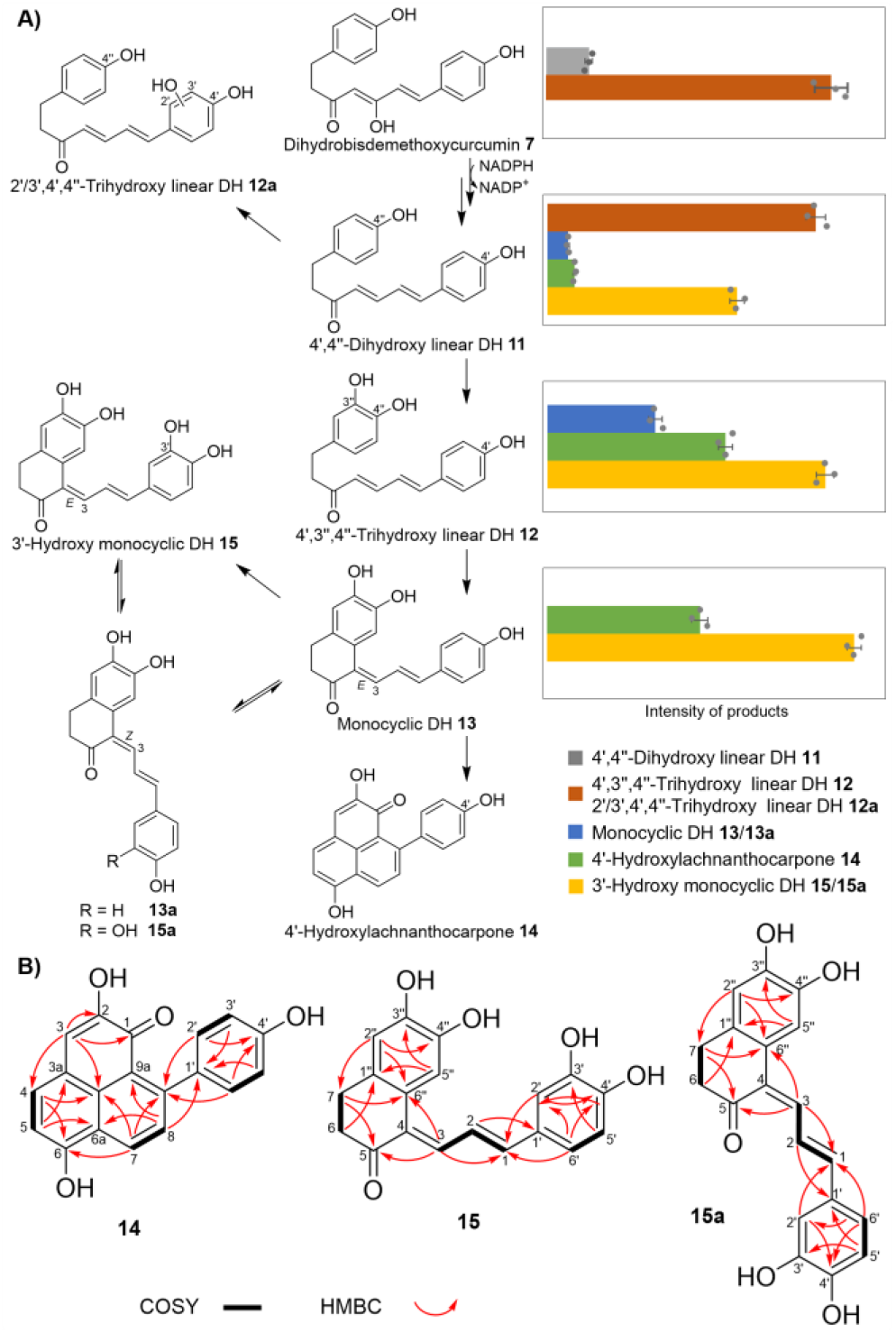
Metabolic conversions catalyzed by protein extracts from *M. lasiocarpa* roots. The substrates dihydrobisdemethoxycurcumin **7**, 4’,4’’-dihydroxy linear DH **11**, 4’,3’’,4’’-trihydroxy linear DH **12**, and monocyclic DH **13** were individually incubated with the crude root protein extract. (A) Depicted are the LCMS peak areas of products detected in incubations with root protein extracts containing the compound immediately to the left. Data are mean ± s.e.m. (n = 3). All products formed are presented in a unified biosynthetic scheme on the left. (B) Key COSY and HMBC correlations for the previously undescribed structures of 4’-hydroxylachnanthocarpone **14** and 3’-hydroxy monocyclic DH **15**/**15a**.

To detect further downstream conversion steps, the isolated compound **12** was incubated with root extract giving rise to three unknown products with ion masses *m/z* 309.1121, 305.0808, and 325.1071 [M + H]^+^ (peak a–c; Figure S16). NMR analysis revealed two monocyclic diarylheptanoid derivatives **13** and the 3’-hydroxy **15** (Figure S57–S64 for compound **13**, Figure S65–S76 for compound **15**, and Table S3) and the PP structure 4’-hydroxylachnanthocarpone **14** (Figure S53–S56 and Table S2). Neither **14** nor **15** have been previously characterized.

The stereochemistry of **13** and **15** align well for the formation of the PP ring system (Figure S18). The idea that monocyclic DHs may be biosynthetic intermediates between linear DHs and PPs was previously raised after a compound related to **13** was isolated from the rhizome of *F. oxysporum*-infected *Musa acuminata* plants.^[23]^ Although no biochemical evidence was provided at the time, the authors were able to chemically transform the compound into a PP structure by demethylation and subsequent oxidation (Figure S19).

To confirm the final cyclization step in PP biosynthesis, isolated **13** and **15** were incubated separately with the crude *M. lasiocarpa* root protein extract. LCMS analysis of assays containing **13** revealed an increase in two peaks at *m/z* 305.0808 and 325.1071 with MS/MS signatures corresponding to those of 4’-hydroxylachnanthocarpone **14** and 3’-hydroxy monocyclic DH **15** (peak a, b; Figure S20). In contrast, no significant increase of the PP product peaks was detected in incubations with **15**. A possible explanation could be the altered reactivity of **15** caused by the additional hydroxy group at the D-ring. Furthermore, it should be noted that **15** is present in the root extract as a trace metabolite, only amounting to approximately 5% of the abundance of **13** (Figure S21).

The identified two-step cyclization of **12** likely proceeds via a sequential 1,4-/1,6-intramolecular Michael addition, initiated by the oxidation of the linear DH substrate to an activated *ortho*-quinone (Figure S22). Given the importance of the keto-enol tautomerized position 6 in the proposed reaction mechanism, it was expected that PP structures without a 6-OH group, which are commonly found in the root extract (Figure S13), are formed in the final step of the pathway. Interestingly, neither the incubation of **14** nor of any of the upstream intermediates with the crude enzyme extract yielded the expected product, 4’-hydroxyanigorufone **R-2** (Figure S13). This finding may be due to either the limitations of the crude protein extract or the existence of an alternate biosynthetic route to anigorufone-type PPs. The NMR data of **13** and **15** showed that they exist as equilibrium isomeric mixtures, with the Δ^3^ double bond in either an *E* (**13** and **15**) or a *Z* configuration (**13a** and **15a**, Table S3). The *Z* isomers **13a** and **15a** are likely intermediates in the formation of a group of diarylheptanoids with a rare bicyclic tetrahydropyran motif. Such diarylheptanoids are known constituents of *M. lasiocarpa* and *Musa* × *paradisiaca* (musellarins A–E; Figure S18).^[22]^

Based on the combined results of the enzymatic assays performed with heterologously expressed enzymes and the administration of intermediates to root protein extracts, a complete reaction pathway can be constructed for the formation of 4’-hydroxylachnanthocarpone **14** in *M. lasiocarpa* starting from *p*-coumaroyl-CoA **1**, which parallels the co-occurring pathway from feruloyl-CoA **2** to linear DH derivatives (Scheme 2).

**Scheme 2.**
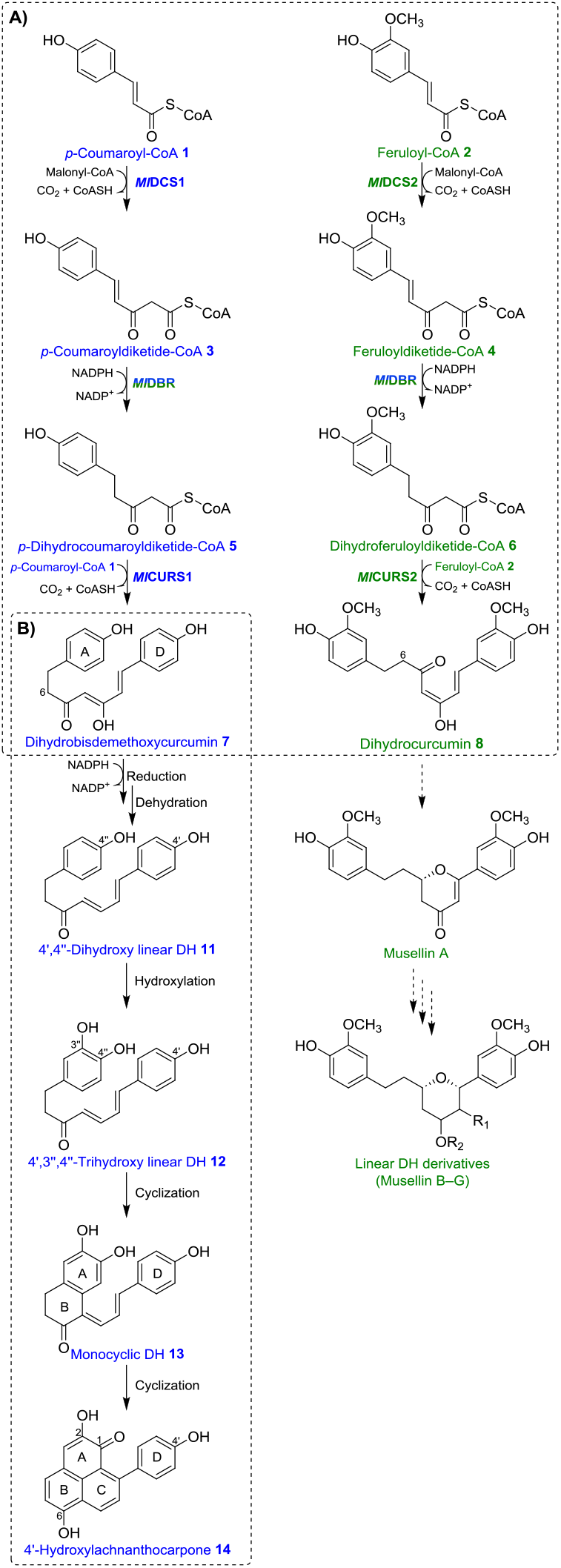
Summary of characterized conversions in *M. lasiocarpa*. (A) Identified enzymes responsible for the parallel synthesis of dihydrobisdemethoxycurcumin **7** and dihydrocurcumin **8**. (B) Downstream biosynthetic transformations observed after incubation with root protein extracts.

The present study provides new molecular and biochemical evidence for the biosynthesis of PPs and linear DH derivatives in *M. lasiocarpa*. Based on the divergent substrate specificities of four type III polyketide synthases and a novel diketide-CoA-accepting medium-chain dehydrogenase/reductase, two distinct assembly lines are present for the production of the dihydrocurcumin-type precursors of either PPs or linear DH derivatives. The stepwise formation of the PP 4’-hydroxylachnanthocarpone constitutes the first complete pathway to a PP scaffold that appears to proceed via two sequential intramolecular Michael additions. The inability of the *M. lasiocarpa* root protein extract to facilitate the formation of anigorufone-type PPs characteristic of this species suggests that there is at least one alternative cyclization route that may not involve monocyclic DH intermediates. Our findings indicate that the biosynthetic network underlying PP metabolism in the Musaceae is more complex than previously expected, and that further studies will be required to fully elucidate the biosynthetic routes to the different PP structures. In the future, it may be possible to exploit this information to introduce disease-resistance to susceptible banana cultivars.

## Supporting information

SI for manuscript

## Supporting Information

The authors have cited additional references within the Supporting Information.^[24–31]^

## Acknowledgements

We gratefully acknowledge the MPI-CE greenhouse team for taking care of the plants; Dr. Veit Grabe for taking pictures of the seeds; Prof. Dr. Yu Chen’s team for assistance with seed collection and RNA sequencing; Dr. Heiko Vogel for assistance with transcriptome analysis; and Dr. Benke Hong and Dr. Ryan Alam for helpful discussions. Seed collection and RNA sequencing were supported by the National Natural Science Foundation of China (32070360). This work was supported by the Max Planck Society.

