## Supplementary material for "Phenylphenalenones and Linear Diarylheptanoid Derivatives are Biosynthesized via Parallel Routes in *Musella lasiocarpa*, the Chinese Dwarf Banana": SI for manuscript

### Table of Contents


Experimental Section 3

Synthesis of Compounds 9

Supplementary Figures 11

NMR Data of Dihydroferuloyl-β-keto acid **6a** 32

NMR Data of Dihydrobisdemethoxycurcumin **7** 37

NMR Data of 4-Hydroxybenzylalcohol-$d_4$ 41

NMR Data of 4-Hydroxybenzaldehyde-$d_4$ 44

NMR Data of Dihydrobisdemethoxycurcumin-$d_4$ $d_4$-**7** 47

NMR Data of Dihydrocurcumin **8** 50

NMR Data of 4',4''-Dihydroxy linear DH **11** 54

NMR Data of 4',3'',4''-Trihydroxy linear DH **12** 58

NMR Data of 4'-Hydroxylachnanthocarpone **14** 62

NMR Data of Monocyclic DH **13** and **13a** 66

NMR Data of 3'-Hydroxy monocyclic DH **15** and **15a** 74

Supplementary Tables 86

References 94

#### Experimental Section

##### Chemicals and Reagents

All solvents used for extractions, chemical synthesis, as well as preparative and analytical HPLC were purchased from VWR in HPLC or HPLC-MS grade quality. Cinnamoyl-CoA, caffeoyl-CoA, *p*-dihydrocoumaroyl-CoA, *p*-coumaroyl-CoA **1**, feruloyl-CoA **2**, and malonyl-CoA were purchased from TransMIT. 3,4-Dihydroxybenzalacetone, raspberry ketone, *p*-hydroxybenzalacetone, and benzalacetone were purchased from Fisher Scientific. Vanillylidenacetone, bisdemethoxycurcumin **9**, and curcumin **10** were purchased from Sigma-Aldrich. Dihydrocurcumin **8** was purchased from Toronto Research Chemicals. Synthesis of dihydrobisdemethoxycurcumin **7** and $d_4$-dihydrobisdemethoxycurcumin $d_4$-**7** is described in **Synthesis of compounds** section.

##### Molecular biology kits

All primers were purchased from Eurofins or Integrated DNA Technologies. PCR reactions were carried out using either Phusion High-Fidelity DNA Polymerase (Thermo Fisher) or Phusion U Hot Start DNA Polymerase (Thermo Fisher), if encountering uracil stalling. PCR products were purified using the innuPREP DOUBLEpure Kit (Analytik Jena). All restriction enzymes were purchased from New England Biolabs. Ligation was achieved using T4 DNA ligase (Thermo Fisher). Plasmid purifications were performed using NucleoSpin Plasmid (Macherey-Nagel). Unless indicated otherwise, all reactions using commercial products were performed according to the guidelines provided by the manufacturer.

##### Plant materials

Seeds of *M. lasiocarpa* were collected at three different developmental stages (yellow, brown, and mature black) from Chuxiong City, Yunnan Province, China. One portion was freeze-dried for metabolite analysis as previously described.[13] Another portion was snap frozen and shipped on dry ice to Novogene for mRNA isolation and RNA-seq analysis.

The outer layer of mature roots of *M. lasiocarpa* plants purchased from Baumschule Eggert, Baumschulenweg, Germany, was used for crude enzyme extraction and metabolite isolation. *M. lasiocarpa* plants were uprooted and the outer layer of mature roots was peeled, snap frozen in liquid nitrogen, and stored at −80 °C until work-up.

For transient gene expression, *Nicotiana benthamiana* plants were grown in the greenhouse of the Max Planck Institute for Chemical Ecology in standard soil under controlled conditions of 22 °C, 60% relative humidity, and a 16-h-light/8-h-dark photoperiod. The leaves of 4–5-week-old plants were used for *Agrobacterium* infiltration.

##### RNA isolation and sequencing

Total RNA was extracted from three different developmental stages of *M. lasiocarpa* seeds (2 biological replicates each) using Trizol reagent kit (Invitrogen) according to the manufacturer’s protocol. The quality and quantity of the RNA obtained were evaluated on an Agilent 2100 Bioanalyzer. After purification of poly(A) mRNA from 1.5 µg total RNA samples using oligo (dT)-coupled beads, the mRNA was fragmented into small pieces in fragmentation buffer. Construction of paired-end cDNA libraries was carried out using the NEBNext Ultra RNA Library Prep Kit for Illumina (New England Biolabs). The average insert size for the paired-end libraries was 300 ± 50 bp. Finally, 2 × 150 bp paired-end sequencing was performed on an Illumina Hiseq platform. Between 23 and 27 million RNA-Seq reads were collected for each sample. FASTQC was used for quality control.

***De novo* transcriptome assembly, annotation and evaluation.**

We applied read quality filtering with trimmomatic software[24] using the following options (LEADING:10 TRAILING:10 SLIDINGWINDOW:4:20 MINLEN:40–normalize_reads). The trimmed reads from all samples were pooled and assembled using CLC Genomics Workbench v8.1. The assembly parameters included automatic word size = yes; bubble size = 250; minimum contig length = 200; auto-detect paired distances = yes; perform scaffolding = yes; map reads back to contigs = yes, with mismatch cost = 2; insertion and deletion cost = 3, length fraction = 0.8, and similarity fraction = 0.9. Scaffolding was chosen, and conflicts among individual bases were resolved in all assemblies by favoring the base with the highest frequency. Contigs shorter than 200 bp were excluded from the final datasets. In case some candidate genes were not assembled in full-length, we also performed *de novo* assembly of subsets of the samples using rnaSPAdes[25] to get mRNA sequences in full-length.

Following the *de novo* assembly, we conducted blastx[26] searches against the NCBI-NR database and used Blast2GO[27] for Gene Ontology (GO) annotation. For expression level analyses, CLC Genomics Workbench v8.1 was used to generate BAM mapping files using previously described parameters for read mapping and normalization.[28] Mapped reads were log2-transformed and normalized using the quantile method. Statistical analysis of the normalized data was performed using the "empirical analysis of digital gene expression" (EDGE) tool, implemented in CLC Genomics Workbench. For both methods, the criteria for differentially expressed genes were a minimum two-fold change in expression and a false discovery rate (FDR)-corrected p-value of <0.05.

**LC-HRESIMS analysis**

For the HPLC-HRESIMS measurement, an Agilent Infinity 1260 HPLC system consisting of a quaternary pump (G1311B), autosampler (G1367E), column oven (G1316A), and diode array detector (G1315D) was coupled to a Bruker Compact OTOF mass spectrometer. The instrumentation was controlled by Bruker Hystar version 3.2/Bruker OTOFControl version 4.0. Ionization was performed by electrospray ionization (ESI) in positive mode with a capillary voltage of 4500 V and an end plate offset of 500 V; a nebulizer pressure of 1.8 bar was used, with nitrogen at 220 °C and a flow of 9 L/min as the drying gas. The acquisition was performed in profile mode with a full scan *m/z* range of 50–1300. The collision energy of the MSMS with multiple reaction monitoring (MRM) mode was set at 10, 15, 20, and 35 eV. A sodium formate-isopropanol solution was injected at the beginning of each run and the *m/z* values were recalibrated using the expected cluster ion *m/z* values. DataAnalysis version 4.3 (Bruker) was used to analyze the LCMS data.

Method 1

HPLC conditions were as follows: An Agilent Zorbax C18 column (3.5 µm; 150 × 4.6 mm) was used with the binary gradient conditions (A: $H_2O$, B: $CH_3CN$, both containing 0.1% formic acid) 0–3 min 5%

B, 3–15 min 5–100% B, and 15–20 min 100% B, at a constant flow rate of 400 µL/min. This method was used to analyze the enzymatic products of *Ml*DCS, *Ml*DBR, and *Ml*CURS.

Method 2

HPLC conditions were as follows: An Agilent EC-C18 column (2.7 µm; 100 × 4.6 mm) was used with the binary gradient conditions (A: $H_2O$, B: $CH_3CN$, both containing 0.1% formic acid) 0–5 min 5% B, 5–35 min 5–100% B, and 35–40 min 100% B, at a constant flow rate of 500 µL/min. This method was used to analyze the enzymatic products of crude protein assays and the metabolites of the outer root layer.

**Candidate gene cloning**

For heterologous expression in *Escherichia coli,* all genes described in this study (Supplementary Table S4) were amplified from cDNA of *M. lasiocarpa* seeds using overhang primers designed for pRSET B cloning (Supplementary Table S5). The resulting PCR products were introduced into the pRSET B expression vector by standard restriction-ligation cloning. One shot TOP10 chemically competent *E. coli* cells (Invitrogen) were transformed with the constructs and selected on LB plates containing ampicillin (100 µg/mL). Sanger sequencing was used to confirm the correct insertion of the genes.

For transient expression in *N. benthamiana*, the candidate genes were cloned into the binary pCambia 2300u vector by adapting the USER cloning strategy. Following the amplification of genes using the uracil-containing primers listed in Supplementary Table S5, USER reactions were carried out as previously described. Reaction mixtures were directly transformed into 10-beta competent *E. coli* cells (New England Biolabs) and recombinant colonies were selected on LB agar plates containing kanamycin (50 µg/mL). Sanger sequencing was used to confirm the correct insertion of the genes.

**Heterologous expression in *E. coli* and protein purification**

Sequence-confirmed pRSET B constructs were transformed into BL21 Star (DE3) pLysS one shot chemically competent *E. coli* cells (Invitrogen) and positive transformants were selected on LB plates supplemented with ampicillin (100 µg/mL) and chloramphenicol (34 µg/mL). Single colonies were inoculated in 5 mL LB medium (100 µg/mL ampicillin and 34 µg/mL chloramphenicol) and incubated over night at 37 °C and constant agitation. On the next day, 2 mL preculture were used to inoculate main cultures containing 100 mL LB medium (100 µg/mL ampicillin). Cultures were grown at 37 °C until reaching $OD_{600}$ = 0.6–0.8. Heterologous gene expression was induced by adding 500 µM isopropyl β-D-1-thiogalactopyranoside (IPTG) and incubation at 18 °C for 16–18 hours. Cells were harvested by centrifugation (5 min at 5000 *g*) and resuspended in 10 mL binding buffer (50 mM tris-HCl pH 8, 50 mM glycine, 500 mM NaCl, 20 mM imidazole, 5% v/v glycerol) supplemented with 0.2 mg/mL lysozyme and complete EDTA-free protease inhibitor cocktail (Roche). All protein purification steps were performed on ice with cooled buffers and centrifuges. After 30 min incubation on ice, the cells were lysed by sonication using a Sonics Vibra Cell at 40% amplitude, 2s ON, 3s OFF, 5 min in total. Crude lysates were centrifuged at 15000 *g* for 15 min, and the clarified lysates were incubated with 250 µL Ni-NTA agarose beads (Qiagen) for 60 min. The beads were then pelleted (1000 *g*, 30 s) and washed 3 times with 10 mL binding buffer. After removal of the supernatant, protein elution was performed by resuspending the beads in 600 µL elution buffer (50 mM tris-HCl pH 8, 50 mM glycine, 500 mM NaCl, 250 mM imidazole, 5% v/v glycerol). Dialysis and buffer exchange were performed with final buffer (100 mM potassium phosphate buffer pH 7.5, 1 mM dithiothreitol) in centrifugal concentrators with 10 kDa size exclusion. Proteins were aliquoted in 40 µL, snap frozen and stored at −80 °C for *in vitro* enzyme assays.

***In vitro* enzyme assays**

Enzymes and various combinations were incubated in enzyme assays comprising different starter substrates, extender substrates, and cofactors as listed in Supplementary Table S6. All reaction mixtures were added up to 100 µL with potassium phosphate buffer pH 7.5 (100 mM) and incubated at 37 °C for 2 hours. Co-incubations of *Ml*DCS2 and *Ml*DBR were also tested at 1, 1.5, 3, and 10 hours. Negative controls contained boiled enzymes (90 °C, 10 min) in the reaction mixtures. All reactions were quenched by addition of 1 volume of MeOH, filtered through 0.22-µm PTFE filters and analyzed by the LC-HRESIMS (Method 1). Identification of products was performed by comparison of retention times and MS/MS spectra with either commercial or structure-elucidated standards.

**Preparative scale *in vitro* reaction and isolation of dihydroferuloyl-β-keto acid 6a**

Reaction mixtures in a total volume of 9 mL consisted of *Ml*DCS2 (1 µM), *Ml*DBR (1 µM), feruloyl-CoA **2** (100 µM), malonyl-CoA (100 µM), NADPH (1 mM), and potassium phosphate pH 7.5 (100 mM). After incubation at 37 °C for 10 hours, the reaction was passed through a preconditioned HR-X solid phase extraction (SPE) cartridge (30 mg, 1 mL, Macherey-Nagel). After washing with acidified $H_2O$, the sample was eluted with 2 mL $CH_3CN$. A dilution (1:1000) of the crude mixture was analyzed by LC-HRESIMS (method 1) and the desired product was confirmed by *m/z* and retention time. The pooled mixture was evaporated with nitrogen gas. The HPLC system was then decoupled from the MS for sample purification (method 1) and the target peak was manually collected at the same retention time. The isolated compound was then subjected to NMR analysis. Since dihydroferuloyl-β-keto acid **6a** is unstable in $CH_3CN$, the entire isolation procedure had to be completed in one day.

**Transient expression of candidate genes in *N. benthamiana***

Sequence-confirmed pCambia 2300u vectors carrying candidate genes were transformed into *Agrobacterium tumefaciens* GV3101 cells. Recombinant colonies were selected on LB agar containing kanamycin (50 µg/mL), gentamycin (25 µg/mL), and rifampicin (10 µg/mL). Single colonies were picked and grown overnight in 10 mL of LB (50 µg/mL kanamycin, 25 µg/mL gentamycin, and 10 µg/mL rifampicin) at 28 °C. The next day, 5 mL of the precultures were used to inoculate 50 mL of LB medium (10 mL was used to inoculate 100 mL in the case of the p19-expressing helper strain) with the identical antibiotics for another overnight growth. On the following day, the cultures were centrifuged (3000 *g*, 10 min) and the supernatant was removed. The cell pellet was resuspended in infiltration buffer (10 mM MES, 10 mM $MgCl_2$, 100 µM acetosyringone, pH 5.6) to an optical density $OD_{600}$ of 0.5. After incubation for 2 h at room temperature, all tested strains were mixed 1:1 with a helper strain expressing P19, which is commonly used to suppress gene silencing.[29] For co-infiltration of multiple constructs, the suspensions were mixed in equal proportions. Finally, 4–5-week-old *N. benthamiana* plants were infected by syringe-infiltration into the abaxial side of the two youngest fully expanded leaves. Each experiment was performed in three biological replicates. Five days after infiltration, leaves were harvested for LC-HRESIMS analysis (Method 1).

##### Sample harvest and analysis

Harvested snap-frozen *N. benthamiana* leaf tissue (100 mg) was homogenized on a TissueLyser II (Qiagen) using 2 mm diameter tungsten beads with vigorous shaking at 22 Hz for 2 min. MeOH (350 µL) was added to each sample before vigorous vortexing for 1 min. Samples were centrifuged at high speed (>13000 *g*, 15 min) and filtered through 0.22 µm PTFE syringe filters. For alkaline lysis, equal volumes of $H_2O$ were added followed by 10 µL of 10 M NaOH and samples were incubated at 65 °C for 10 min. To neutralize the solution, 17 µL of 6 M HCl was introduced. 1 volume of ethyl acetate was used for extraction. The solvent was then evaporated under nitrogen gas, and the residue was reconstituted with 350 µL MeOH for injection into LC-HRESIMS (Method 1). Other filtered samples were directly injected into the LC-HRESIMS (Method 1). Identification of metabolites was accomplished by comparing retention times and MS/MS spectra with those of authentic standards.

##### Preparation of crude protein extracts from *M. lasiocarpa* roots

To isolate crude protein extracts, the outer layer of mature roots was homogenized using a mortar and pestle with liquid nitrogen and a small scoop of polyvinylpolypyrrolidone (PVPP). The powdered tissue was extracted with ice-cold extraction buffer [50 mM Tris-HCl pH 7.4, 50 mM glycine, 5% glycerol, 0.5 M NaCl, 1 mM phenylmethylsulfonyl fluoride, EDTA free protease inhibitor cocktail (1 tablet in 50 mL)] per 1 g of powered tissue. The mixture was incubated in a cold chamber (4 °C) for 30 min with periodic gentle inversions. The extracted homogenate was pelleted at 3500g for 5 min at 4 °C to remove plant cell debris, and the supernatant was filtered through Miracloth (Merck-Millipore) and collected in a pre-cooled tube. The filtered extract was centrifuged again using the same parameters. The supernatant was removed and aliquoted into prechilled microfuge tubes, which were then snap frozen in liquid nitrogen and stored at −80 °C.

##### *In vitro* assays with crude protein extract of *M. lasiocarpa* roots

Assays with synthetic compounds: dihydrobisdemethoxycurcumin **7** and $d_4$-dihydrobisdemethoxycurcumin $d_4$-**7**

For *in vitro* experiments, a 100 µL volume assay contained 100 µM dihydrobisdemethoxycurcumin **7** or $d_4$-dihydrobisdemethoxycurcumin $d_4$-**7** and 1 mM NADPH, and the remaining volume was crude protein extract. Negative controls consisted of 1) boiled crude protein extract (90 °C, 10 min) with substrates and NADPH, and 2) crude protein extract with substrates but without NADPH in the reaction mixtures.

Assays with isolated metabolites: 4',4''-dihydroxy linear DH **11**, 4',3'',4''-trihydroxy linear DH **12**, monocyclic DH **13**, and 3'-hydroxy monocyclic DH **15**

For *in vitro* experiments, a 100 µL volume assay contained 100 µM 4',4''-dihydroxy linear DH **11** or 4',3'',4''-trihydroxy linear DH **12** or monocyclic DH **13** or 3'-hydroxy monocyclic DH **15** and the remaining volume was crude protein extract. Negative controls consisted of 1) boiled crude protein extract (90 °C, 10 min) with substrates and 2) crude protein extract without substrates (to account for background activity) in the reaction mixtures.

All reaction mixtures were incubated at 37 °C for 2 hours. Reactions were quenched by adding 1 volume of MeOH, filtered through 0.22-µm PTFE filters and analyzed by LC-HRESIMS (Method 2).

**Extraction and purification of 4',4''-dihydroxy linear DH 11, 4',3'',4''-trihydroxy linear DH 12, monocyclic DH 13, 4'-hydroxylachnanthocarpone 14, and 3'-hydroxy monocyclic DH 15 from *M. lasiocarpa* roots**

The 4 g freeze-dried outer layer of mature roots was ground to powder using an IKA M20 universal mill, suspended in 50 mL MeOH, and then shaken on a rotary shaker (180 rpm) for 4 h at room temperature. After filtration, the remaining residue was extracted twice more with MeOH (50 mL). The combined MeOH extract was evaporated with nitrogen gas to yield a residue of 306 mg, which was then reconstituted with MeOH and subjected to separation by preparative HPLC. Preparative HPLC was performed on a Shimadzu Prominence HPLC system consisting of a degasser (DGU-20A5), gradient pump (LC-20AT), autosampler (SIL-20AC), column oven (CTO-20A), UV detector (SPD-20A), fraction collector (FRC-10A), and system controller (CBM-20A). The HPLC was equipped with a Nucleodur C-18 HTec column (5 µm; 250 × 4.6 mm; Macherey-Nagel) and a binary solvent system of $H_2O$ (solvent A) and $CH_3CN$ (solvent B), both containing 0.1% formic acid. The constant flow rate was set at 800 µL/min and a gradient was used: 0–5 min 5% B, 5–30 min 5–45% B, 30–40 min 45–100% B, and 40–45 min 100% B. The eluted fractions were analyzed by LC-HRESIMS (1:1000 dilution) and the desired fractions were pooled and evaporated to dryness. The isolated compounds [4',4''-dihydroxy linear DH **11** (0.4 mg), 4',3'',4''-trihydroxy linear DH **12** (0.5 mg), monocyclic DH **13** (0.3 mg), 4'-hydroxylachnanthocarpone **14** (0.5 mg), and 3'-hydroxy monocyclic DH **15** (0.08 mg)] were subjected to NMR analysis and then used as substrates for *in vitro* assays with crude protein extract. As compound 3'-hydroxy monocyclic DH **15** required additional purification by isocratic elution (27% B) the remaining amount was only sufficient for $^1$H NMR analysis.

**NMR**

NMR spectra ($^1$H NMR with $H_2O$ suppression, COSY, HMBC, HSQC, and ROESY spectra) were measured at 298 K on Bruker Advance III HD 700 and Bruker AV 500 NMR spectrometers. Spectrometer control, data acquisition, and processing were performed using Bruker TopSpin version 3.6.1. Chemical shifts are expressed in δ (ppm) relative to the residual solvent signals of $CD_3OD$ ($\delta_{H/C}$ 3.31/49.15), $CD_3CN$ ($\delta_{H/C}$ 1.94/1.39), or acetone-$d_6$ ($\delta_{H/C}$ 2.05/29.92).

#### Synthesis of Compounds

##### Synthesis of dihydrobisdemethoxycurcumin 7

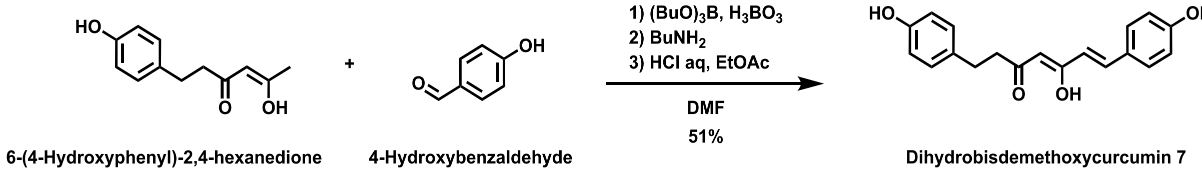

Dihydrobisdemethoxycurcumin **7** ((1*E*)-1,7-bis(4-hydroxyphenyl)-1-heptene-3,5-dione) was synthesized according to a known protocol[30] with minor modifications. 6-(4-Hydroxyphenyl)-2,4-hexanedione was synthesized from 4-hydroxybenzaldehyde (Thermo scientific) as previously described.[31] Under argon atmosphere, a mixture of 6-(4-hydroxyphenyl)-2,4-hexanedione (19.7 mg, 0.096 mmol) and $H_3BO_4$ (8.9 mg, 0.14 mmol, Riedel-de Haën) in anhydrous DMF (0.4 mL) was stirred at 85 °C for 30 min, followed by the addition of 4-hydroxybenzaldehyde (6 mg, 0.05 mmol) and $(BuO)_3B$ (26 µL, 0.10 mmol, Thermo scientific) at room temperature. *n*-$BuNH_2$ (0.007 mmol, Sigma-Aldrich) as anhydrous DMF solution (70 µL, 0.1 mmol/mL) was added dropwise at 85 °C. After stirring at 85−90 °C for 2 hours, aqueous HCl (1M, 0.75 mL) and EtOAc (0.75 mL) were added at room temperature and the mixture was stirred at 60 °C for 30 min. The reaction mixture was extracted twice with EtOAc. The organic phase was washed with brine, dried over anhydrous $Na_2SO_4$ and concentrated *in vacuo*. The residue was purified by flash silica gel column chromatography (*n*-hexane/EtOAc from 97:3-65:35) to give dihydrobisdemethoxycurcumin **7** (7.7 mg, 0.025 mmol, 50%). NMR data are available in Figure S28–S31.

##### Synthesis of dihydrobisdemethoxycurcumin-$d_4$ $d_4$-7

Synthesis of 4-hydroxybenzylalcohol-$d_4$

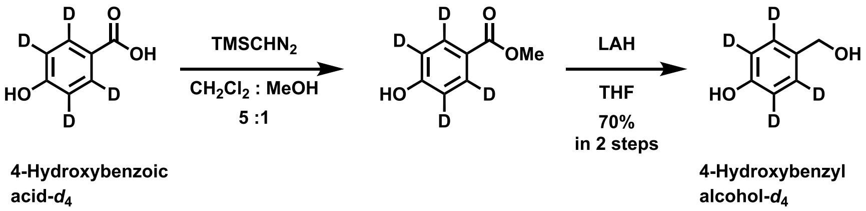

$TMSCHN_2$ (158 µL, 0.32 mmol, 2.0 M in diethyl ether, Aldrich) was added to 4-hydroxybenzoic acid-$d_4$ (30 mg, 0.21 mmol, BOC sciences) in anhydrous $CH_2Cl_2$: anhydrous MeOH (1 mL: 0.2 mL) at 0 °C under argon atmosphere. After stirring for 1 hour, the solvent was evaporated and the resulting ester was used for further reactions without purification.

To a solution of the crude methyl ester in anhydrous THF (1 mL), lithium aluminum hydride solution (0.63 mL, 0.63 mmol, 1.0 M in THF, Aldrich) was added at 0 °C. The reaction mixture was stirred at 0 °C under argon for 30 min, then at room temperature for 40 min. EtOAc and saturated aqueous Rochell salt were carefully added at 0 °C. After stirring for 30 min, the reaction mixture was extracted with EtOAc (x 3). The organic phase was washed with saturated aqueous $NH_4Cl$ and brine, dried over anhydrous $Na_2SO_4$ and concentrated *in vacuo*. The residue was purified by flash silica gel column chromatography (*n*-hexane/EtOAc from 95:5-50:50) to give 4-hydroxybenzylalcohol-$d_4$ (19 mg, 0.148

mmol, 70%): HRMS (ESI-TOF, positive) *m/z*: calcd for $C_7H_3D_4O^{\cdot}$ [M–OH]$^{\cdot}$ 111.0748, found 111.0747. NMR data are available in Figure S32–S34.

Synthesis of 4-hydroxybenzaldehyde-$d_4$

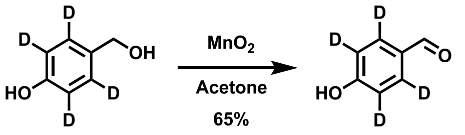


**4-Hydroxybenzyl alcohol-*d*4** **4-Hydroxybenzaldehyde-*d*4**

$MnO_2$ (51 mg, 0.59 mmol, Merck) was added in three portions every 40 min to 4-hydroxybenzylalcohol-$d_4$ (5 mg, 0.039 mmol) in anhydrous acetone at room temperature. After 2 hours of stirring under argon from the first $MnO_2$ addition, the reaction mixture was filtered through short celite pad and concentrated *in vacuo*. The residue was purified by flash silica gel column chromatography (*n*-hexane/EtOAc from 95:5-50:50) to give 4-hydroxybenzaldehyde-$d_4$ (3.3 mg, 0.026 mmol, 65%): HRMS (ESI-TOF, positive) *m/z*: calcd for $C_7H_3D_4O_2$ $[M+H]^+$ 127.0692, found 127.0690. NMR data are available in Figure S35–S37.

Synthesis of dihydrobisdemethoxycurcumin-$d_4$ $d_4$-**7**

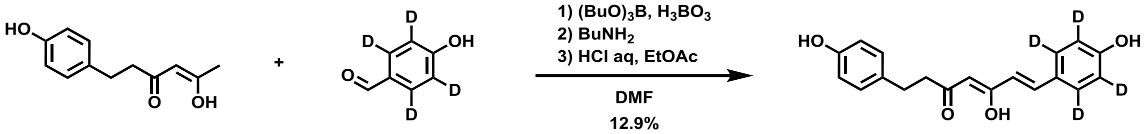


**6-(4-Hydroxyphenyl)-2,4-hexanedione** **4-Hydroxybenzaldehyde-*d*4** **Dihydrobisdemethoxycurcumin *d*4-7**

Dihydrobisdemethoxycurcumin-$d_4$ $d_4$-**7** was synthesized as described for dihydrobisdemethoxycurcumin **7**. Under argon atmosphere, a mixture of 6-(4-hydroxyphenyl)-2,4-hexanedione (26.4 mg, 0.128 mmol) and $H_3BO_4$ (12.0 mg, 0.19 mmol, Riedel-de Haën) in anhydrous DMF (0.3 mL) was stirred at 90 °C for 30 min, followed by addition of 4-hydroxybenzaldehyde-$d_4$ (8.2 mg, 0.065 mmol) in anhydrous DMF (0.3 mL) and $(BuO)_3B$ (35 μL, 0.13 mmol, Thermo scientific) at room temperature. *n*-$BuNH_2$ (0.0097 mmol, Sigma-Aldrich) as anhydrous DMF solution (97 μL, 0.1 mmol/mL) was added dropwise at 90 °C. After stirring at 90 °C for 3 hours, aqueous HCl (1M, 0.75 mL) and EtOAc (0.75 mL) were added at room temperature and the mixture was stirred at 60 °C for 30 min. The reaction mixture was extracted twice with EtOAc. The organic phase was washed with brine, dried over anhydrous $Na_2SO_4$ and concentrated *in vacuo*. The residue was purified by flash silica gel column chromatography (*n*-hexane/EtOAc from 97:3-65:35) to give dihydrobisdemethoxycurcumin-$d_4$ $d_4$-**7** (2.6 mg, 0.0083 mmol, 13%). The chemical data are shown in Figure S38–S40.

#### Supplementary Figures

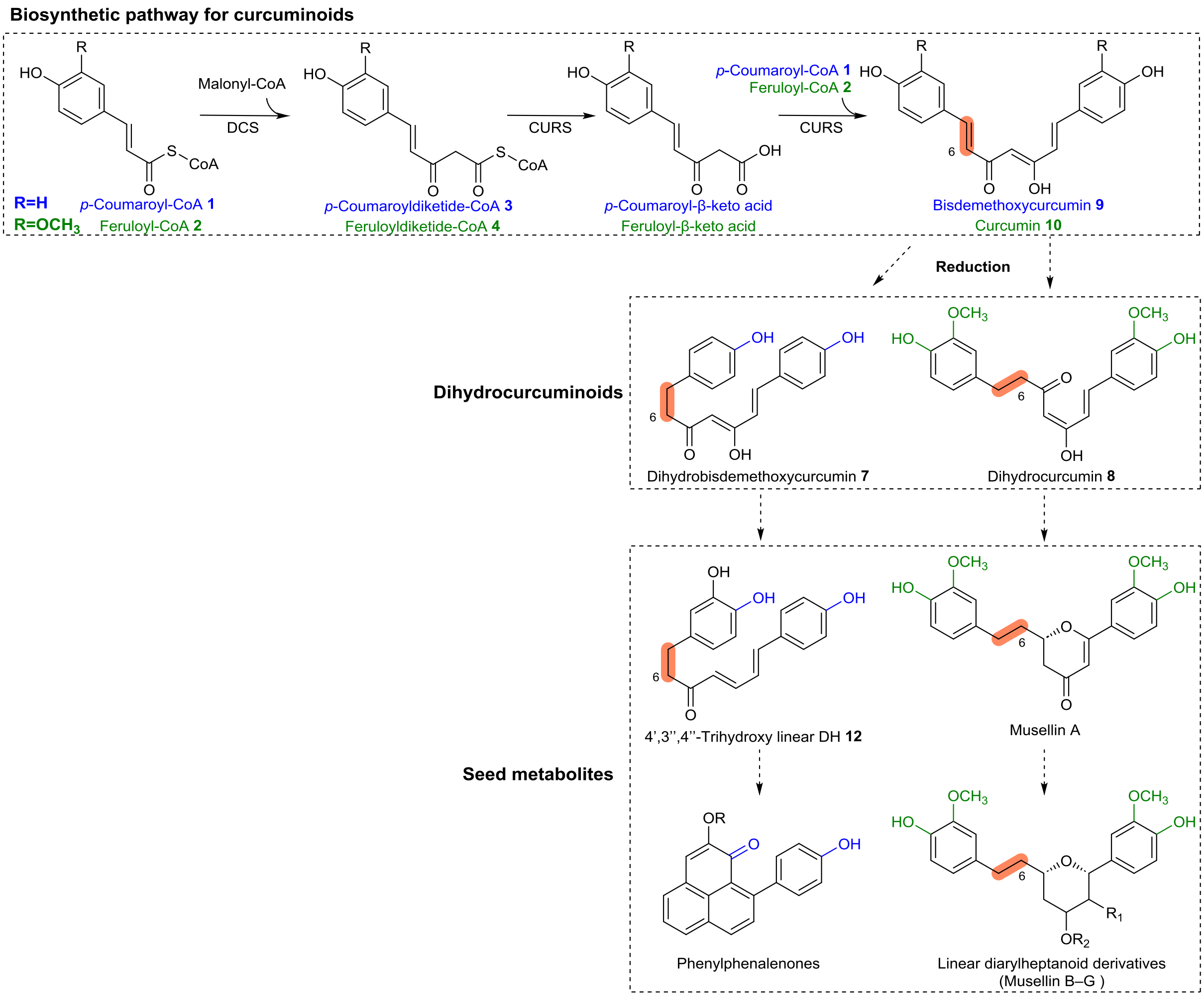


**Figure S1.** The putative biosynthetic pathway for phenylphenalenones and linear diarylheptanoid derivatives found in the seeds of *M. lasiocarpa*.

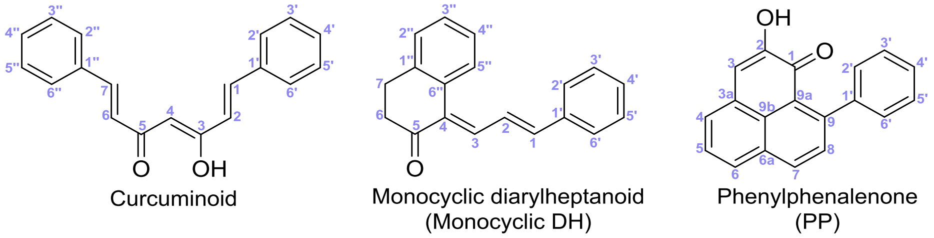


**Figure S2.** Nomenclature. Atomic numbering of the scaffolds of the main compounds in this manuscript.

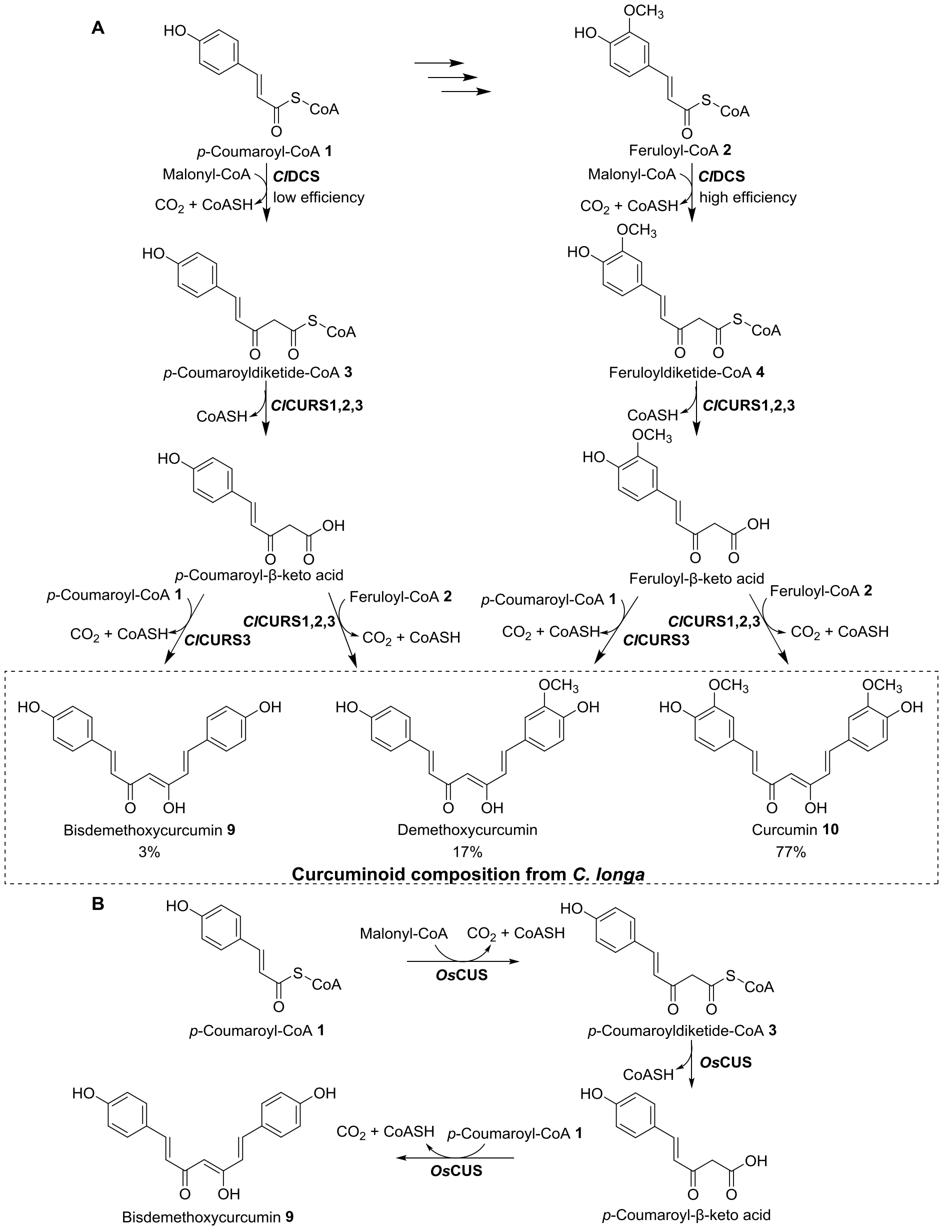


**Figure S3.** Biosynthesis of curcuminoids in *C. longa* and *O. sativa*. (A) Two-step formation of curcuminoids by *Cl*DCS and *Cl*CURS in *C. longa. In vitro* analysis revealed that *Cl*DCS prefers feruloyl-CoA as starter substrate, but could also accept *p*-coumaroyl-CoA with lower efficiency. While *Cl*CURS1 and *Cl*CURS2 prefer feruloyl-CoA as a starter substrate, *Cl*CURS3 can accept both feruloyl-CoA and *p*-coumaroyl-CoA almost equally.[14] (B) One-pot formation of bisdemethoxycurcumin by *Os*CUS in *O. sativa. Os*CUS prefers *p*-coumaroyl-CoA.[15]

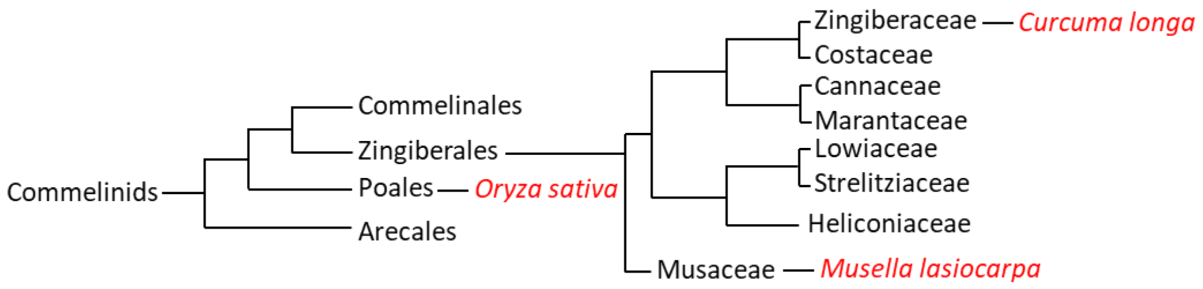


**Figure S4.** Phylogenetic relationship of *M. lasiocarpa*, *C. longa*, and *O. sativa*.[17]

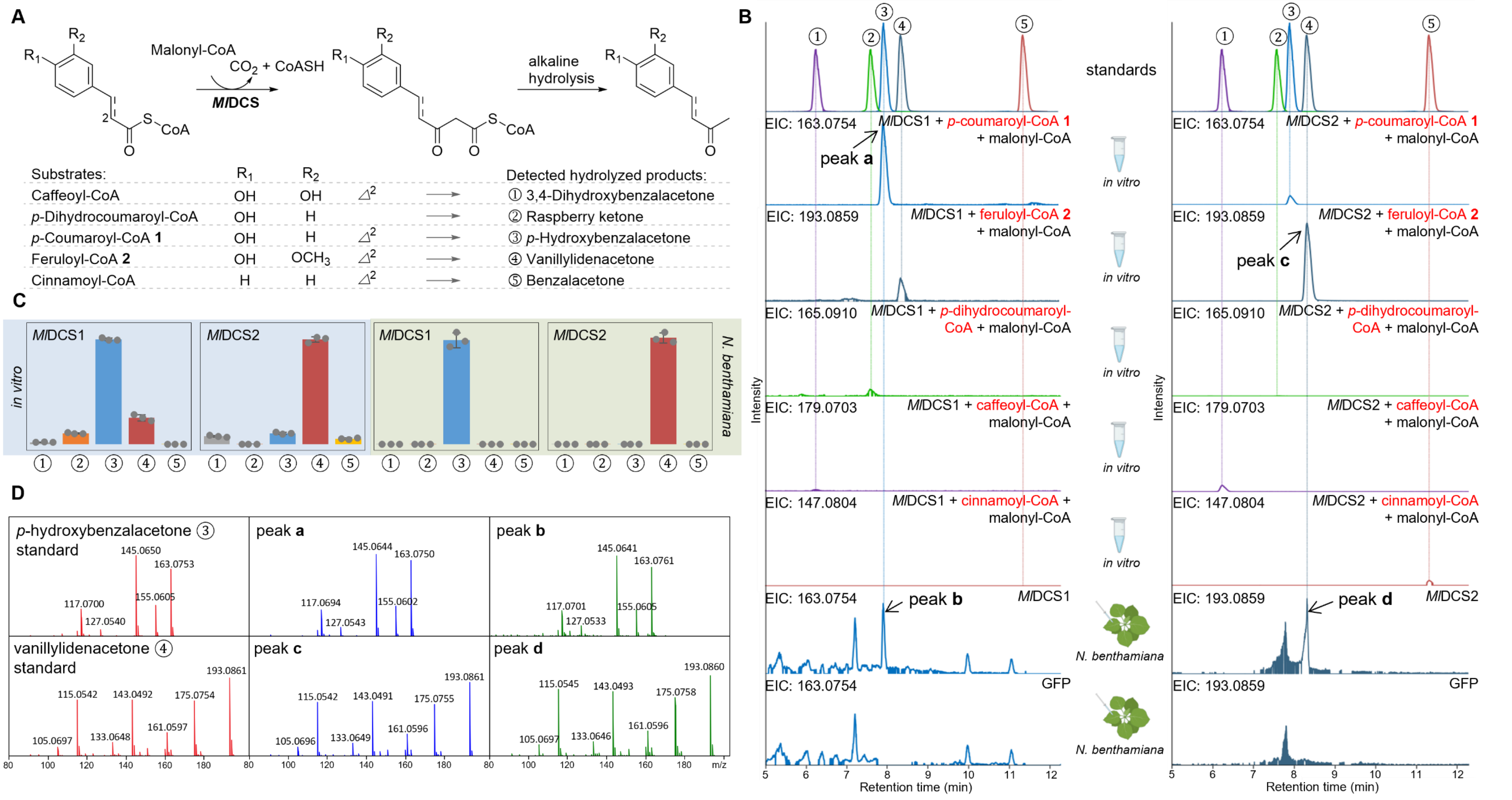


**Figure S5.** Functional characterization and substrate preferences of *Ml*DCS1 and *Ml*DCS2 *in vitro* and in *N. benthamiana*. (A) Scheme of the *Ml*DCS-catalyzed reaction with different starter substrates. (B) Extracted ion chromatograms (EIC) corresponding to the mass of the diketide-CoA hydrolysis products obtained from *in vitro* assays using purified *Ml*DCS1 (left) and *Ml*DCS2 (right) with five different starter substrates. The chromatograms obtained from *N. benthamiana* plants transiently expressing either *Ml*DCS1, *Ml*DCS2, or GFP as a negative control are shown at the bottom. (C) The LCMS peak area of the products obtained from *in vitro* assays and transgenic *N. benthamiana* plants, both containing the indicated enzymes. Data are mean ± s.e.m. (n = 3). (D) MS/MS (20 eV) spectra of *p*-hydroxybenzalacetone ③ produced *in vitro* (peak **a**, blue) and in *N. benthamiana* (peak **b**, green) compared to the standard (red). MS/MS (20 eV) spectra of vanillylidenacetone ④ produced *in vitro* (peak **c**, blue) and in *N. benthamiana* (peak **d**, green) in comparison to the standard (red).

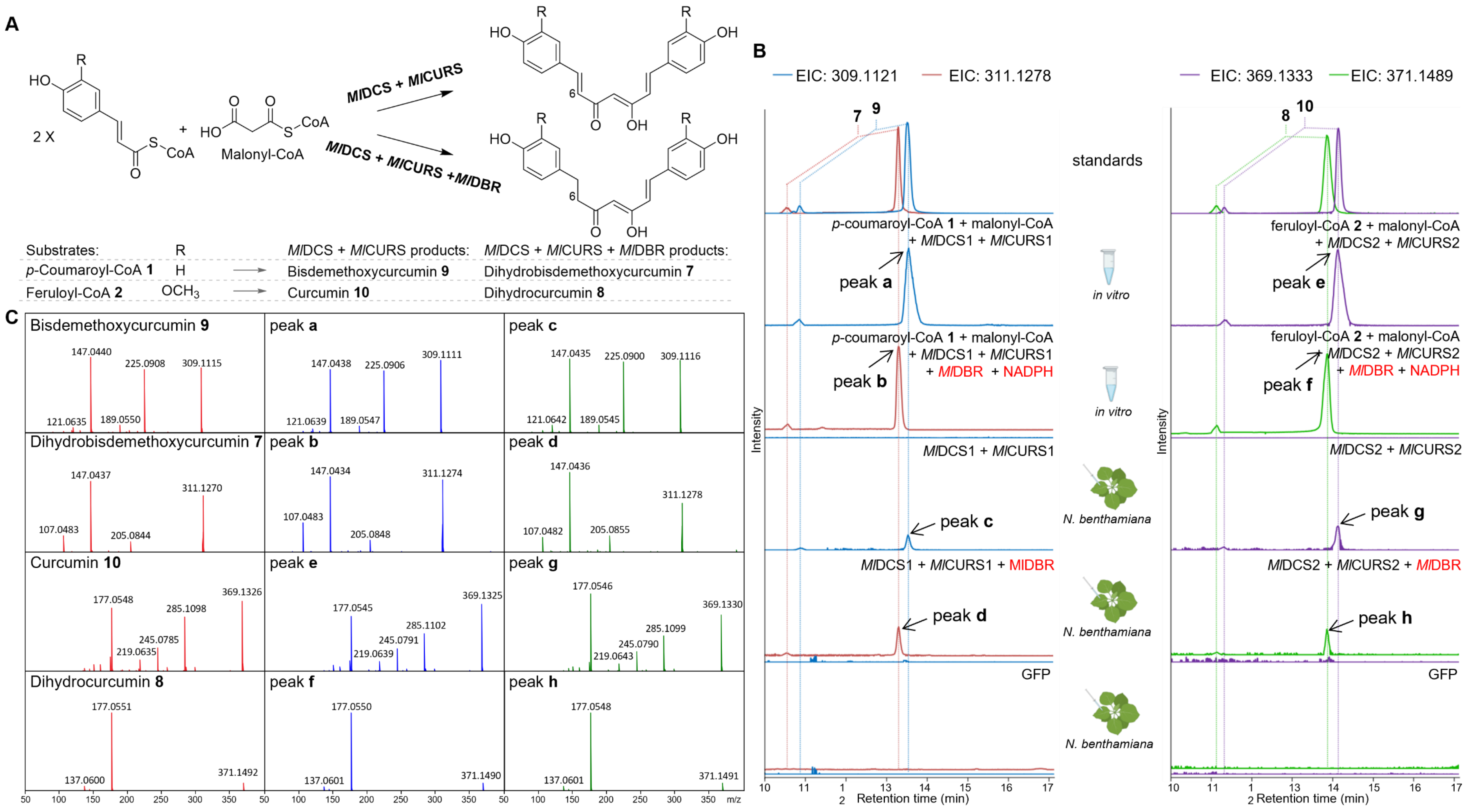


**Figure S6.** Combined activity of *Ml*DCS, *Ml*CURS, and *Ml*DBR. (A) Scheme of the enzymatic reactions catalyzed by the combination of *Ml*DCS/*Ml*CURS and *Ml*DCS/*Ml*CURS/*Ml*DBR. (B) Extracted ion chromatograms (EIC) showing the reaction products of *in vitro* assays using the indicated combinations of purified enzymes with their corresponding preferred substrates. The chromatograms obtained from transiently transformed *N. benthamiana* plants are shown at the bottom. The combination of *Ml*DCS/*Ml*CURS formed curcuminoids, while the introduction of *Ml*DBR led to the formation of dihydrocurcuminoids. All products were observed as two peaks due to the presence of diketo–ketoenol tautomers. The *in vitro* assays and *N. benthamiana* infiltrations were repeated three times with similar results as summarized in **Figure 1** and **Figure S7**, respectively. (C) MS/MS (15 eV) spectra of bisdemethoxycurcumin **9** produced *in vitro* (peak **a**, blue) and in *N. benthamiana* (peak **c**, green) in comparison to the standard (red). MS/MS (15 eV) spectra of dihydrobisdemethoxycurcumin **7** produced *in vitro* (peak **b**, blue) and in *N. benthamiana* (peak **d**, green) in comparison to the standard (red). MS/MS (15 eV) spectra of curcumin **10** produced *in vitro* (peak **e**, blue) and in *N. benthamiana* (peak **g**, green) in comparison to the standard (red). MS/MS (15 eV) spectra of dihydrocurcumin **8** produced *in vitro* (peak **f**, blue) and in *N. benthamiana* (peak **h**, green) in comparison to the standard (red).

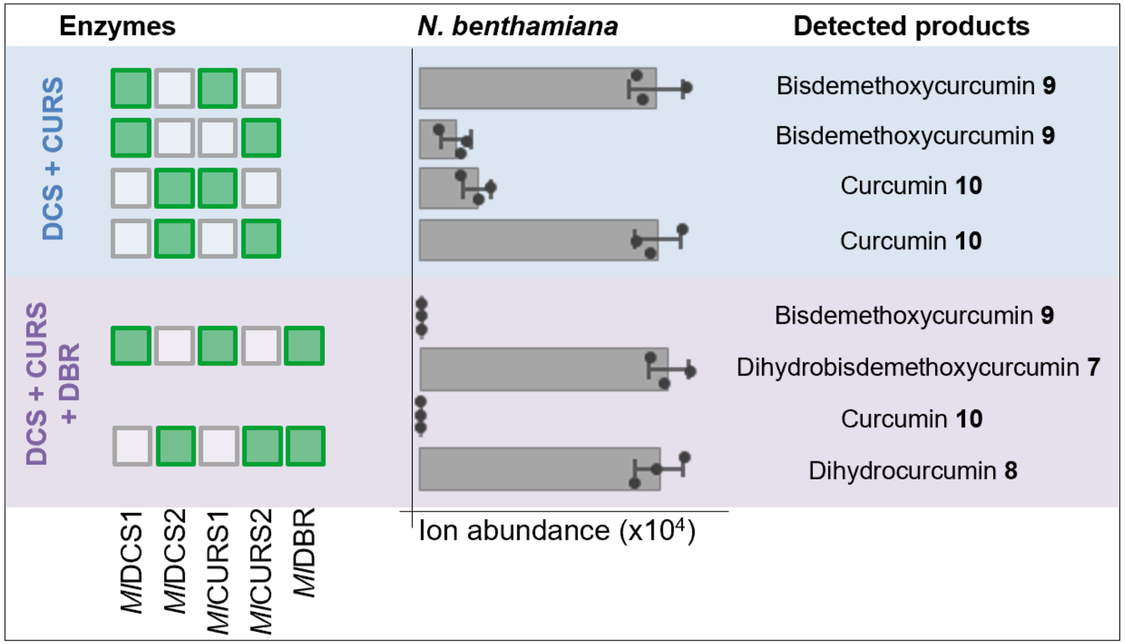


**Figure S7.** The LCMS peak area of products formed in *N. benthamiana* plants transiently transformed with different combinations of *Ml*DCS, *Ml*CURS, and *Ml*DBR. All products were validated by comparison to either purchased or synthetic authentic standards. Data are mean ± s.e.m. (n = 3 biological replicates).

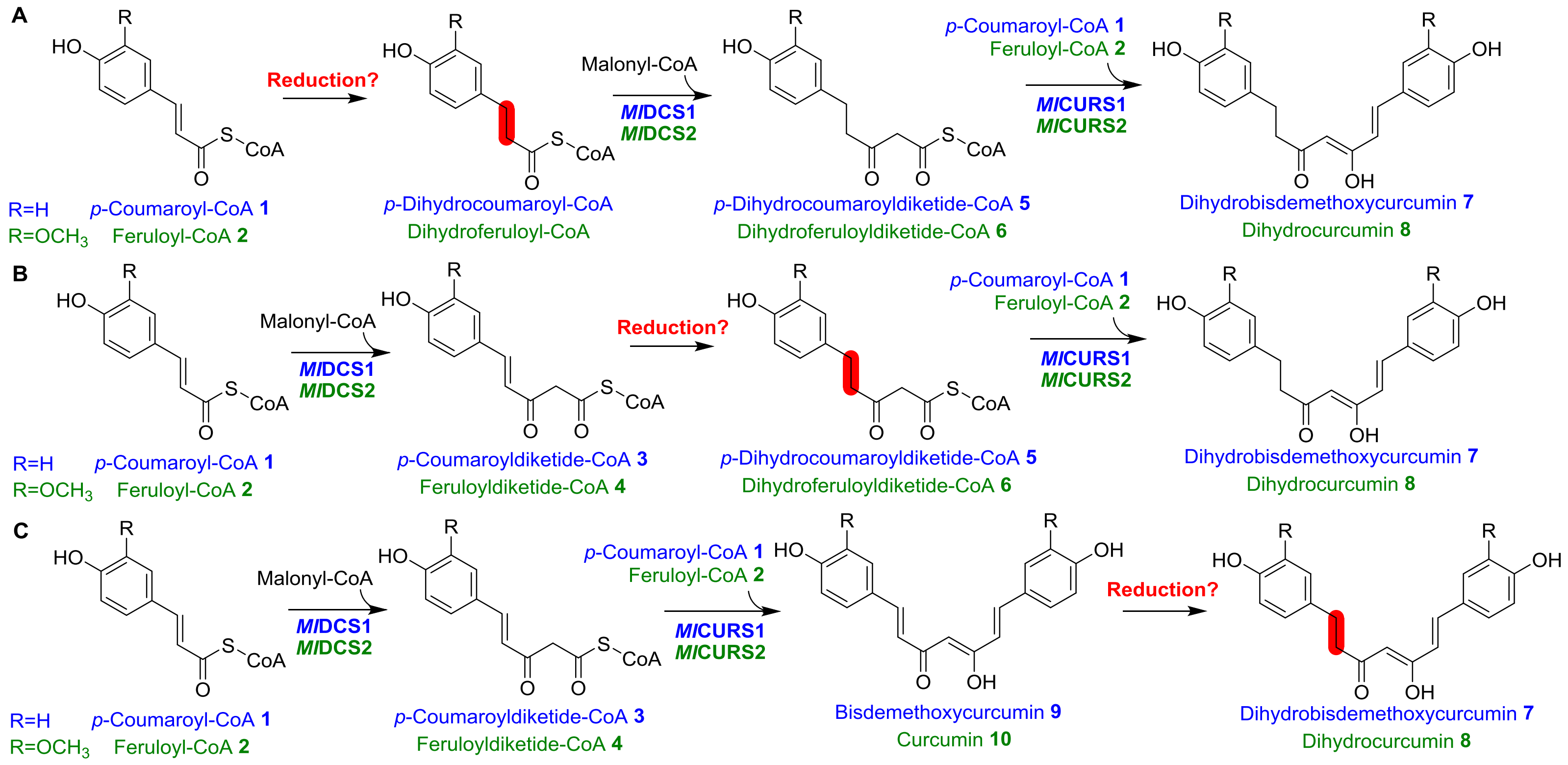


**Figure S8.** Hypothetical formation of dihydrocurcuminoids. The putative double bond reduction step may occur at either the (A) phenylpropanoid-CoA, (B) diketide-CoA, or (C) curcuminoid stage.

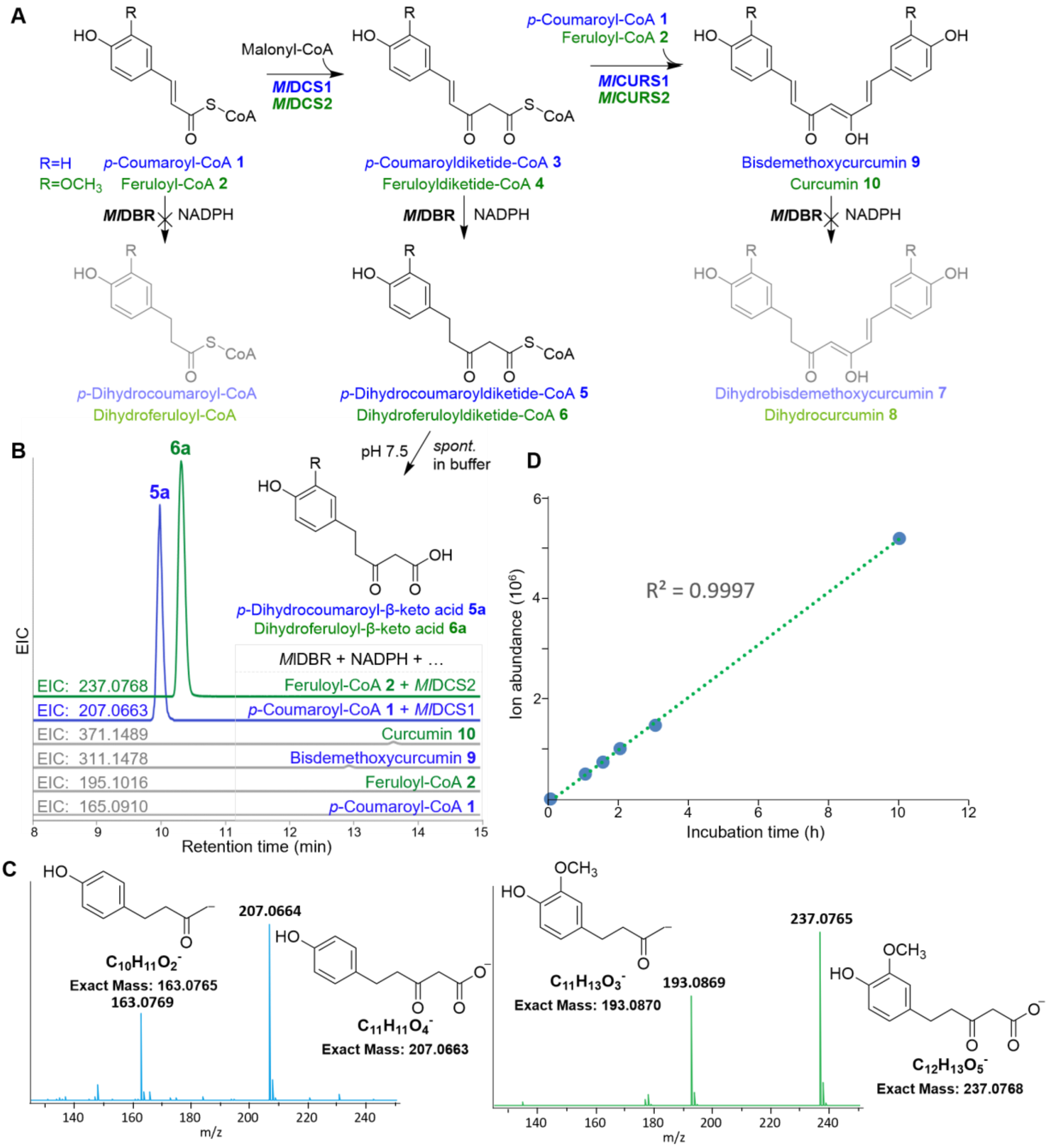


**Figure S9.** Functional characterization of *Ml*DBR. (A) Reaction scheme showing the three possible substrate stages of *Ml*DBR. (B) Extracted ion chromatograms (EIC) of the corresponding products. (C) MS/MS spectra and putative ion fragments of the generated *p*-dihydrocoumaroyl-β-keto acid **5a** and dihydroferuloyl-β-keto acid **6a**. (D) Spontaneous hydrolysis of dihydroferuloyldiketide-CoA **6** in pH 7.5 buffer exhibited a linear correlation between **6a** formation and incubation time (from 1 h to 10 h).

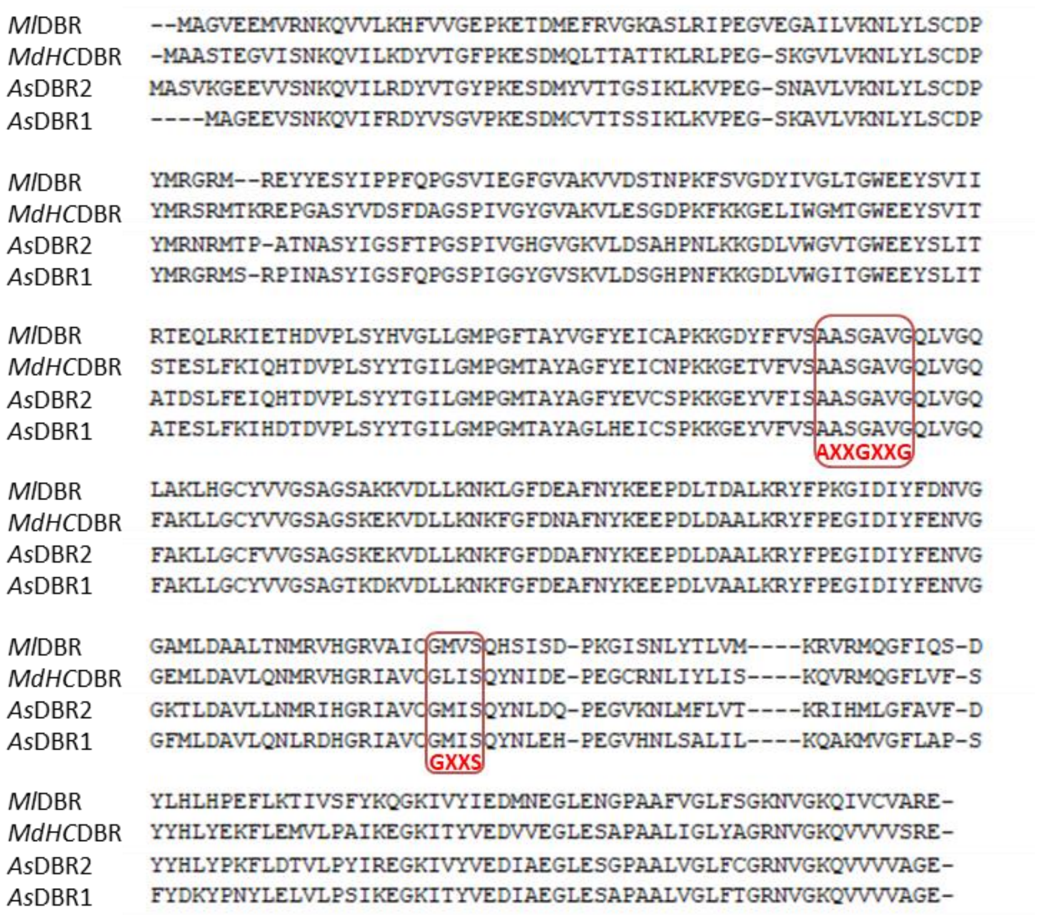


**Figure S10.** Amino acid sequence alignment of *Ml*DBR and other DBRs. The conserved cofactor binding motifs AXXGXXG and GXXS of zinc-independent medium-chain dehydrogenase/reductase superfamily members are shown in boxes.

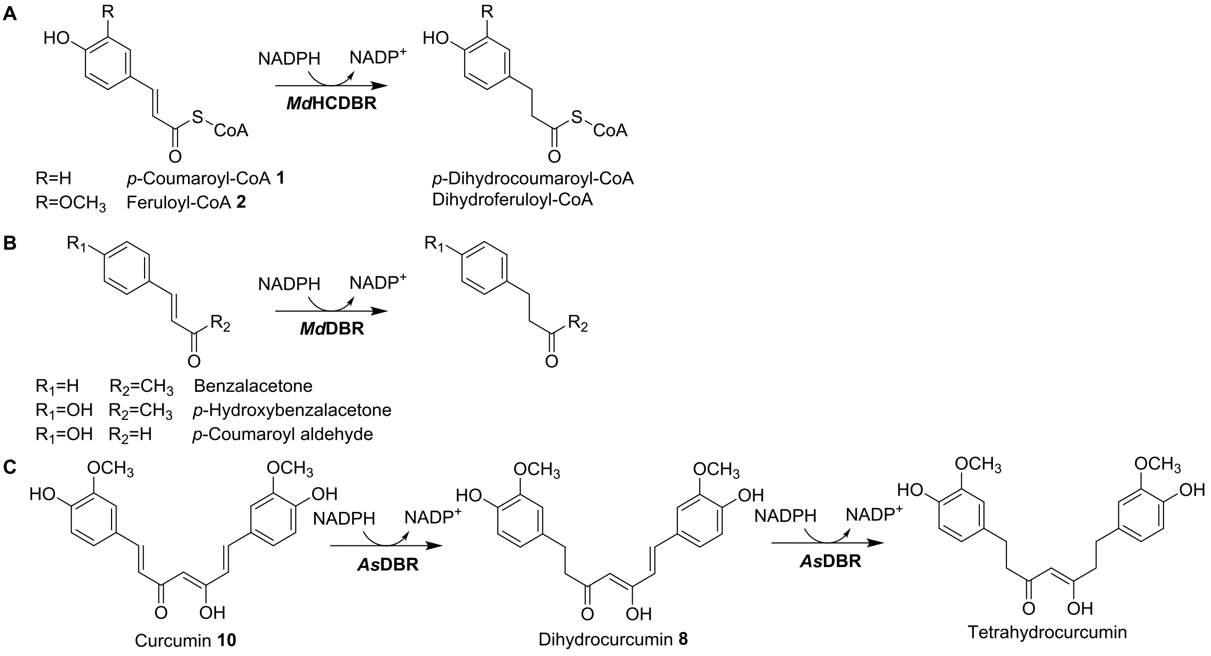


**Figure S11.** Examples of reported plant DBRs. (A) Ibadh *et al*. identified *Md*HCDBR in apple, which reduces *p*-coumaroyl-CoA to *p*-dihydrocoumaroyl-CoA and feruloyl-CoA to dihydroferuloyl-CoA.[19] (B) Caliandro *et al*. described the crystal structure of *Md*HCDBR and tested its activity. The enzyme showed no activity on *p*-coumaroyl-CoA or feruloyl-CoA, but did show activity on benzalacetone, *p*-hydroxybenzalacetone, and *p*-coumaroyl aldehyde. Thus, *Md*HCDBR was renamed to *Md*DBR.[20] (C) Takemoto *et al*. identified *As*DBR from *A. sieboldiana*, which catalyzes the two-step reduction of curcumin **10** to dihydrocurcumin **8** and then further to tetrahydrocurcumin.[21]

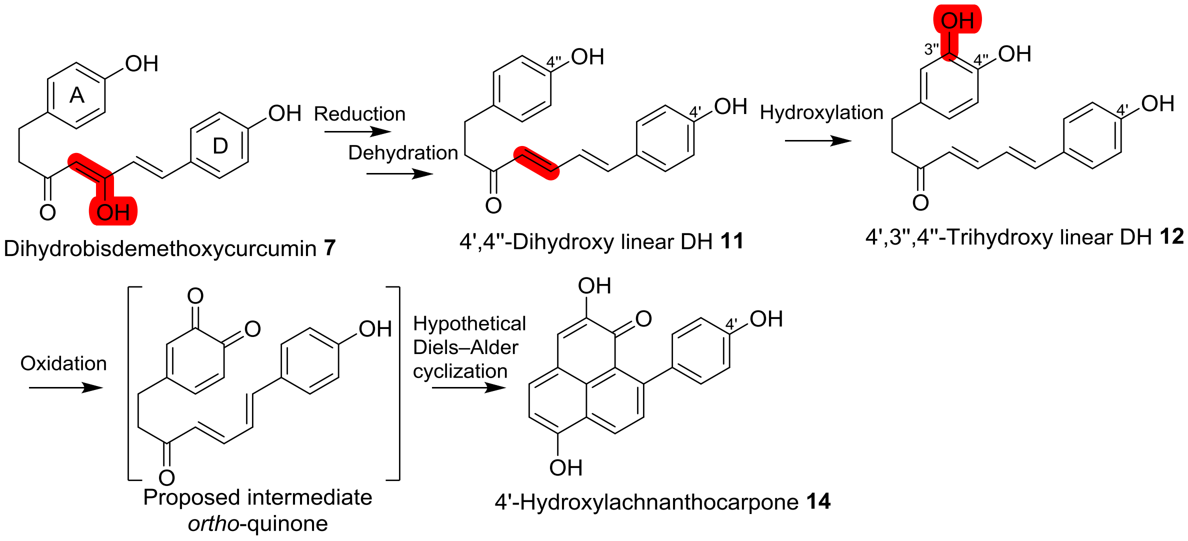


**Figure S12.** The putative biosynthetic pathway from dihydrobisdemethoxycurcumin **7** to the corresponding PP, 4'-hydroxylachnanthocarpone **14**.

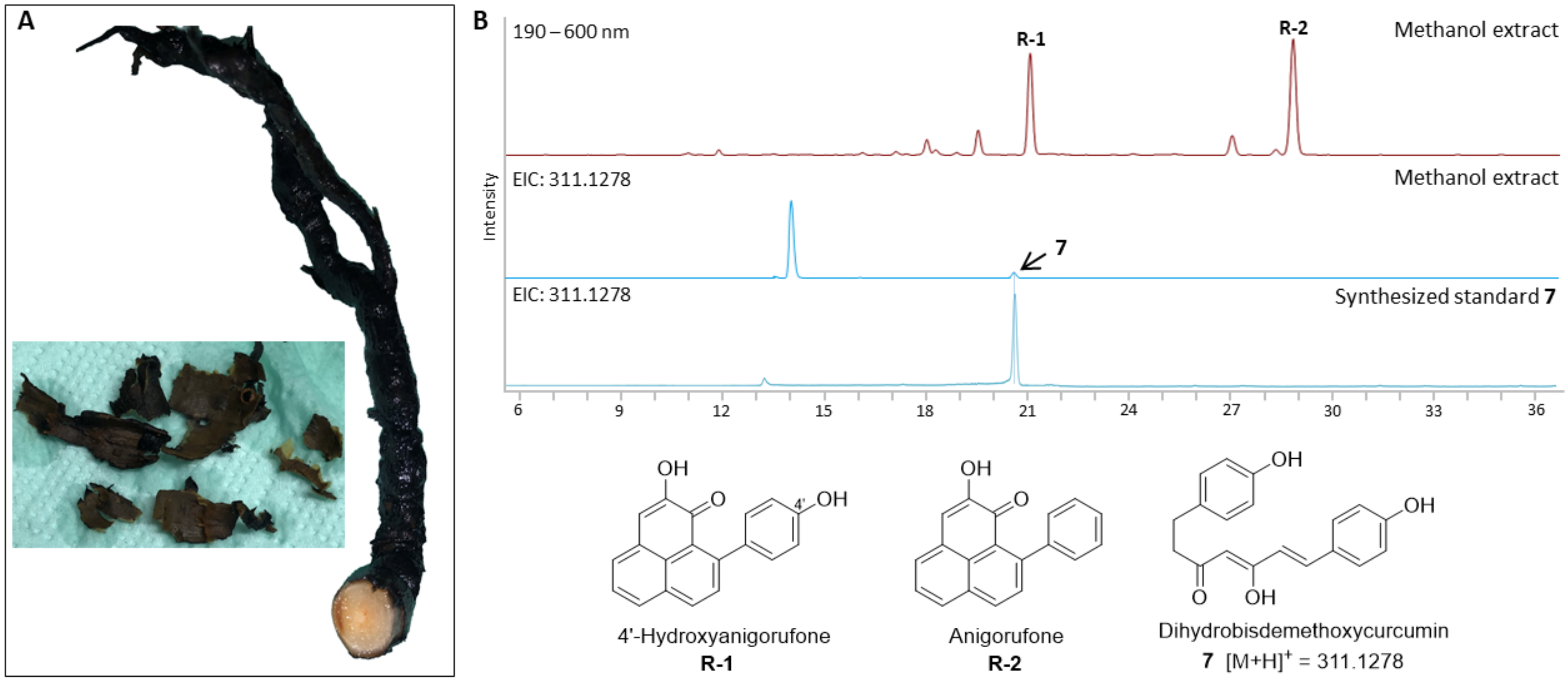


**Figure S13.** PPs are the major metabolites of the outer layer of *M. lasiocarpa* roots. (A) *M. lasiocarpa* root and separated dark outer layer. (B) 4'-Hydroxyanigorufone **R-1** and anigorufone **R-2** are the major metabolites in the methanolic extract of the dark outer layer of the root. Dihydrobisdemethoxycurcumin **7** is present among the trace metabolites.

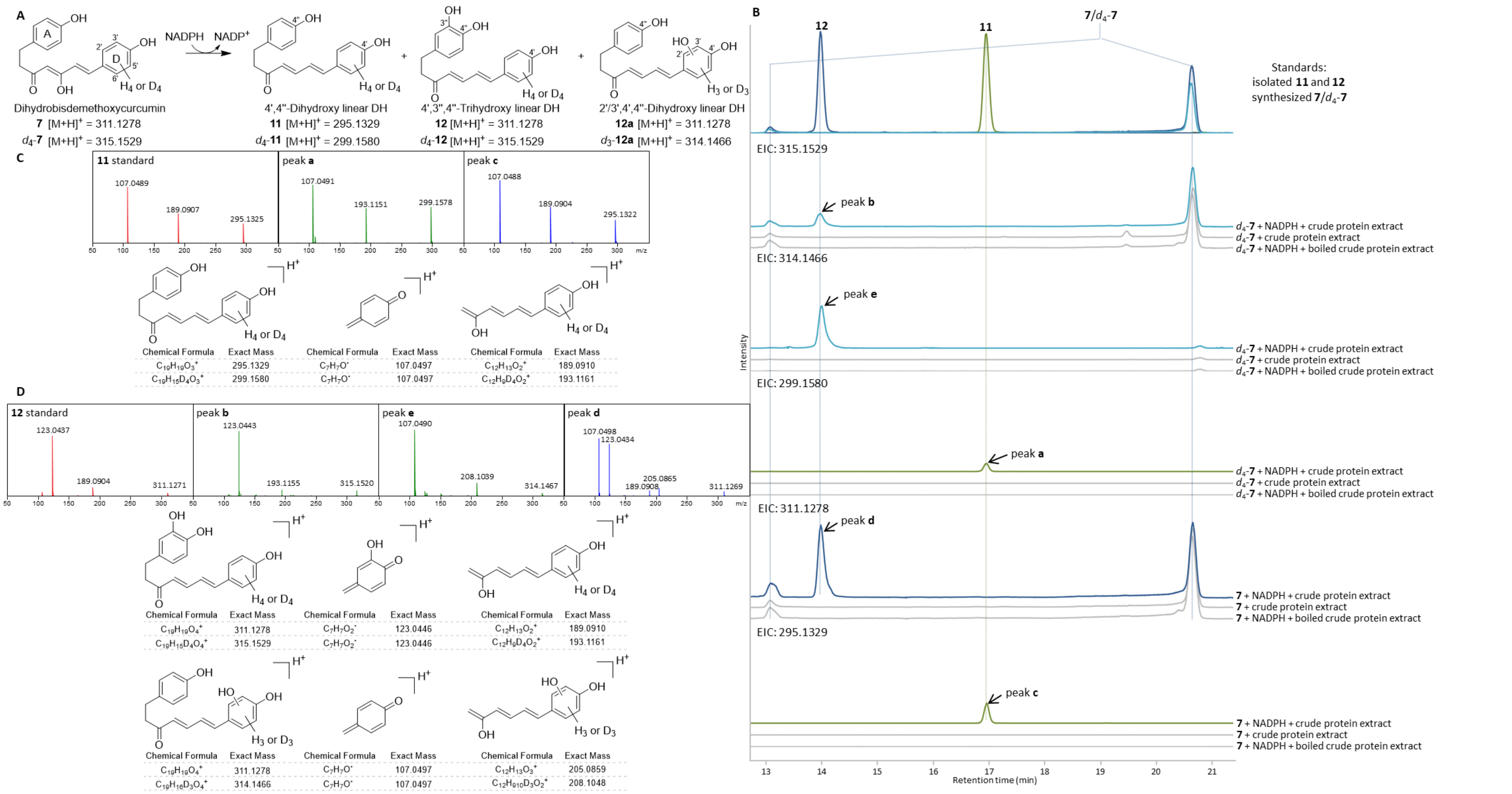

**Figure S14.** *In vitro* assays with dihydrobisdemethoxycurcumin **7**/$d_4$-**7** using crude protein extract from the outer layer of *M. lasiocarpa* roots. (A) Summary of detected enzymatic products. (B) Extracted ion chromatograms (EIC) obtained from the incubation of **7** ($[M+H]^+$ = *m/z* 311.1278 ± 0.005)/$d_4$-**7** ($[M+H]^+$ = *m/z* 315.1529 ± 0.005) with the crude protein extract in the presence of NADPH showing the formation of 4',4''-dihydroxy linear DH **11** ($[M+H]^+$ = *m/z* 295.1329 ±

0.005; peak **c**)/$d_4$-**11** ([M+H]$^+$ = *m/z* 299.1580 ± 0.005; peak **a**), 4',3'',4''-trihydroxy linear DH **12** ([M+H]$^+$ = *m/z* 311.1278 ± 0.005; peak **d**)/$d_4$-**12** ([M+H]$^+$ = *m/z* 315.1529 ± 0.005; peak **b**), and 2'/3',4',4''-trihydroxy linear DH **12a** ([M+H]$^+$ = *m/z* 311.1278 ± 0.005; peak **d**)/$d_3$-**12a** ([M+H]$^+$ = *m/z* 314.1466 ± 0.005; peak **e**). No activity was observed in control reactions without NADPH or using boiled crude protein extract. Dihydrobisdemethoxycurcumin **7**/$d_4$-**7** were observed as two peaks due to the presence of diketo–ketoenol tautomers (**Figure S15**). This experiment was repeated three times with similar results as summarized in **Figure 2**. (C) MS/MS (15 eV) spectra and putative ion fragments of the generated 4',4''-dihydroxy linear DH $d_4$-**11** (peak **a**, green) and **11** (peak **c**, blue), compared to standard **11** (red). (D) MS/MS (20 eV) spectra and putative ion fragments of the generated 4',3'',4''-trihydroxy linear DH $d_4$-**12** (peak **b**, green), 2'/3',4',4''-trihydroxy linear DH $d_3$-**12a** (peak **e**, green), and **12**/**12a** (peak **d**, blue), compared to standard **12** (red).

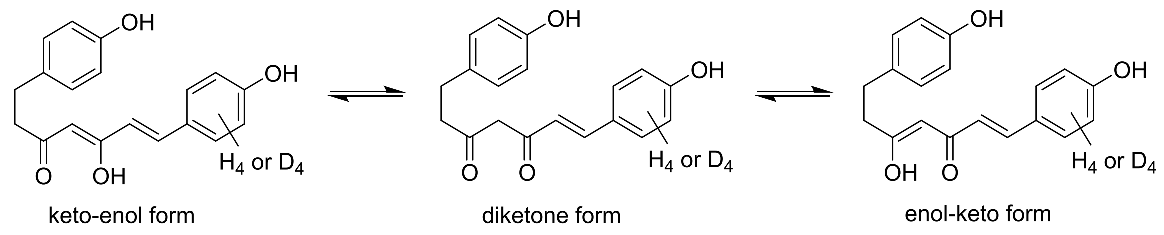


**Figure S15.** Diketo–ketoenol tautomers of dihydrobisdemethoxycurcumin **7**/$d_4$-**7**.

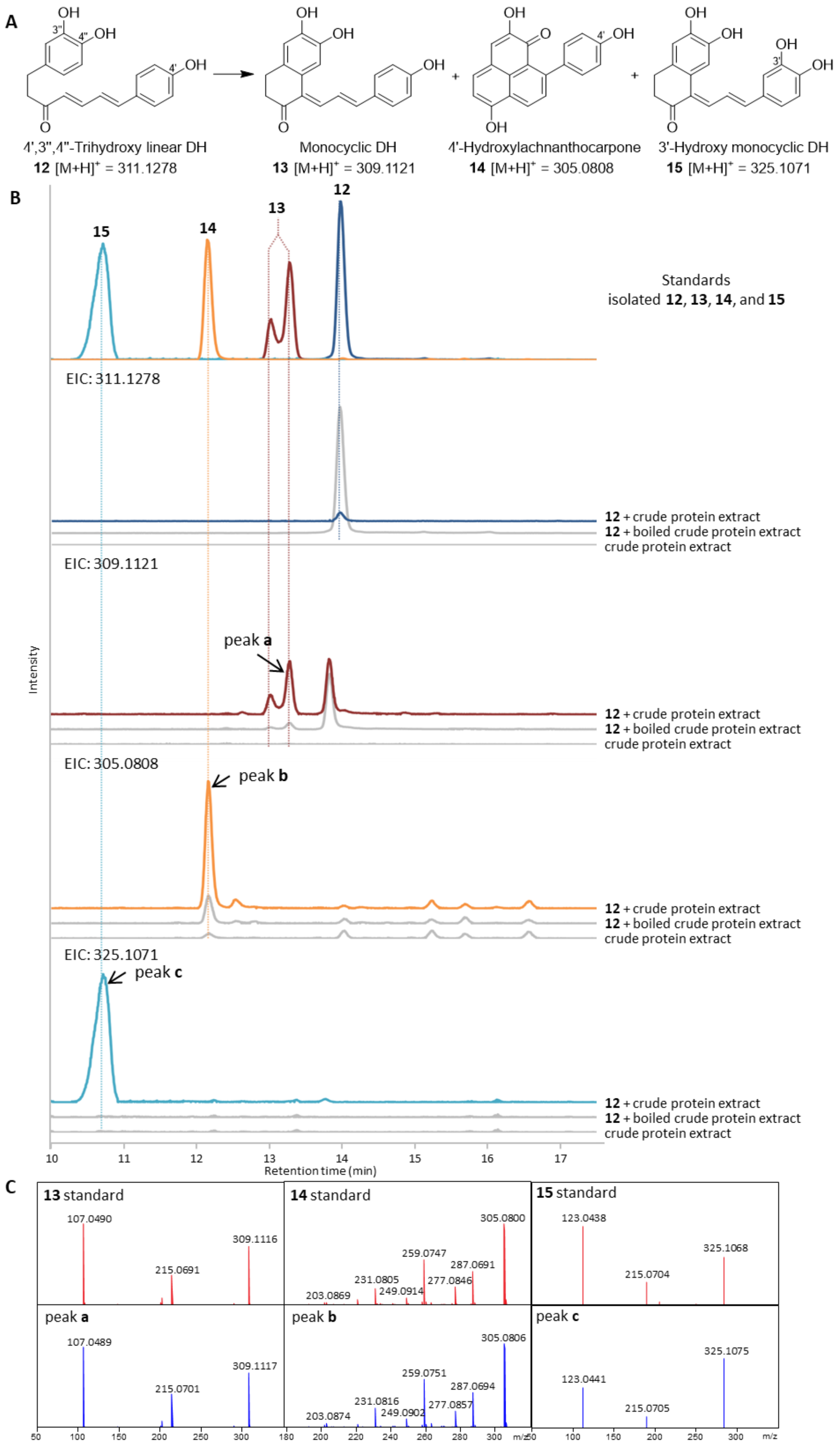


**Figure S16.** *In vitro* assays with 4',3'',4''-trihydroxy linear DH **12** using crude protein extract from the outer layer of *M. lasiocarpa* roots. (A) Summary of detected enzymatic products. (B) Extracted ion chromatograms (EIC) obtained from the incubation of **12** ($[M+H]^+$ = *m/z* 311.1278 ± 0.005) with the crude protein extract showing the formation of monocyclic DH **13** ($[M+H]^+$ = *m/z* 309.1121 ± 0.005;

peak **a**), 4'-hydroxylachnanthocarpone **14** ($[M+H]^+$ = *m/z* 305.0808 ± 0.005; peak **b**), and 3'-hydroxy monocyclic DH **15** ($[M+H]^+$ = *m/z* 325.1071 ± 0.005; peak **c**). No activity was observed in control reactions without substrate or using boiled crude protein extract. Compound **13**/**15** was observed as a double or broad peak due to the presence of *E*/*Z* isomers (**Figure S18**). This experiment was repeated three times with similar results as summarized in **Figure 2**. (C) MS/MS (10 eV) spectra of the generated monocyclic DH **13** (peak **a**, blue) compared to standard **13** (red). MS/MS (35 eV) spectra of the generated 4'-hydroxylachnanthocarpone **14** (peak **b**, blue) compared to standard **14** (red). MS/MS (10 eV) spectra of the generated 3'-hydroxy monocyclic DH **15** (peak **c**, blue) compared to standard **15** (red).

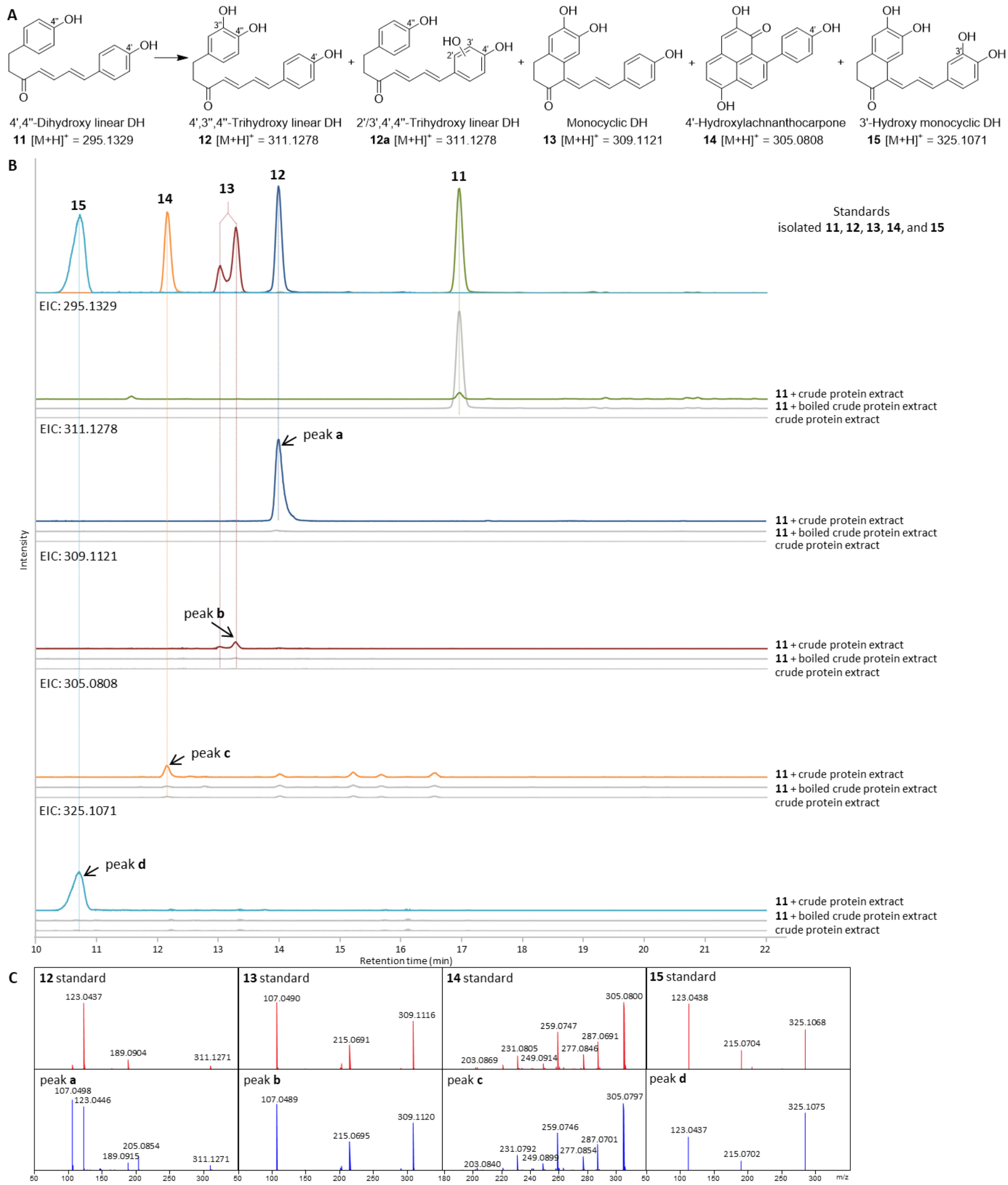


**Figure S17.** *In vitro* assays with 4',4''-dihydroxy linear DH **11** using crude protein extract from the outer layer of *M. lasiocarpa* roots. (A) Summary of detected enzymatic products. (B) Extracted ion chromatograms (EIC) obtained from the incubation of **11** ($[M+H]^+$ = *m/z* 295.1329 ± 0.005) with the crude protein extract showing the formation of 4',3'',4''-trihydroxy linear DH **12** ($[M+H]^+$ = *m/z* 311.1278 ± 0.005; peak **a**), 2'/3',4',4''-trihydroxy linear DH **12a** ($[M+H]^+$ = *m/z* 311.1278 ± 0.005; peak **a**), monocyclic DH **13** ($[M+H]^+$ = *m/z* 309.1121 ± 0.005; peak **b**), 4'-hydroxylachnanthocarpone **14** ($[M+H]^+$ = *m/z* 305.0808 ± 0.005; peak **c**), and 3'-hydroxy monocyclic DH **15** ($[M+H]^+$ = *m/z* 325.1071 ± 0.005; peak **d**). No activity was observed in control reactions without substrate or using boiled crude protein extract. Compound **13**/**15** was observed as a double or broad peak due to the presence of *E*/*Z* isomers (**Figure S18**). This experiment was repeated three times with similar results as summarized in

**Figure 2**. (C) MS/MS (15 eV) spectra of the generated 4',3'',4''-trihydroxy linear DH **12** and 2'/3',4',4''-trihydroxy linear DH **12a** (peak **a**, blue) compared to standard **12** (red). MS/MS (10 eV) spectra of the generated monocyclic DH **13** (peak **b**, blue) compared to standard **13** (red). MS/MS (35 eV) spectra of the generated 4'-hydroxylachnanthocarpone **14** (peak **c**, blue) compared to standard **14** (red). MS/MS (10 eV) spectra of the generated 3'-hydroxy monocyclic DH **15** (peak **d**, blue) compared to standard **15** (red).

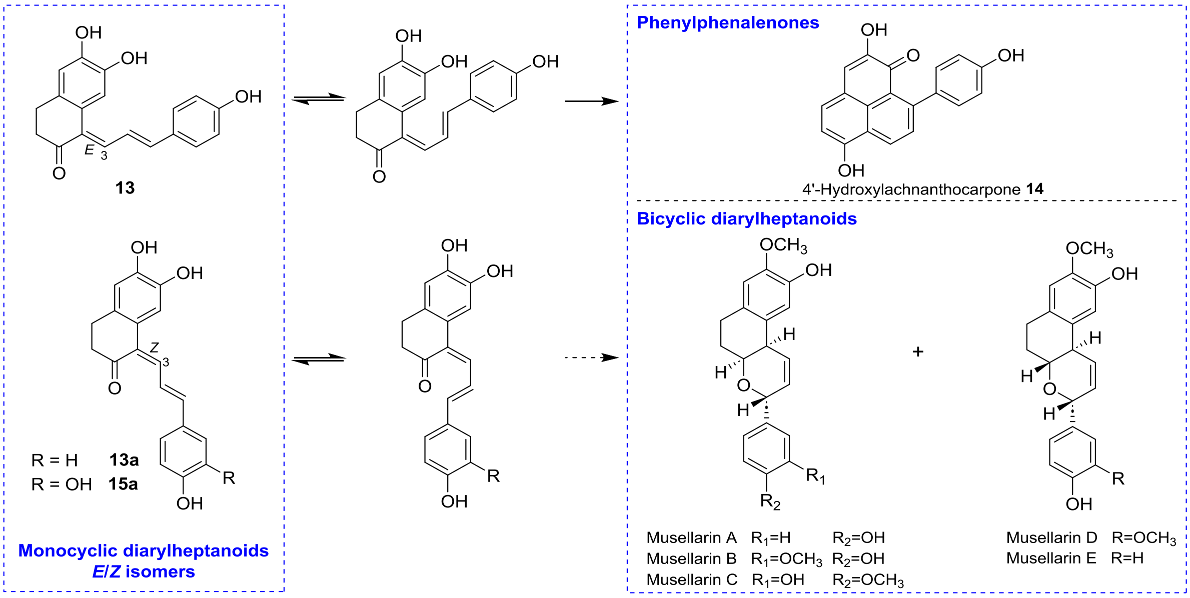


**Figure S18.** Monocyclic diarylheptanoids are the precursors of phenylphenalenones and bicyclic diarylheptanoids (musellarins A–E).

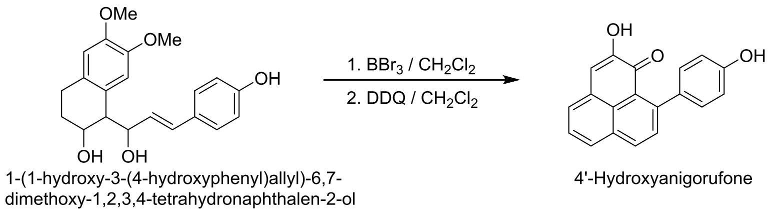


**Figure S19.** Reported chemical conversion of a monocyclic diarylheptanoid into 4'-hydroxyanigorufone by demethylation and subsequent oxidation. [23]

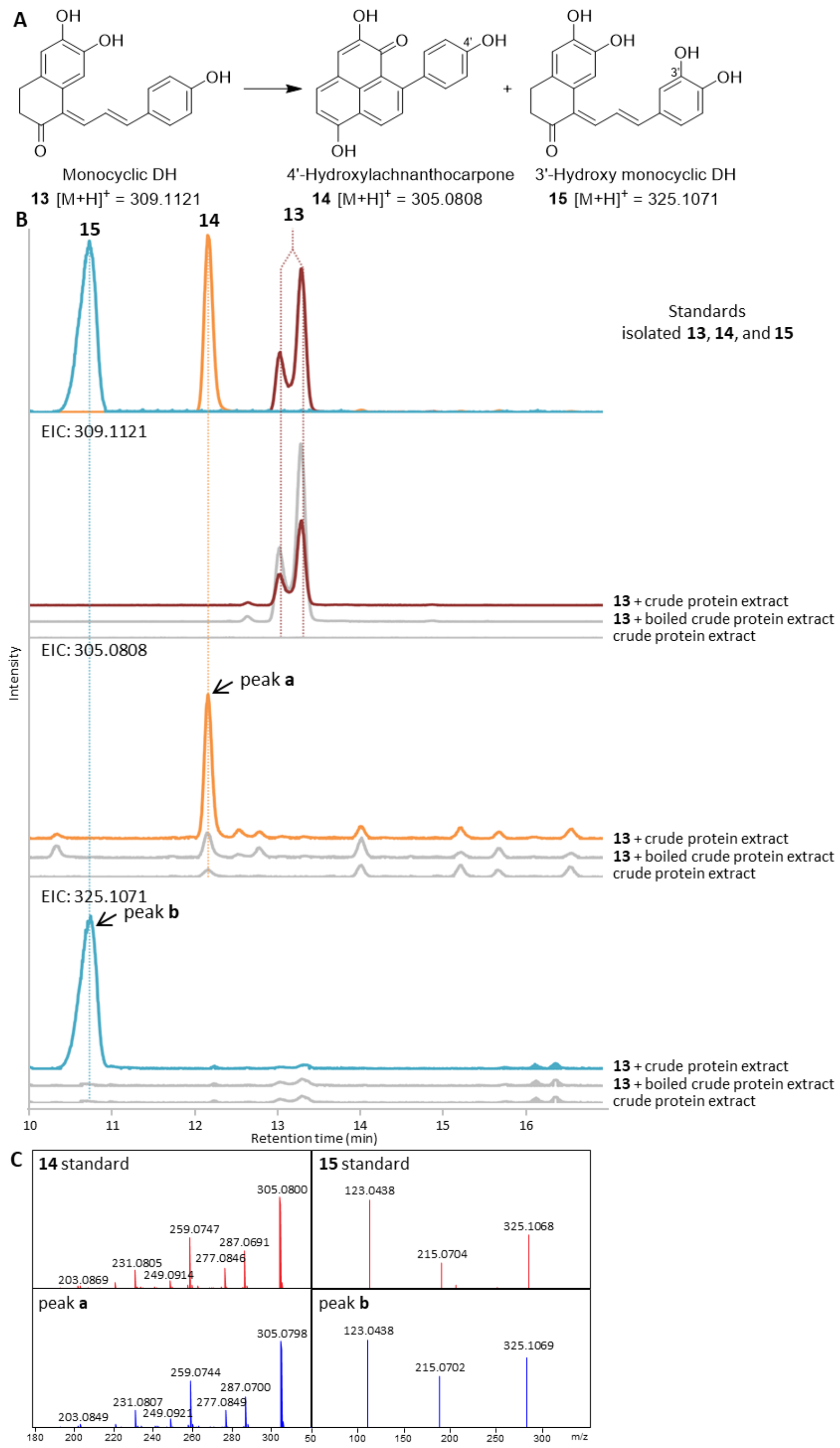


**Figure S20.** *In vitro* assays with monocyclic DH **13** using crude protein extract from the outer layer of *M. lasiocarpa* roots. (A) Summary of detected enzymatic products. (B) Extracted ion chromatograms (EIC) obtained from the incubation of **13** ($[M+H]^+$ = *m/z* 309.1121 ± 0.005) with the crude protein extract showing the formation of 4'-hydroxylachnanthocarpone **14** ($[M+H]^+$ = *m/z* 305.0808 ± 0.005; peak **a**) and 3'-hydroxy monocyclic DH **15** ($[M+H]^+$ = *m/z* 325.1071 ± 0.005; peak **b**). No activity was observed in control reactions without substrate or using boiled crude protein extract. Compound **13**/**15** was observed as a double or broad peak due to the presence of *E*/*Z* isomers (**Figure S18**). This experiment was repeated three times with similar results as summarized in **Figure 2**. (C) MS/MS (35 eV) spectra of the generated 4'-hydroxylachnanthocarpone **14** (peak **a**, blue) compared to standard **14** (red). MS/MS (10 eV) spectra of the generated 3'-hydroxy monocyclic DH **15** (peak **b**, blue) compared to standard **15** (red).

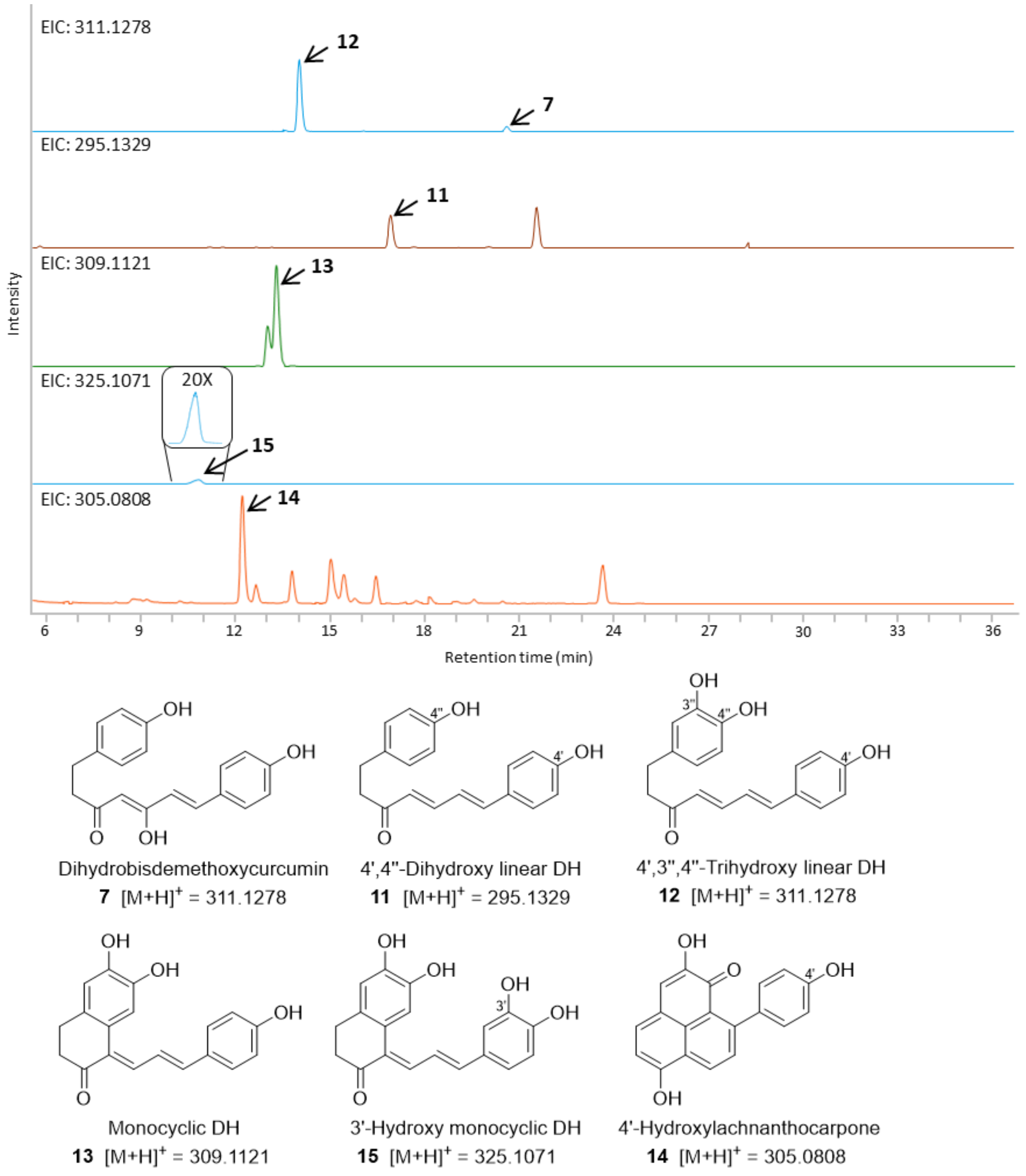


**Figure S21.** Abundance of target metabolites found in the methanolic extract of the dark outer layer of *M. lasiocarpa* roots. Compound **13**/**15** was observed as a double or broad peak due to the presence of *E*/*Z* isomers (**Figure S18**).

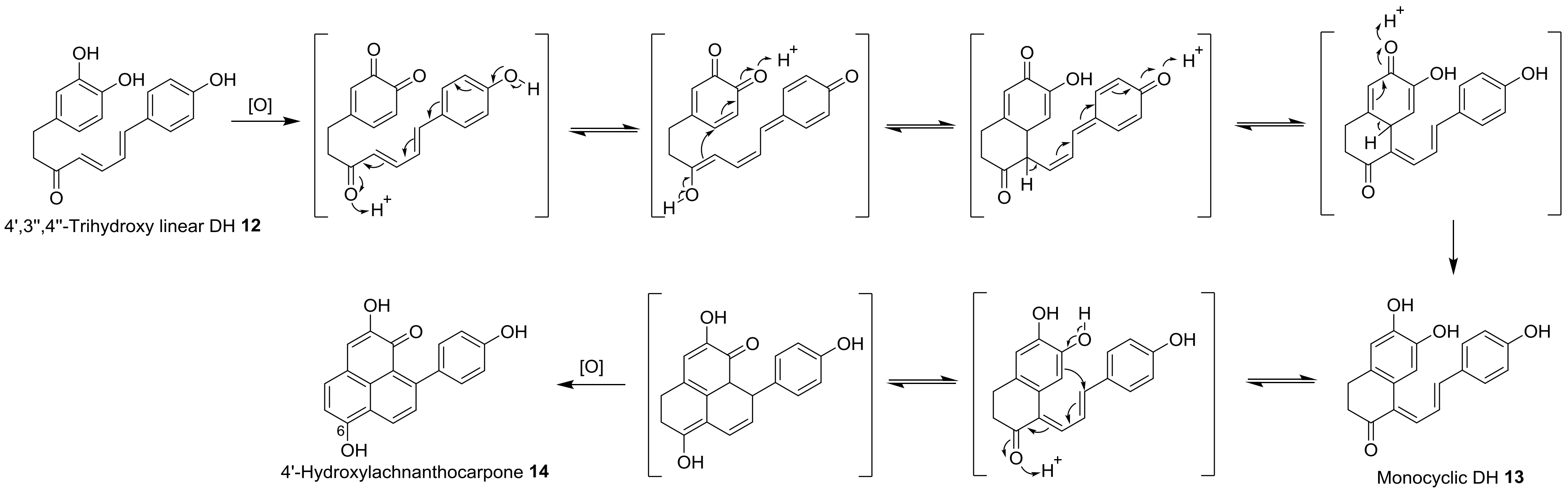


**Figure S22.** Proposed reaction mechanism for the two-step cyclization of 4',3'',4''-trihydroxy linear DH **12** to form the PP scaffold (4'-hydroxylachnanthocarpone **14**).

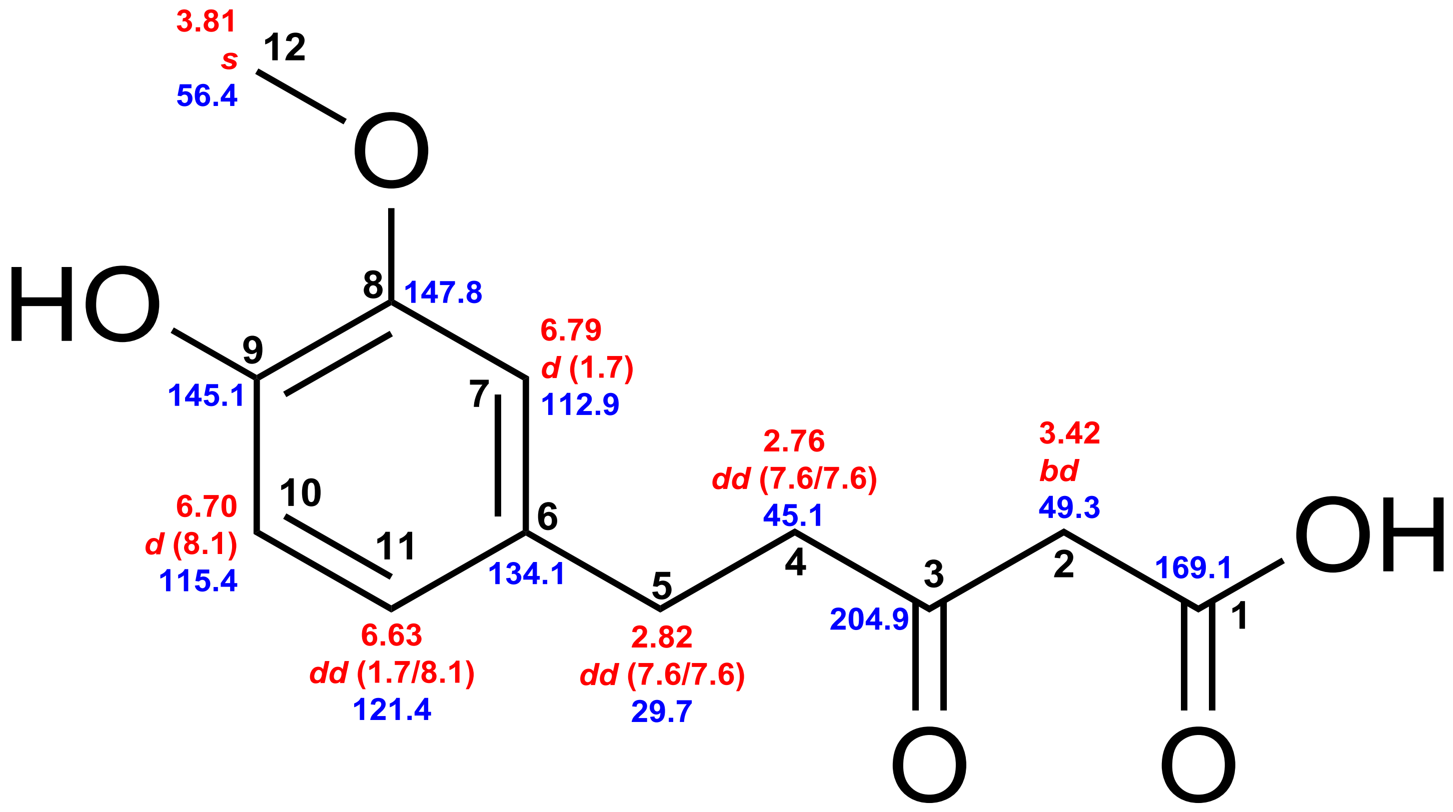


Dihydroferuloyl-β-keto acid **6a**

**Figure S23.** Chemical shifts of dihydroferuloyl-β-keto acid **6a**. Red: $^{1}$H chemical shifts (δ ppm, *mult*., $^{3}J_{HH}$ in Hz). Blue: $^{13}$C chemical shifts (δ ppm).

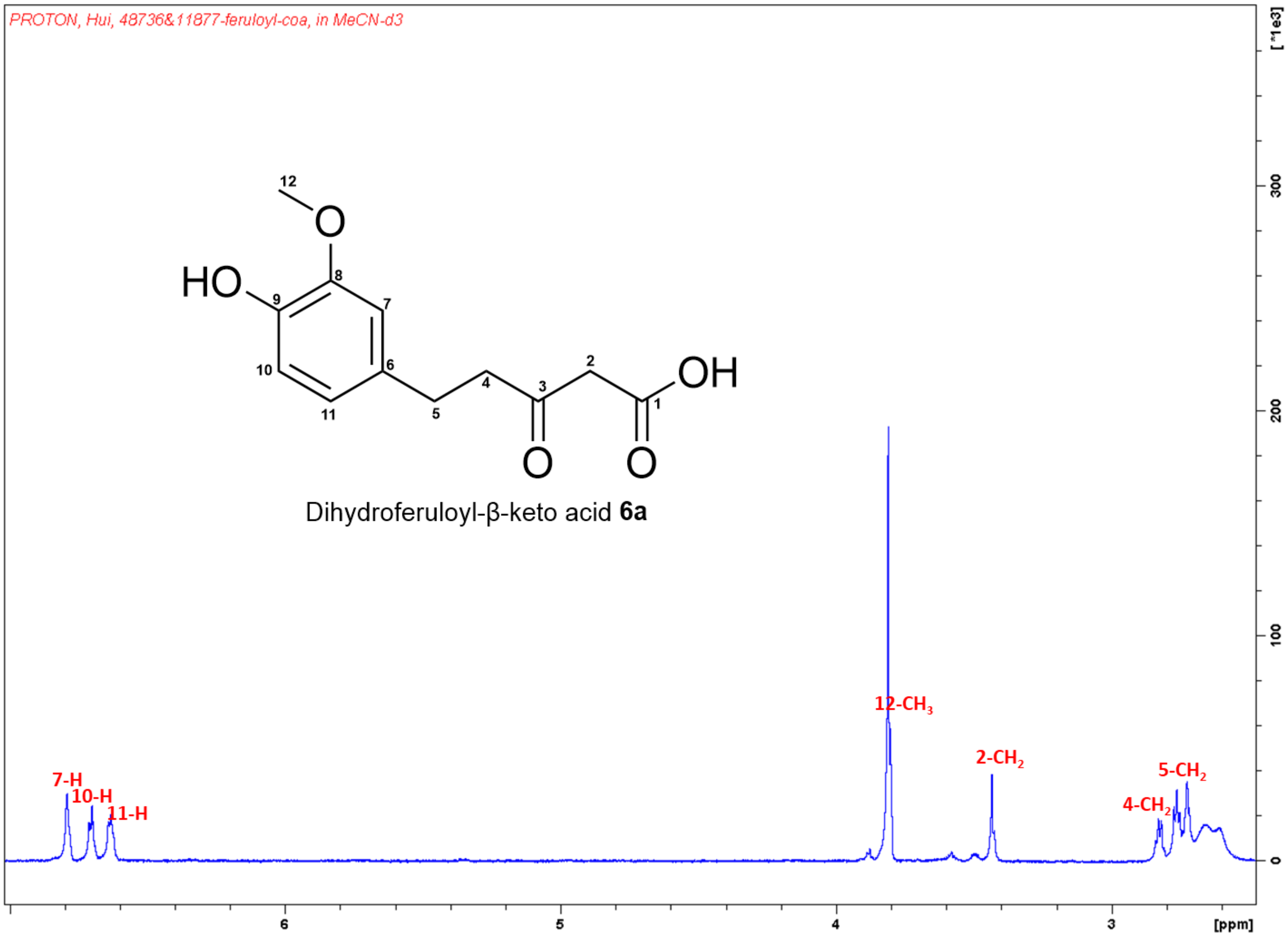


**Figure S24.** $^{1}H$ NMR spectrum (700 MHz, $CD_3CN$) of dihydroferuloyl-β-keto acid **6a**.

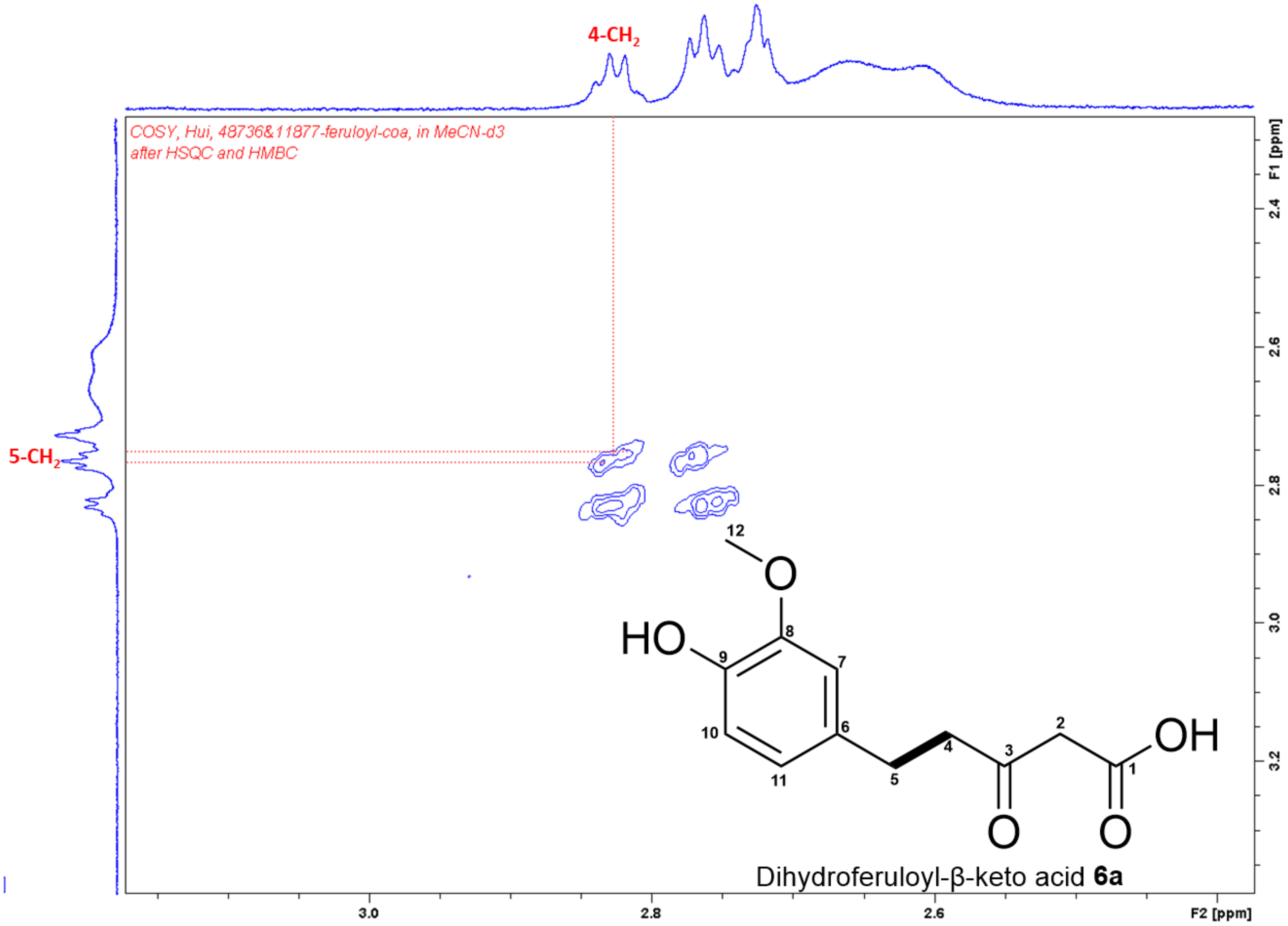


**Figure S25.** $^{1}H$–$^{1}H$ COSY spectrum of dihydroferuloyl-β-keto acid **6a** in $CD_3CN$.

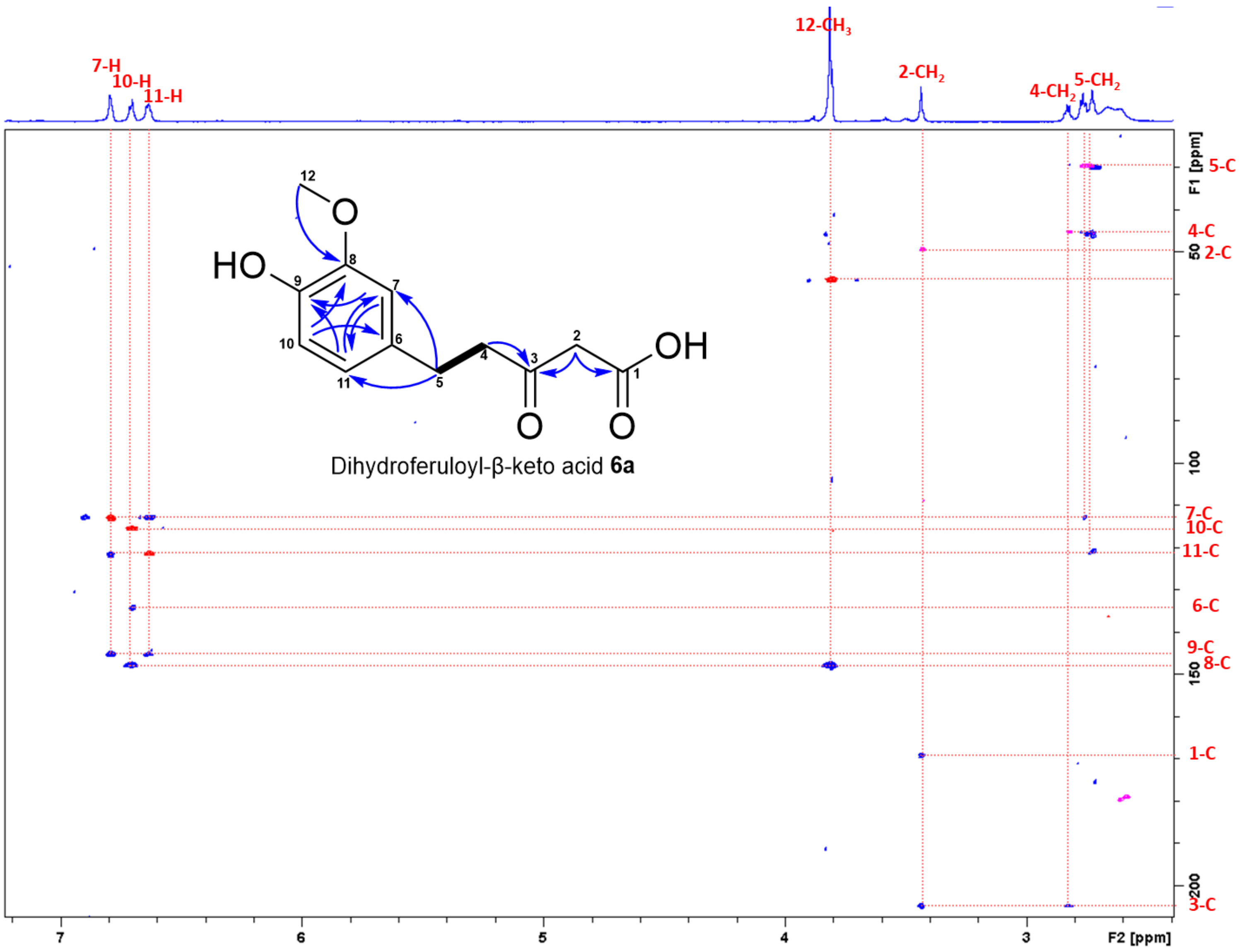


**Figure S26.** Superimposed HSQC and HMBC spectra of dihydroferuloyl-β-keto acid **6a** in $CD_3CN$.

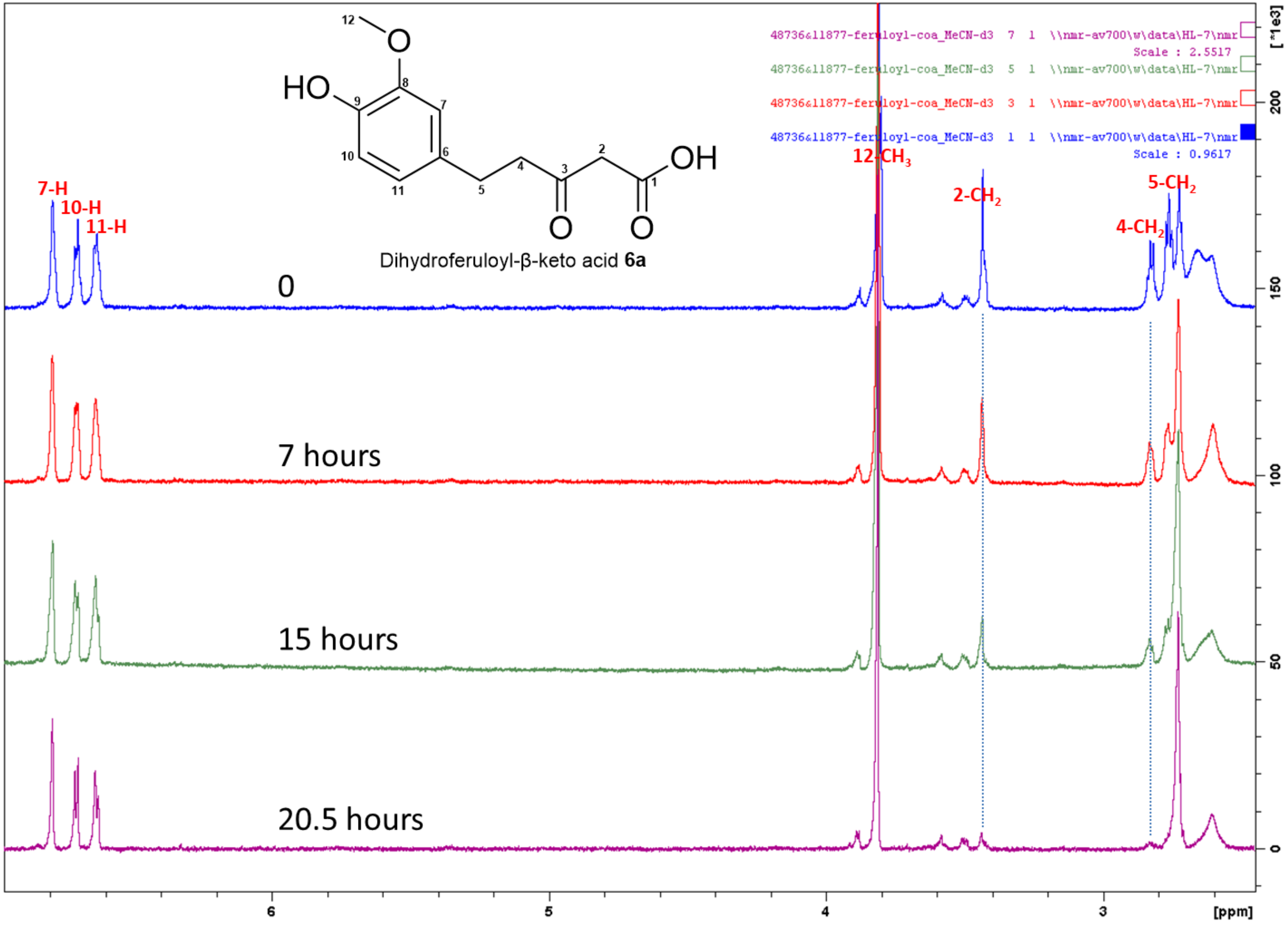


**Figure S27.** Time-dependent [1]H NMR spectra (700 MHz, $CD_3CN$) showing the degradation of dihydroferuloyl-β-keto acid **6a**.

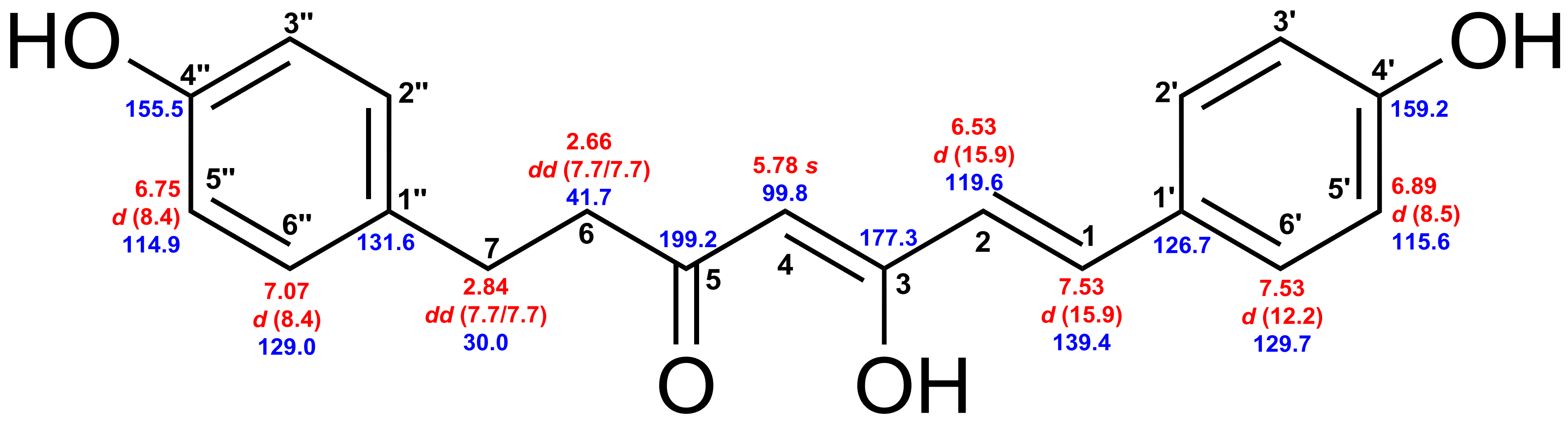


Dihydrobisdemethoxycurcumin **7**

**Figure S28.** Chemical shifts of dihydrobisdemethoxycurcumin **7**. Red: $^{1}$H chemical shifts (δ ppm, *mult*., $^{3}J_{HH}$ in Hz). Blue: $^{13}$C chemical shifts (δ ppm).

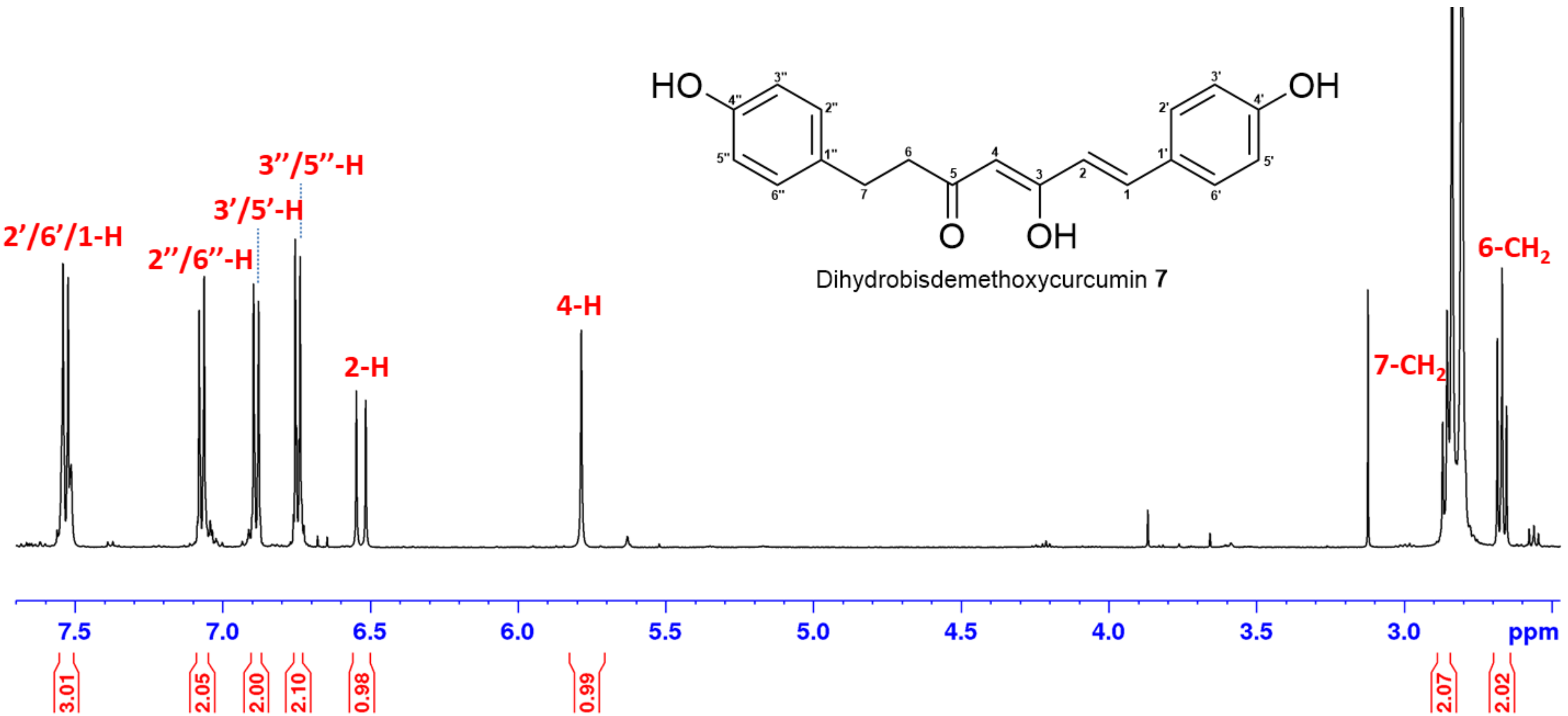


**Figure S29.** $^1$H NMR spectrum (700 MHz, acetone-$d_6$) of dihydrobisdemethoxycurcumin **7**.

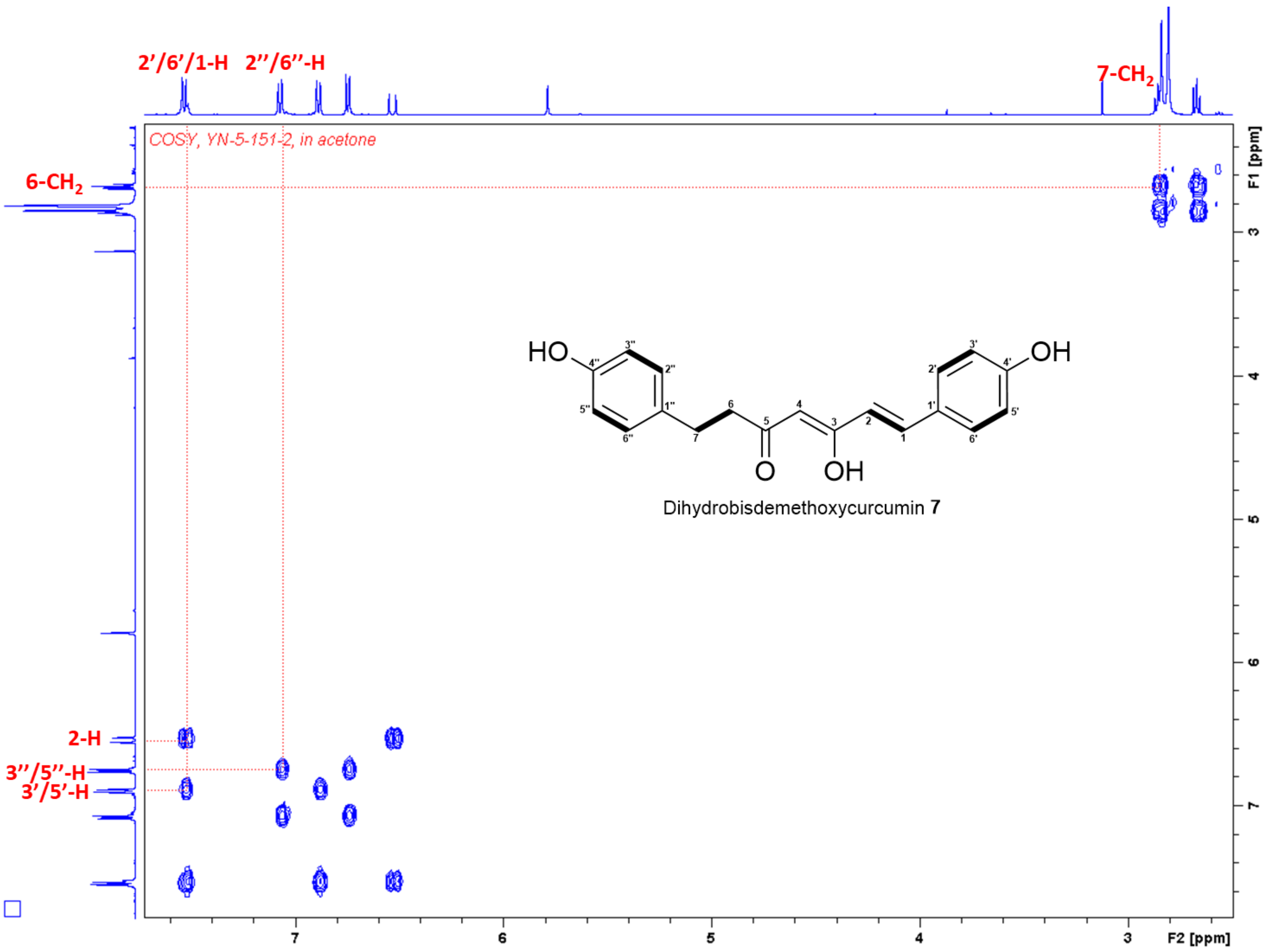


**Figure S30.** $^{1}H$–$^{1}H$ COSY spectrum of dihydrobisdemethoxycurcumin **7** in acetone-$d_6$.

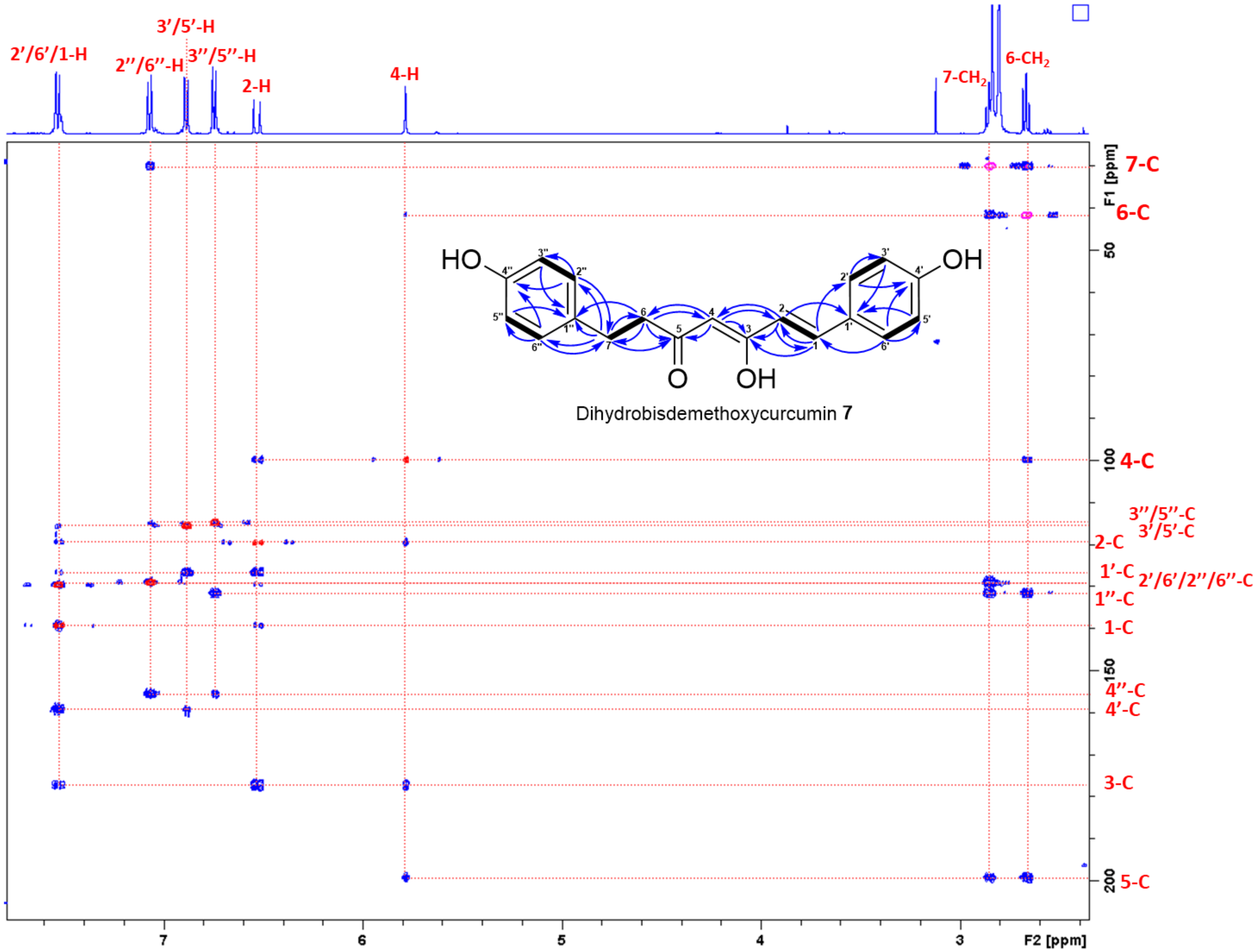


**Figure S31.** Superimposed HSQC and HMBC spectra of dihydrobisdemethoxycurcumin **7** in acetone-$d_6$.

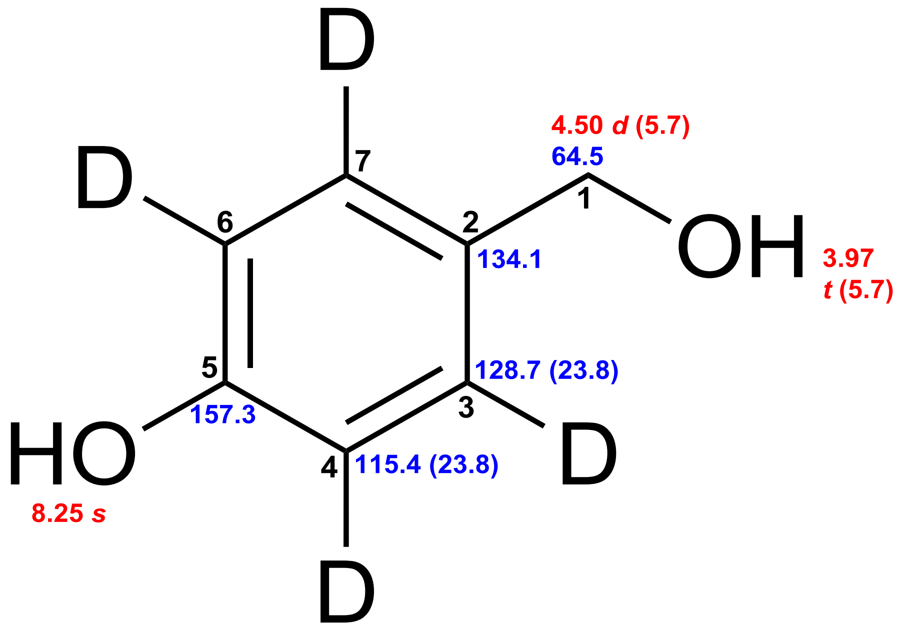


4-Hydroxybenzylalcohol-$d_4$

**Figure 32.** Chemical shifts of 4-hydroxybenzylalcohol-$d_4$. Red: $^1$H chemical shifts (δ ppm, *mult*., $^3J_{HH}$ in Hz). Blue: $^{13}$C chemical shifts (δ ppm, $^1J_{CD}$ in Hz).

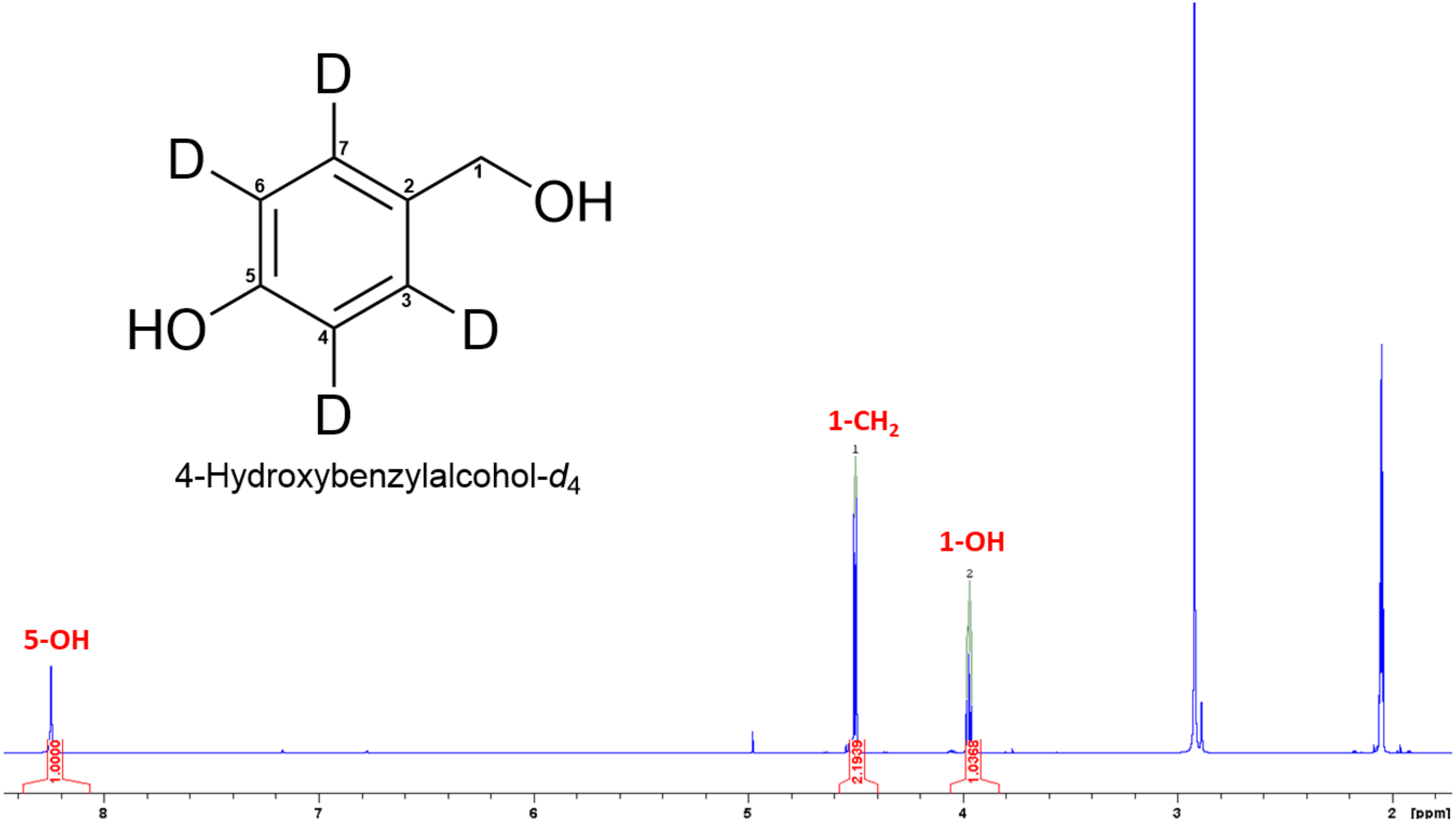


**Figure S33.** $^1$H NMR spectrum (500 MHz, acetone-$d_6$) of 4-hydroxybenzylalcohol-$d_4$.

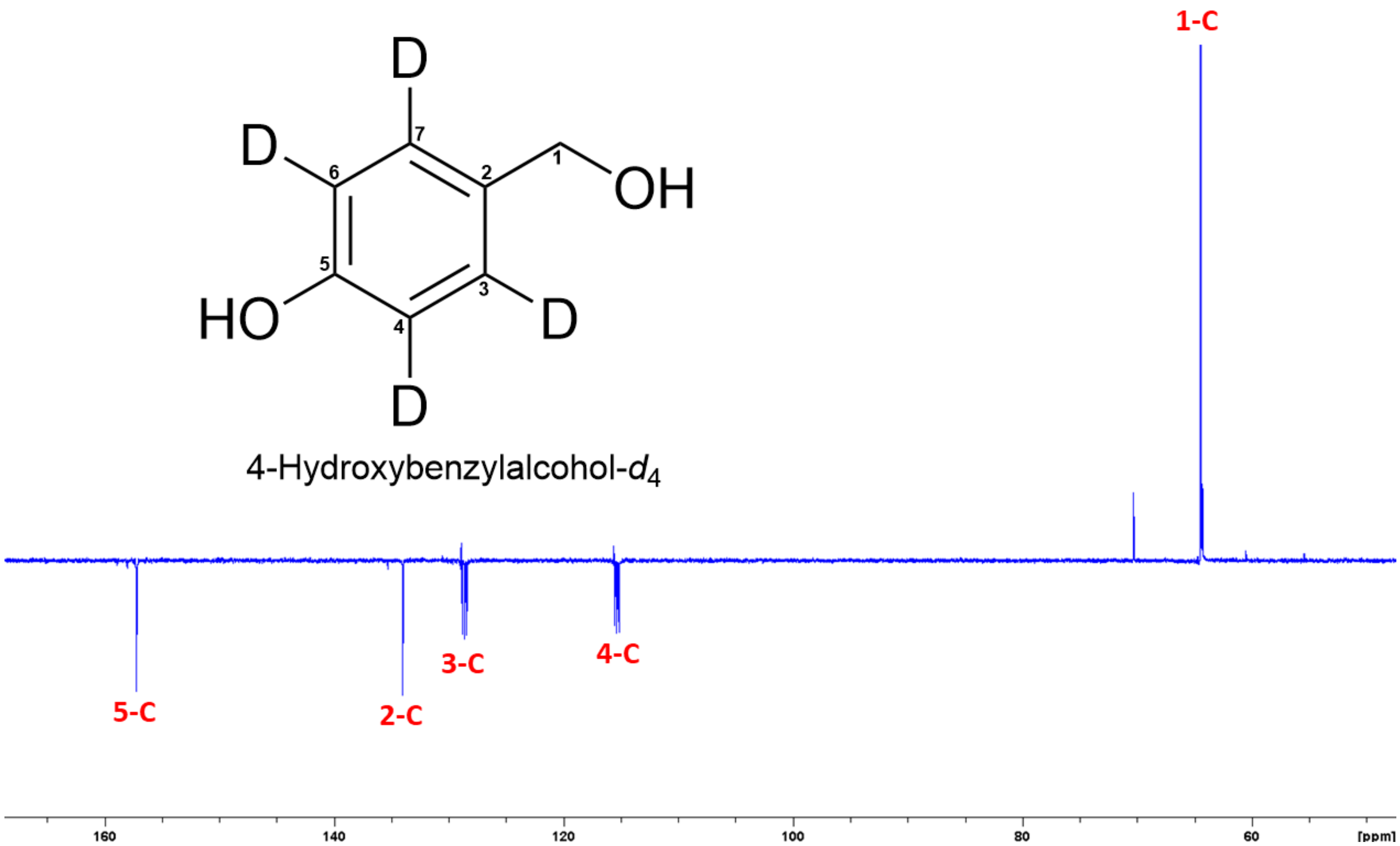


**Figure S34.** DEPTQ spectrum (125 MHz, acetone-$d_6$) of 4-hydroxybenzylalcohol-$d_4$.

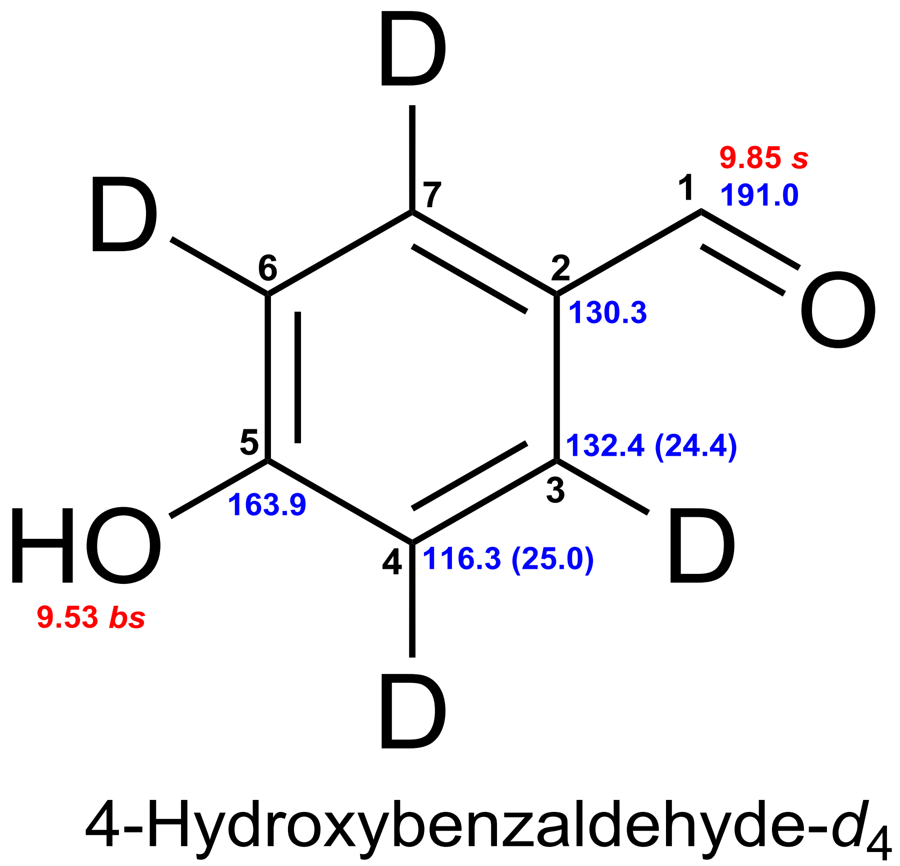


**Figure 35.** Chemical shifts of 4-hydroxybenzaldehyde-$d_4$. Red: $^1$H chemical shifts (δ ppm, *mult.*, $^3J_{HH}$ in Hz). Blue: $^{13}$C chemical shifts (δ ppm, $^1J_{CD}$ in Hz).

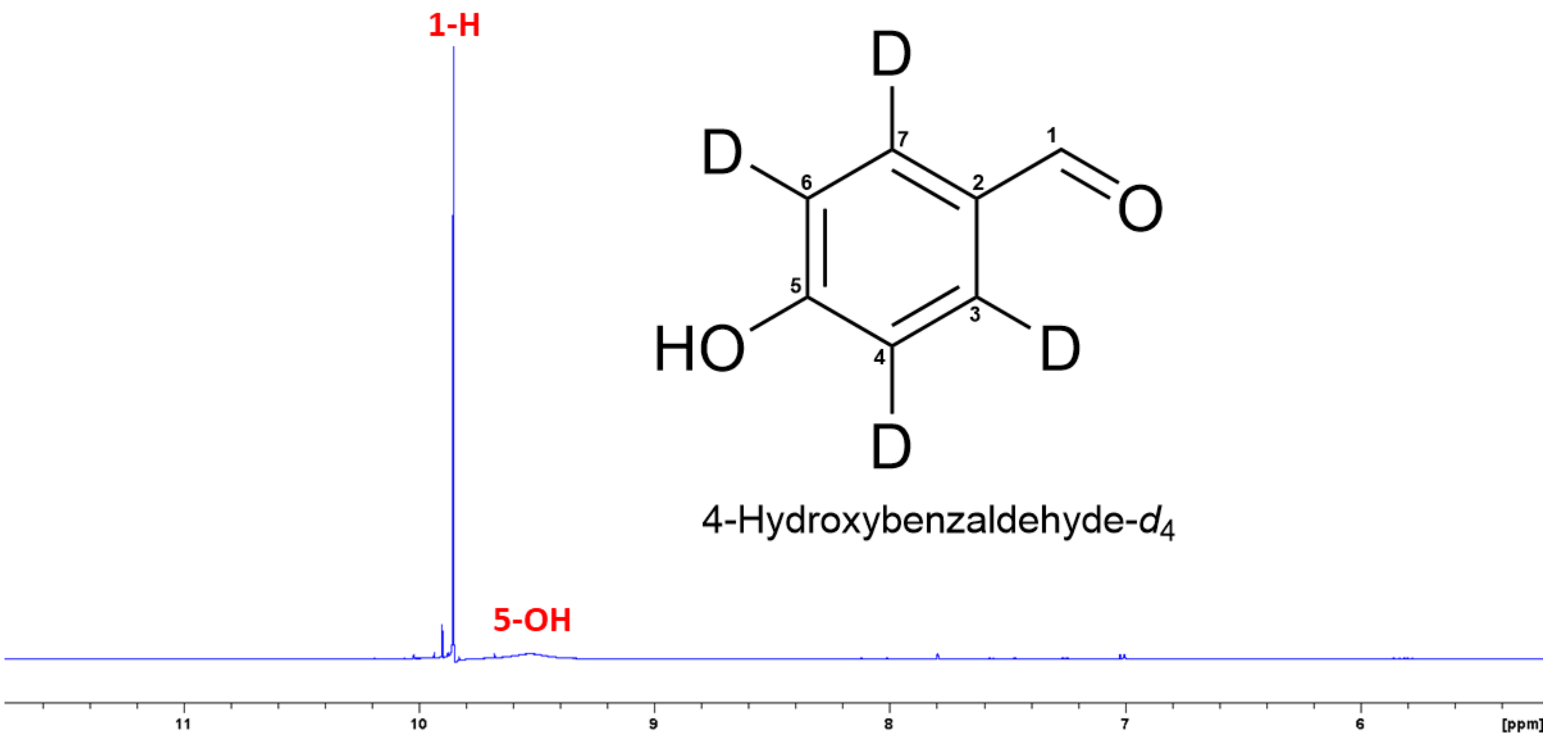


**Figure S36.** $^{1}$H NMR spectrum (500 MHz, acetone-$d_6$) of 4-hydroxybenzylalcohol-$d_4$.

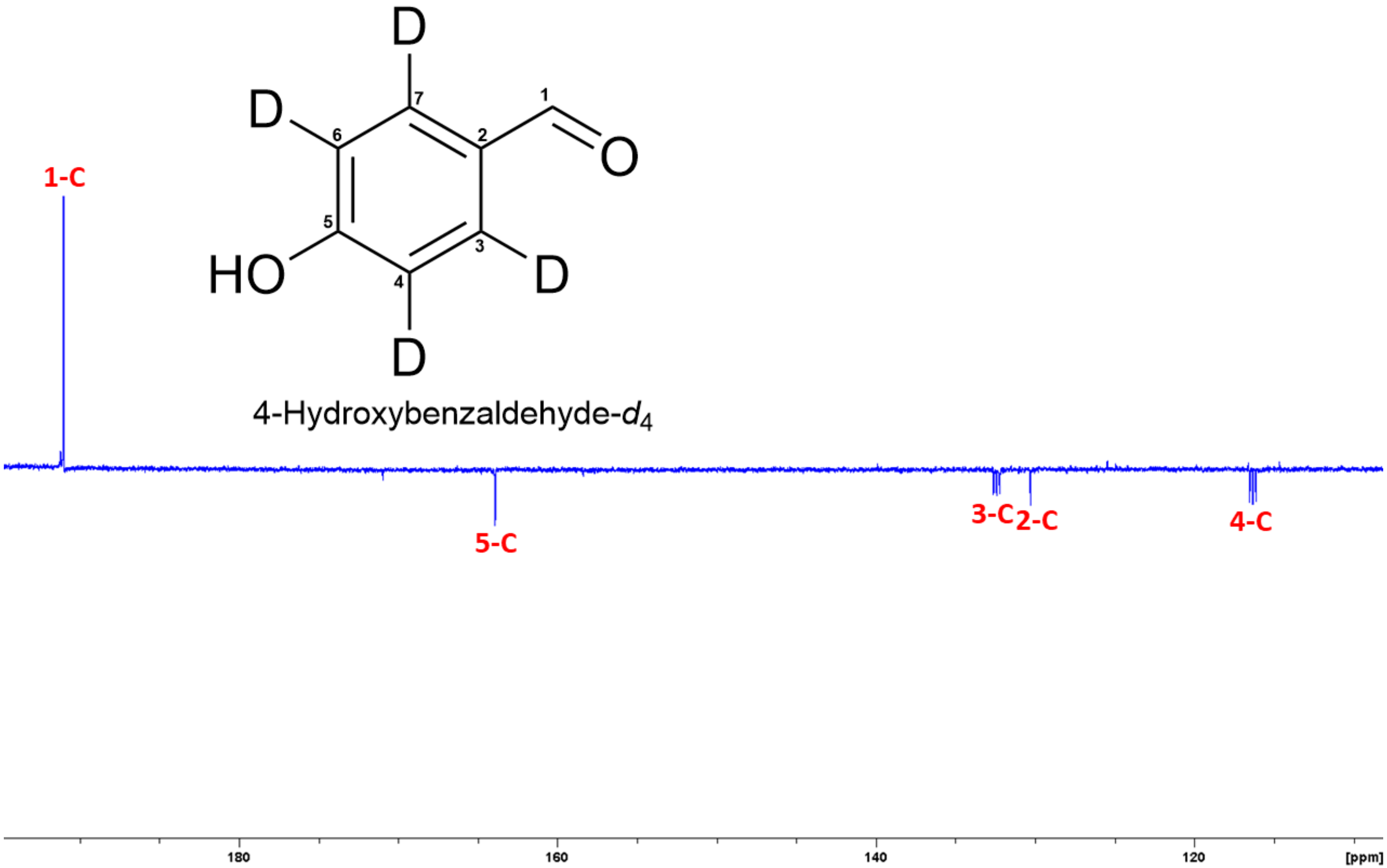


**Figure S37.** DEPTQ spectrum (125 MHz, acetone-$d_6$) of 4-hydroxybenzylalcohol-$d_4$.

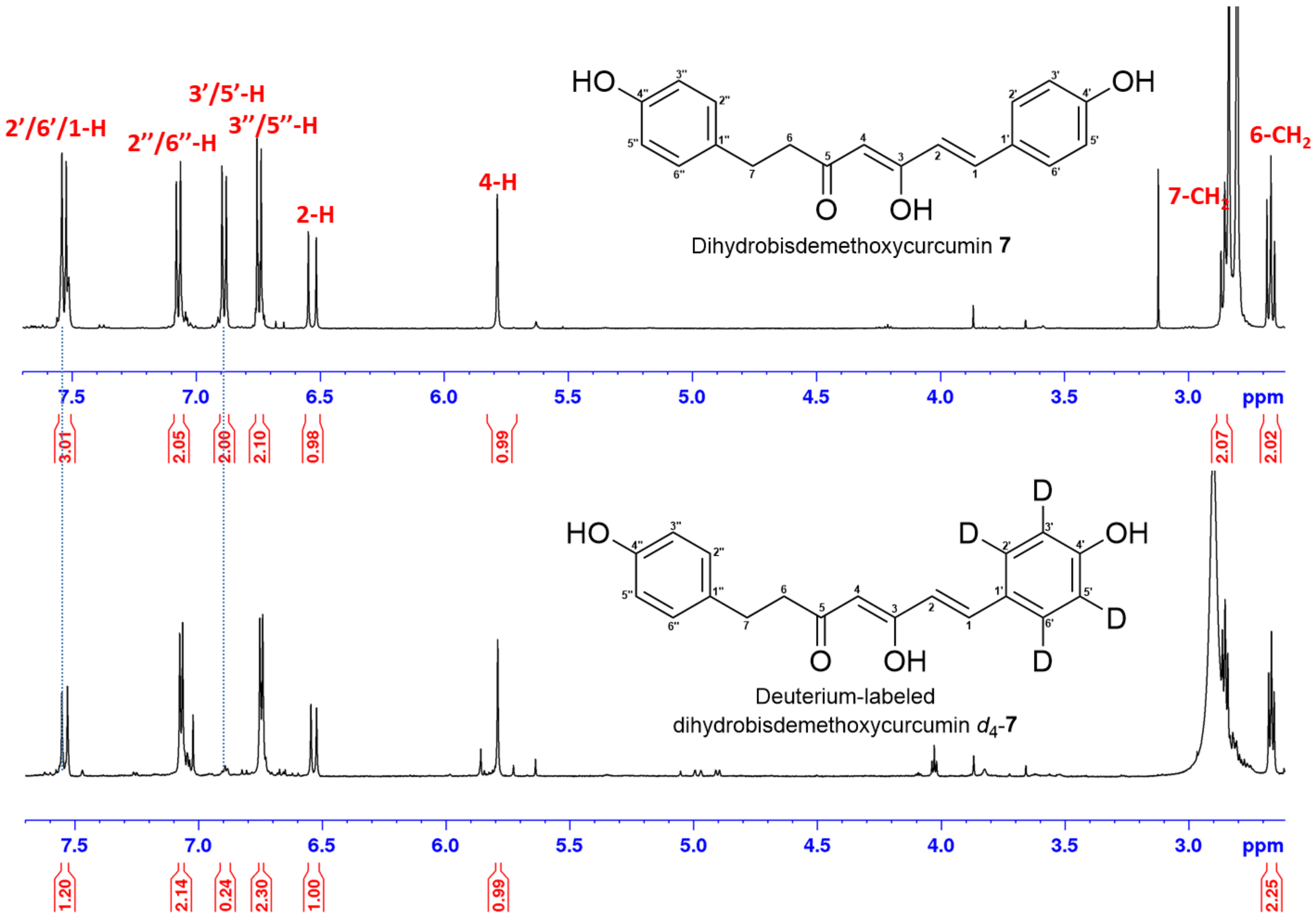


**Figure S38.** Comparison of [1]H NMR spectra of dihydrobisdemethoxycurcumin **7** (upper) and $d_4$-**7** (lower) in acetone-$d_6$.

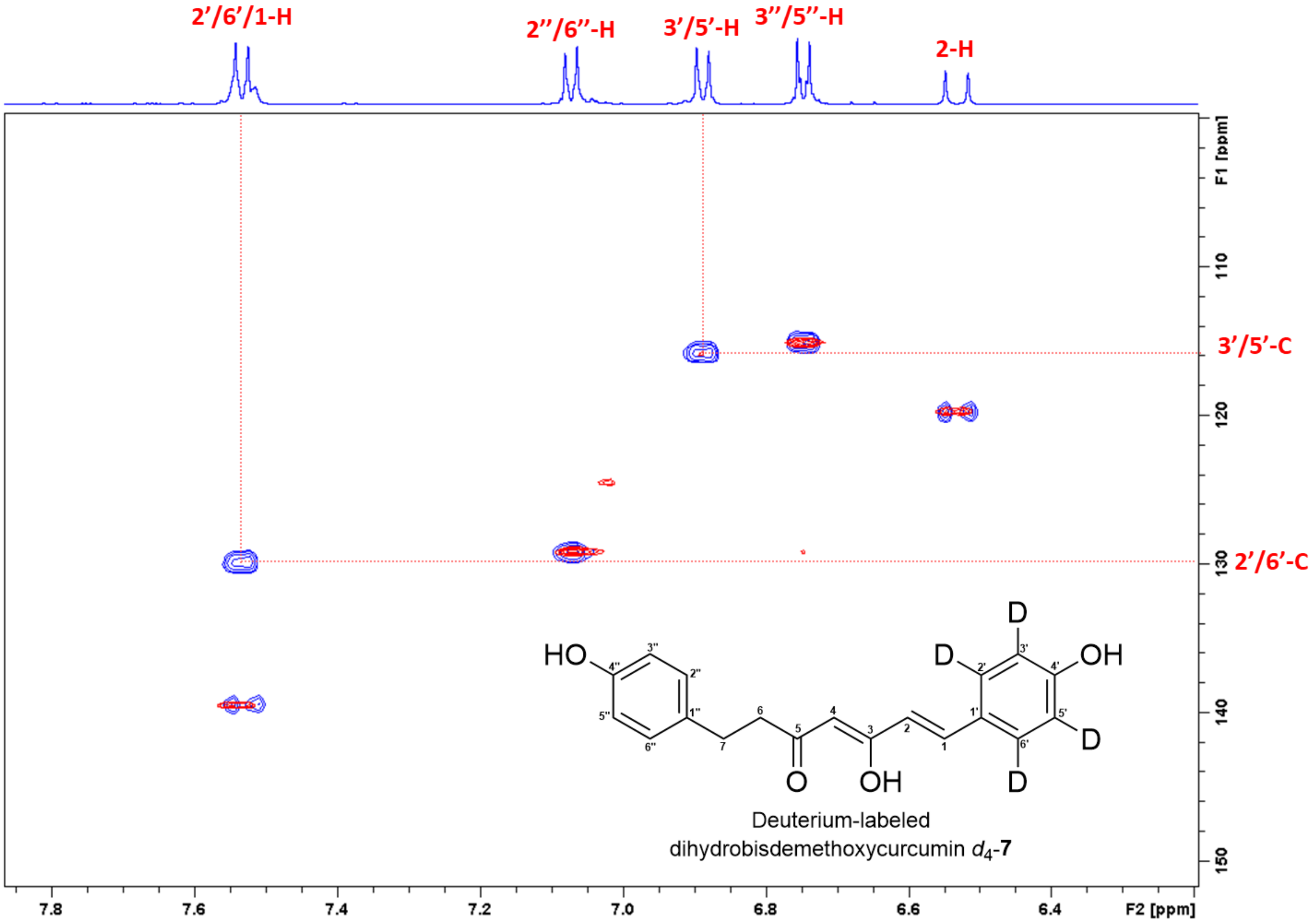


**Figure S39.** Superimposed HSQC spectra of dihydrobisdemethoxycurcumin **7** (blue) and $d_4$-**7** (red) in acetone-$d_6$.

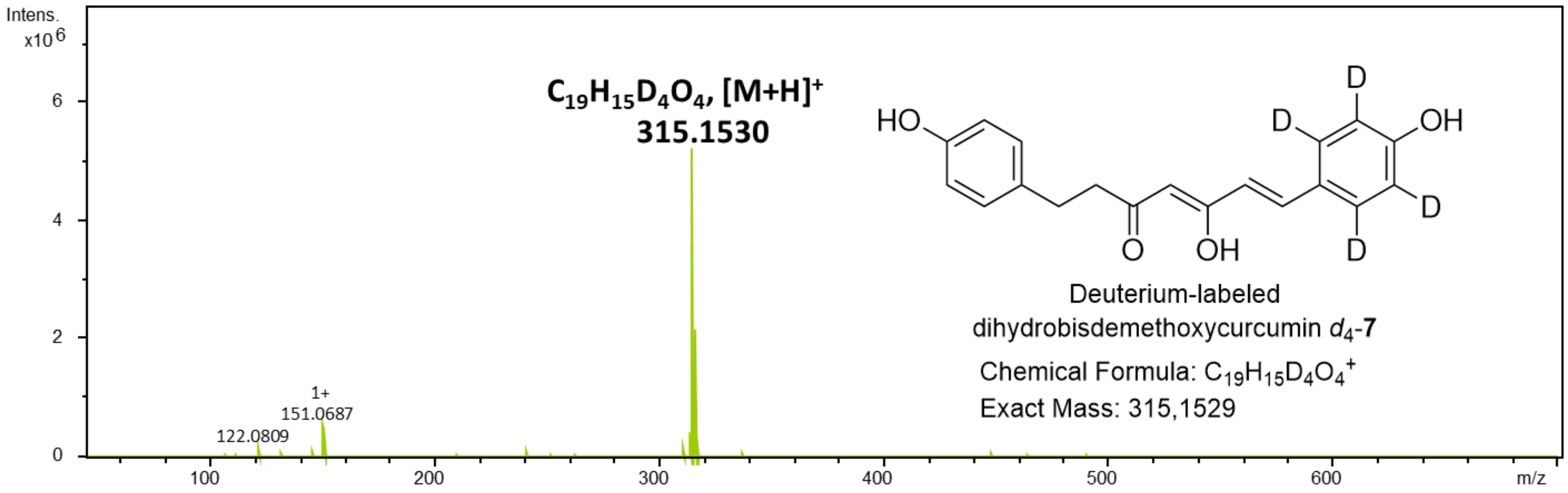


**Figure S40.** HR-ESI-MS spectrum of dihydrobisdemethoxycurcumin $d_4$-**7**.

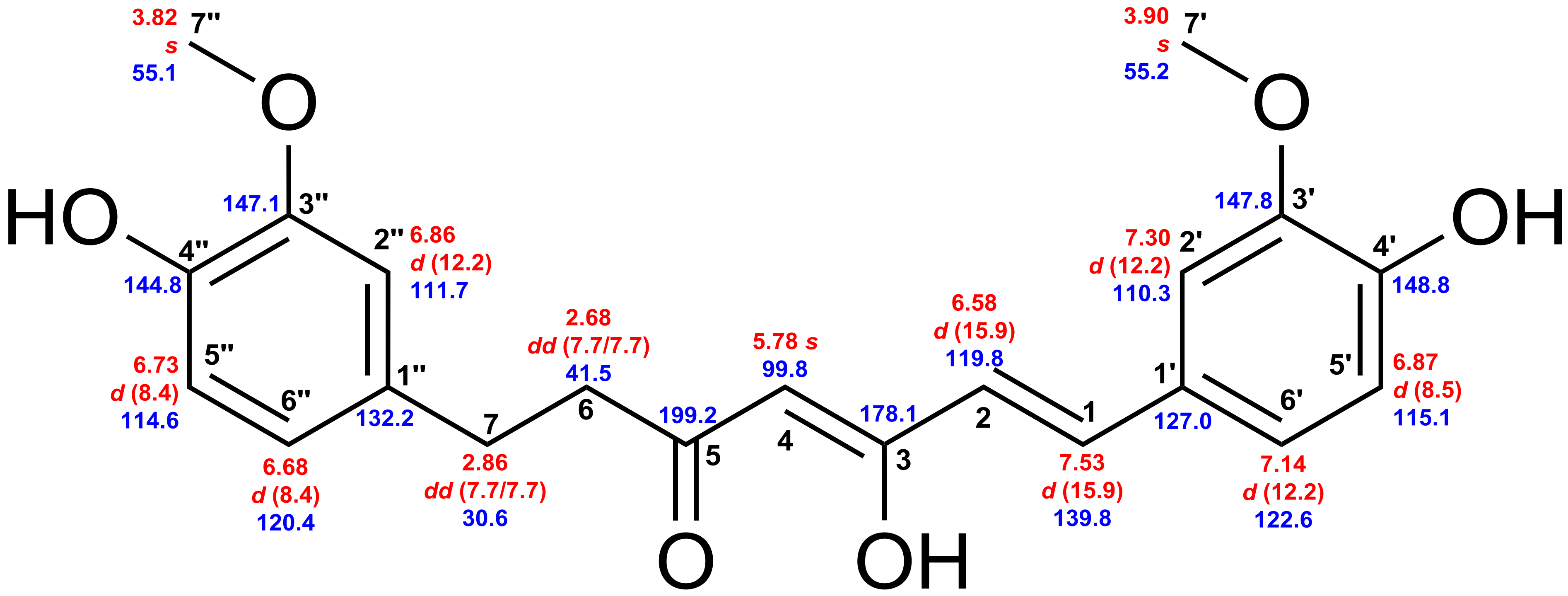


Dihydrocurcumin **8**

**Figure S41.** Chemical shifts of dihydrocurcumin **8**. Red: $^{1}$H chemical shifts (δ ppm, *mult*., $^{3}J_{HH}$ in Hz). Blue: $^{13}$C chemical shifts (δ ppm).

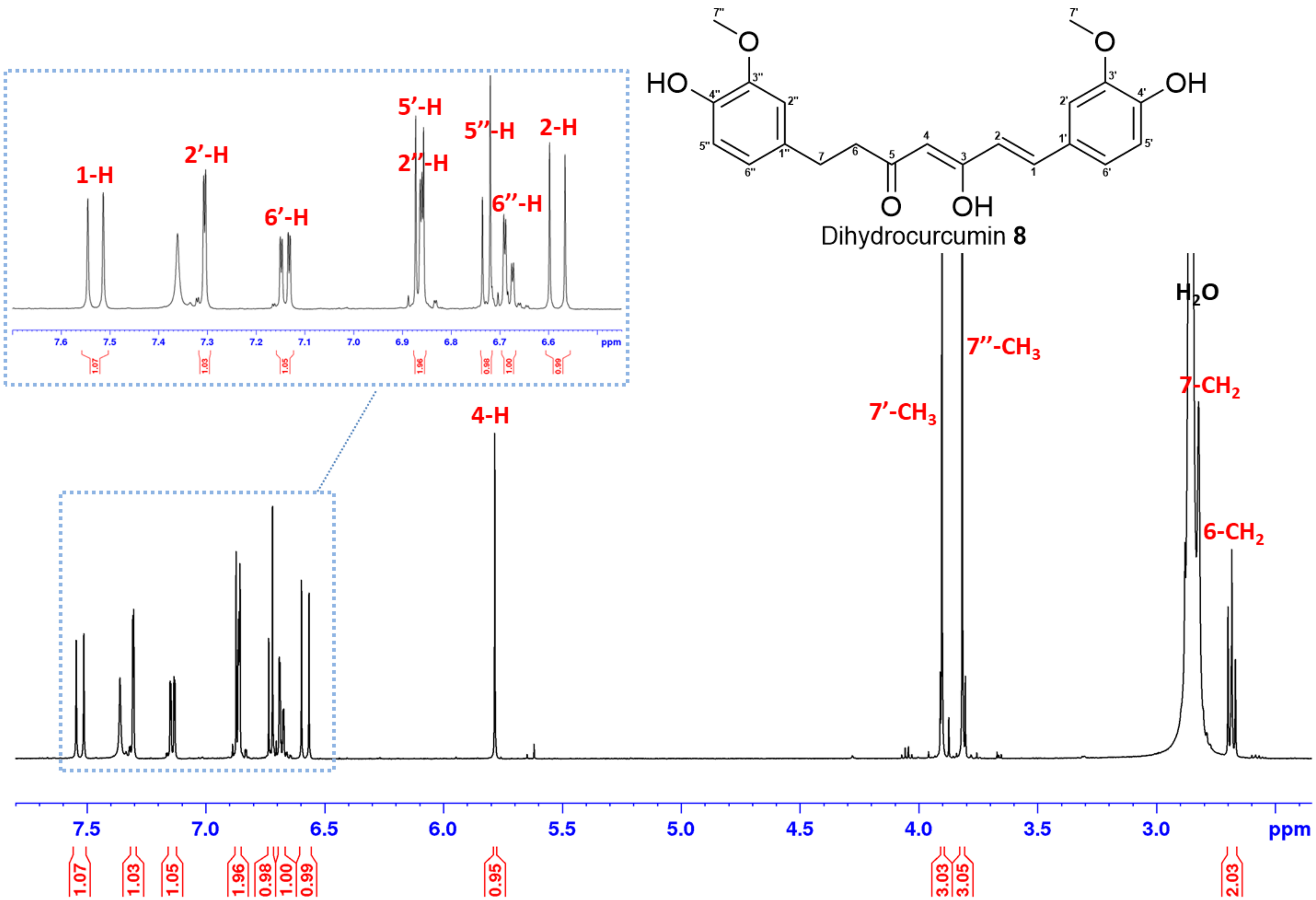


**Figure S42.** $^1$H NMR spectrum (700 MHz, acetone-$d_6$) of dihydrocurcumin **8**.

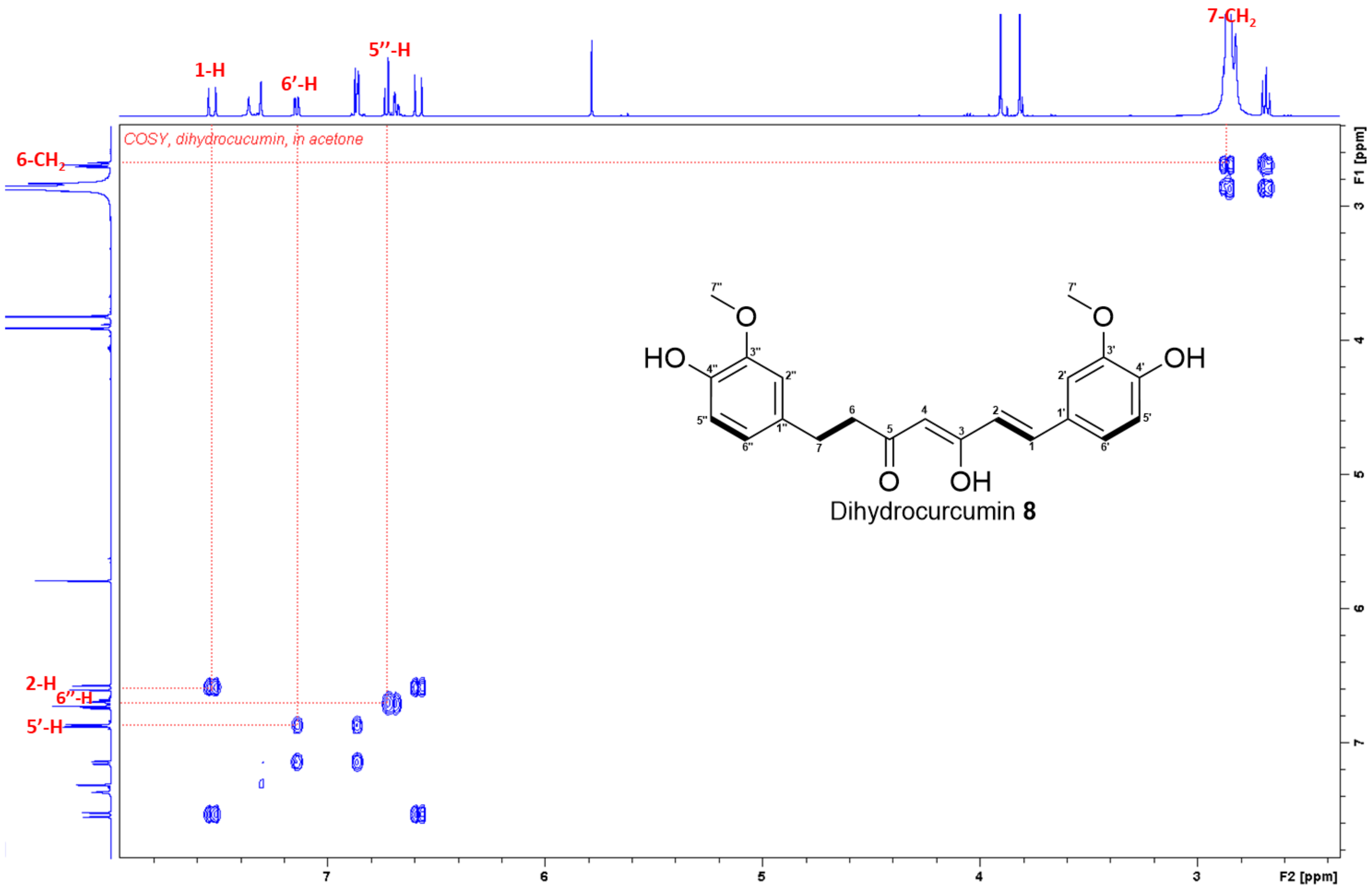


**Figure S43.** $^{1}$H–$^{1}$H COSY spectrum of dihydrocurcumin **8** in acetone-$d_6$.

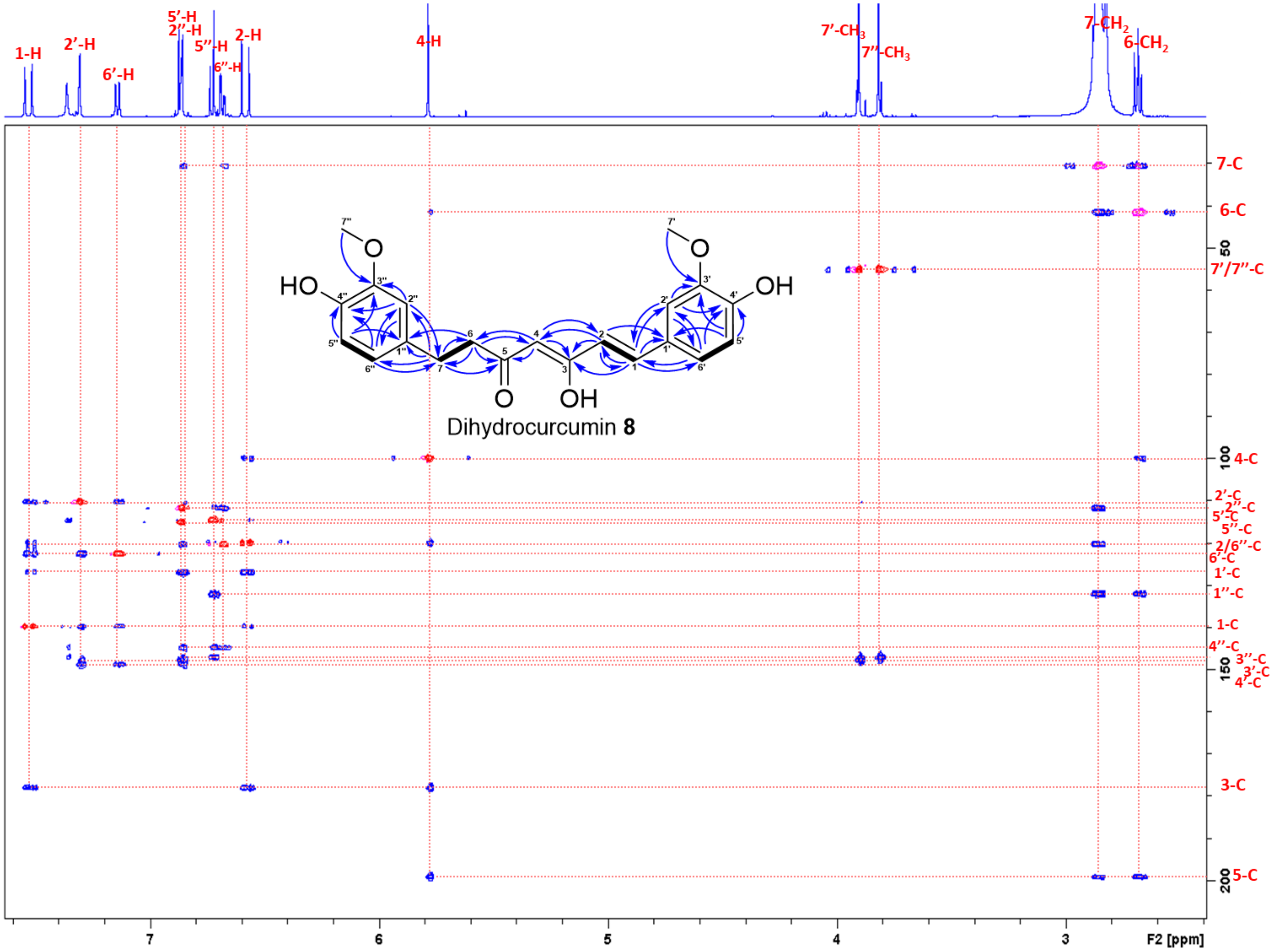

**Figure S44.** Superimposed HSQC and HMBC spectra of dihydrocurcumin **8** in acetone-$d_6$.

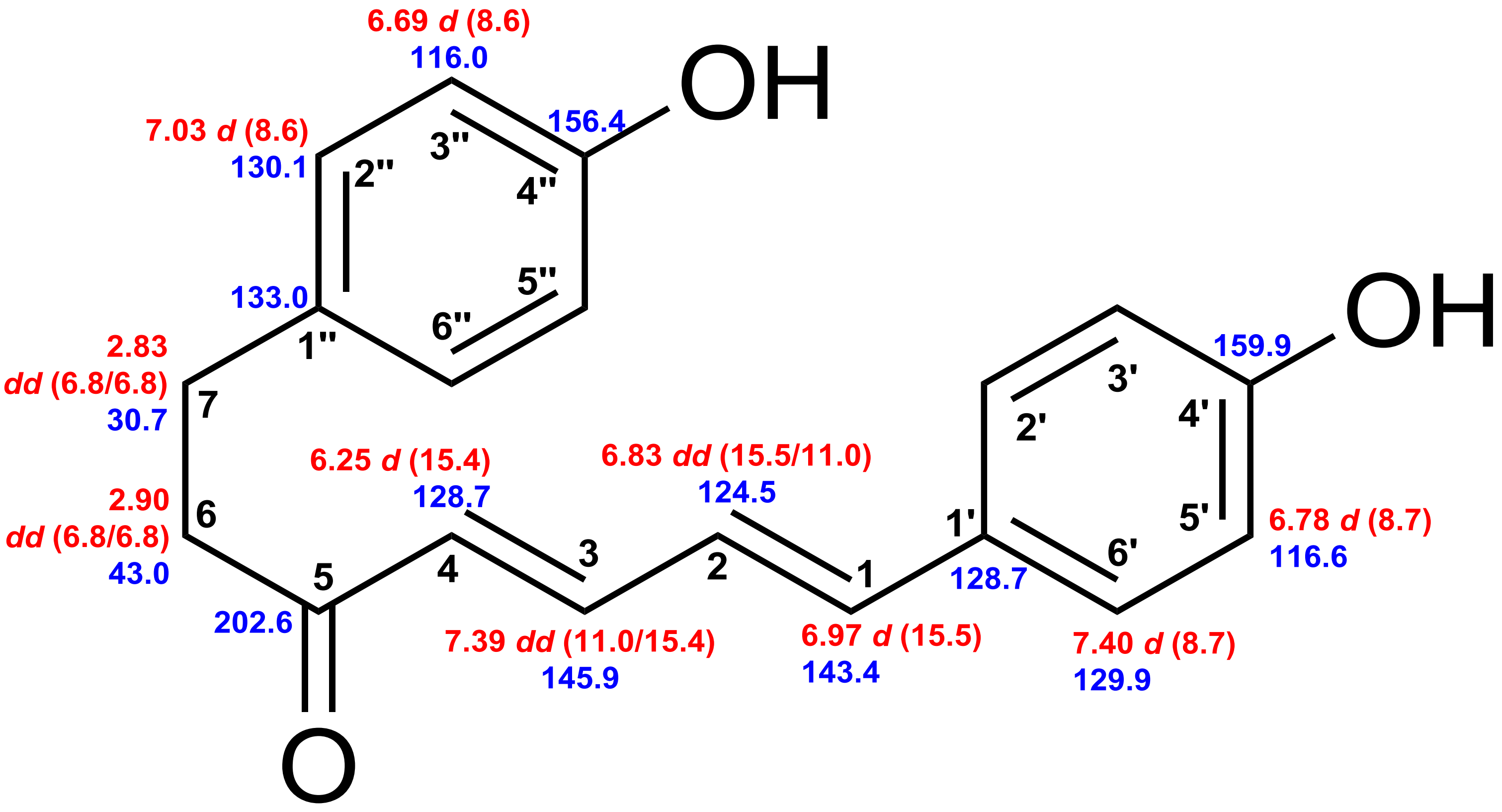


### 4',4''-Dihydroxy linear DH **11**

**Figure S45.** Chemical shifts of 4',4''-dihydroxy linear DH **11**. Red: $^{1}$H chemical shifts (δ ppm, *mult*., $^{3}J_{HH}$ in Hz). Blue: $^{13}$C chemical shifts (δ ppm).

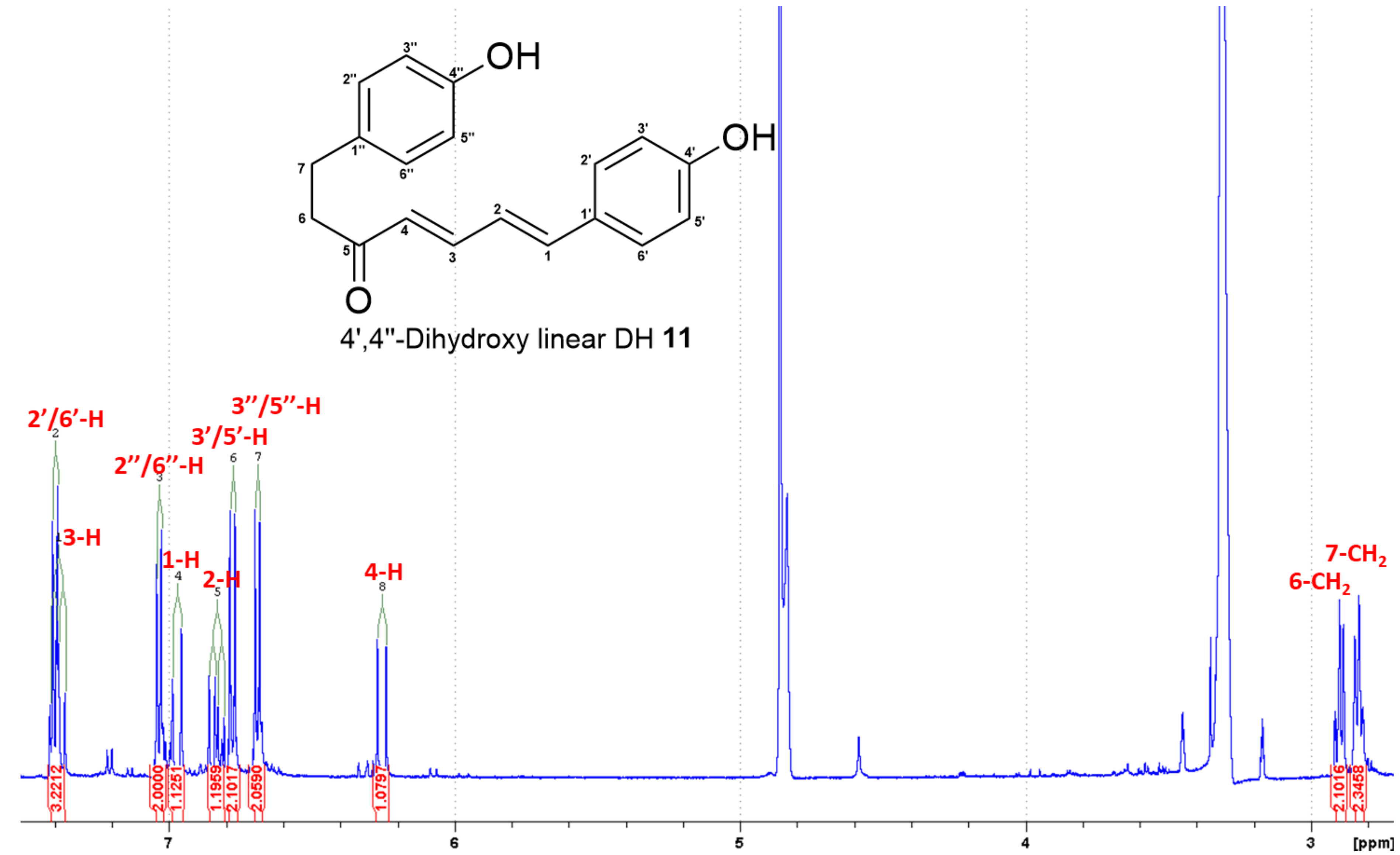


**Figure S46.** $^{1}$H NMR spectrum (700 MHz, $CD_3OD$) of 4',4''-dihydroxy linear DH **11**.

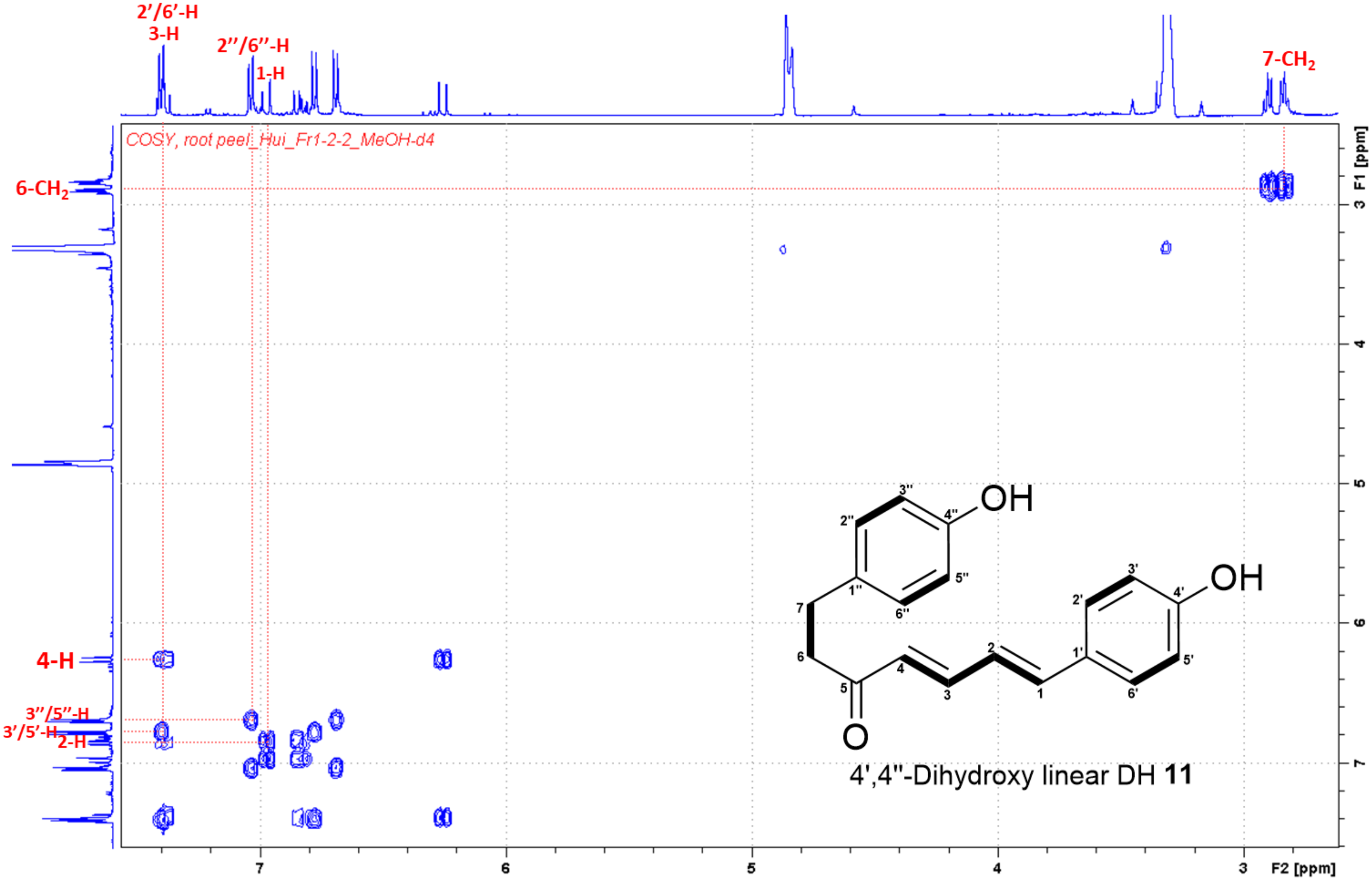


**Figure S47.** $^{1}H$–$^{1}H$ COSY spectrum of 4',4''-dihydroxy linear DH **11** in $CD_3OD$.

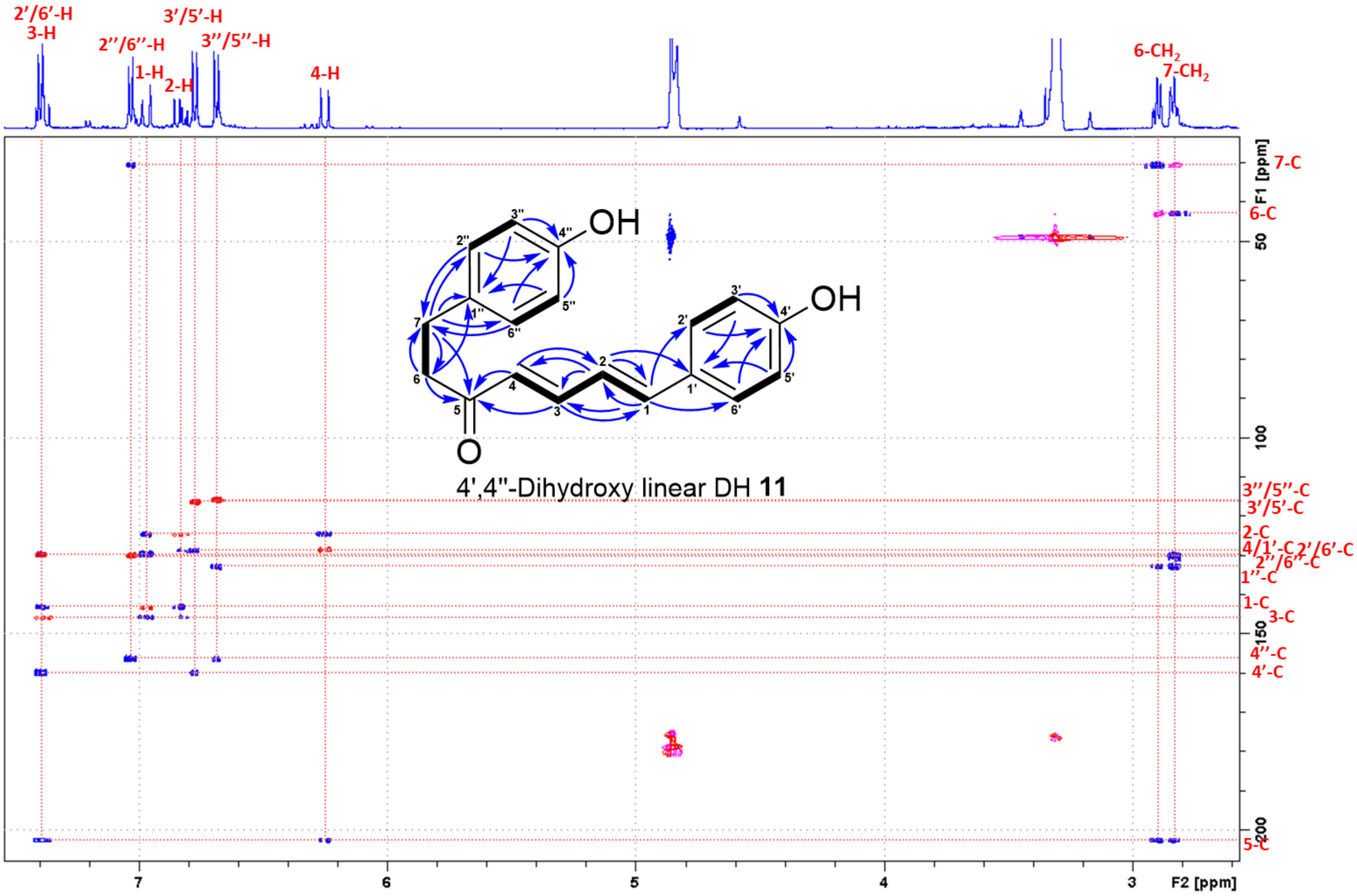


**Figure S48.** Superimposed HSQC and HMBC spectra of 4',4''-dihydroxy linear DH **11** in $CD_3OD$.

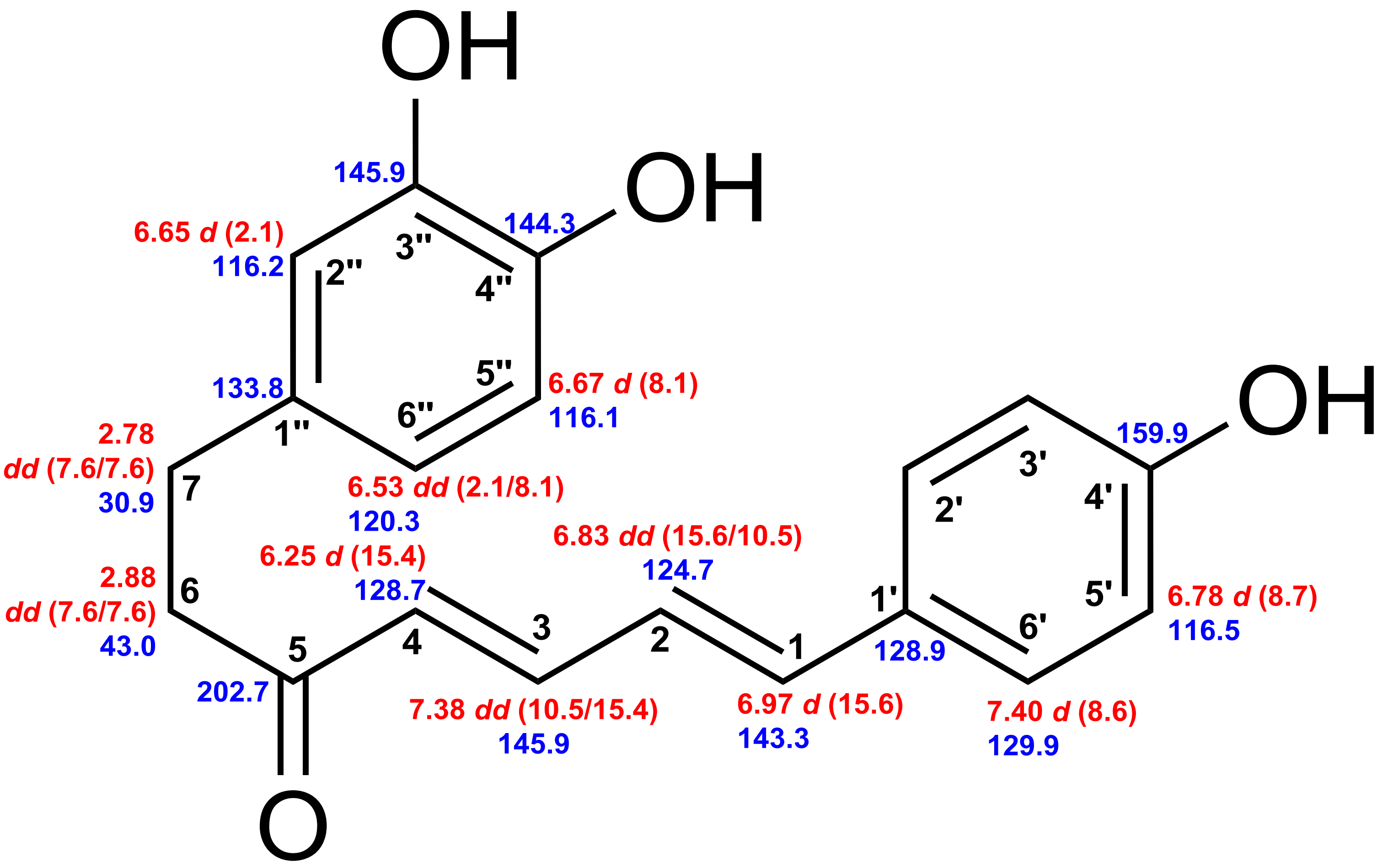


4',3'',4''-Trihydroxy linear DH **12**

**Figure S49.** Chemical shifts of 4',3'',4''-trihydroxy linear DH **12**. Red: $^{1}$H chemical shifts (δ ppm, *mult*., $^{3}J_{HH}$ in Hz). Blue: $^{13}$C chemical shifts (δ ppm).

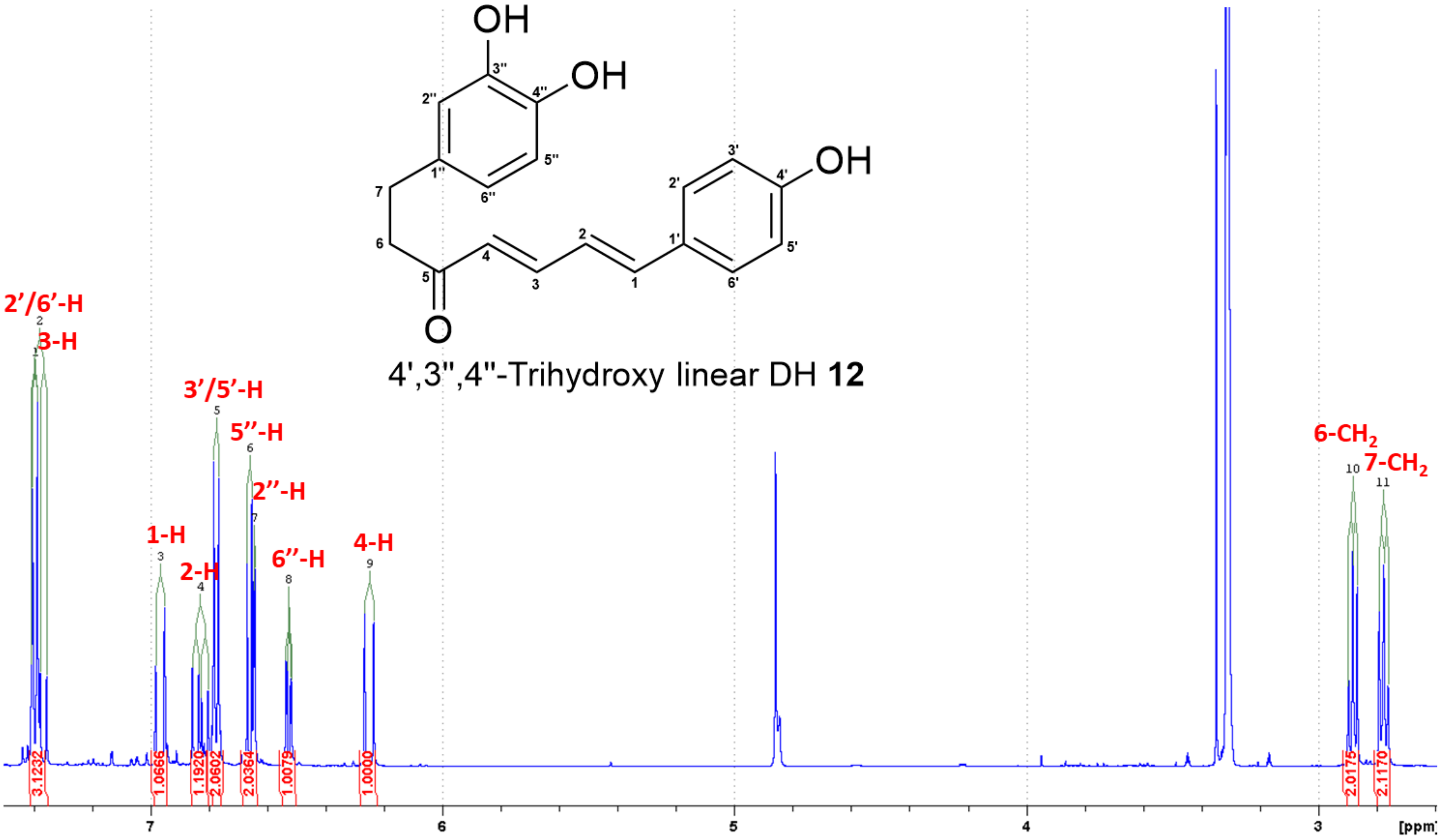


**Figure S50.** $^{1}H$ NMR spectrum (700 MHz, $CD_3OD$) of 4',3'',4''-trihydroxy linear DH **12**.

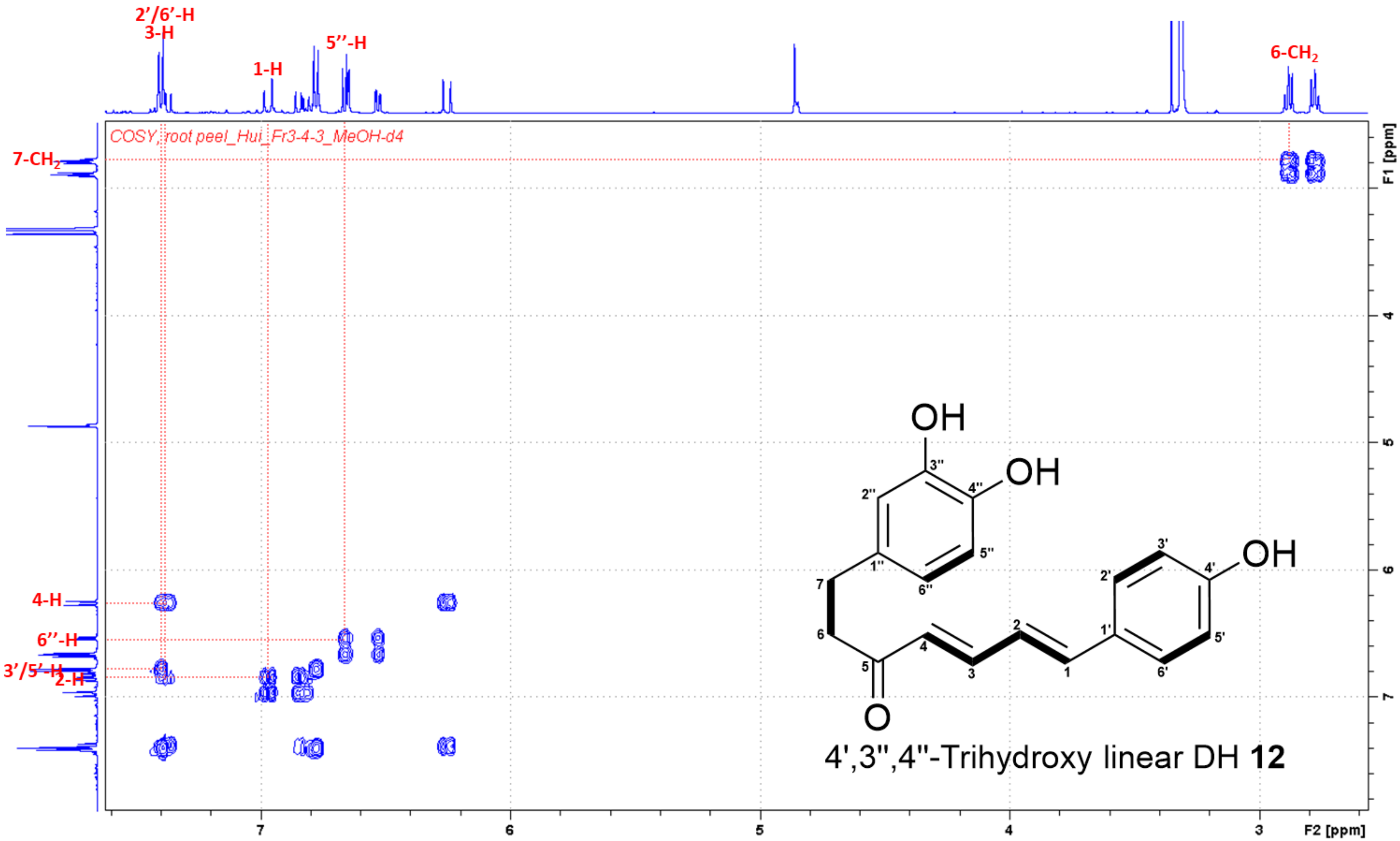


**Figure S51.** $^1H$–$^1H$ COSY spectrum of 4',3'',4''-trihydroxy linear DH **12** in $CD_3OD$.

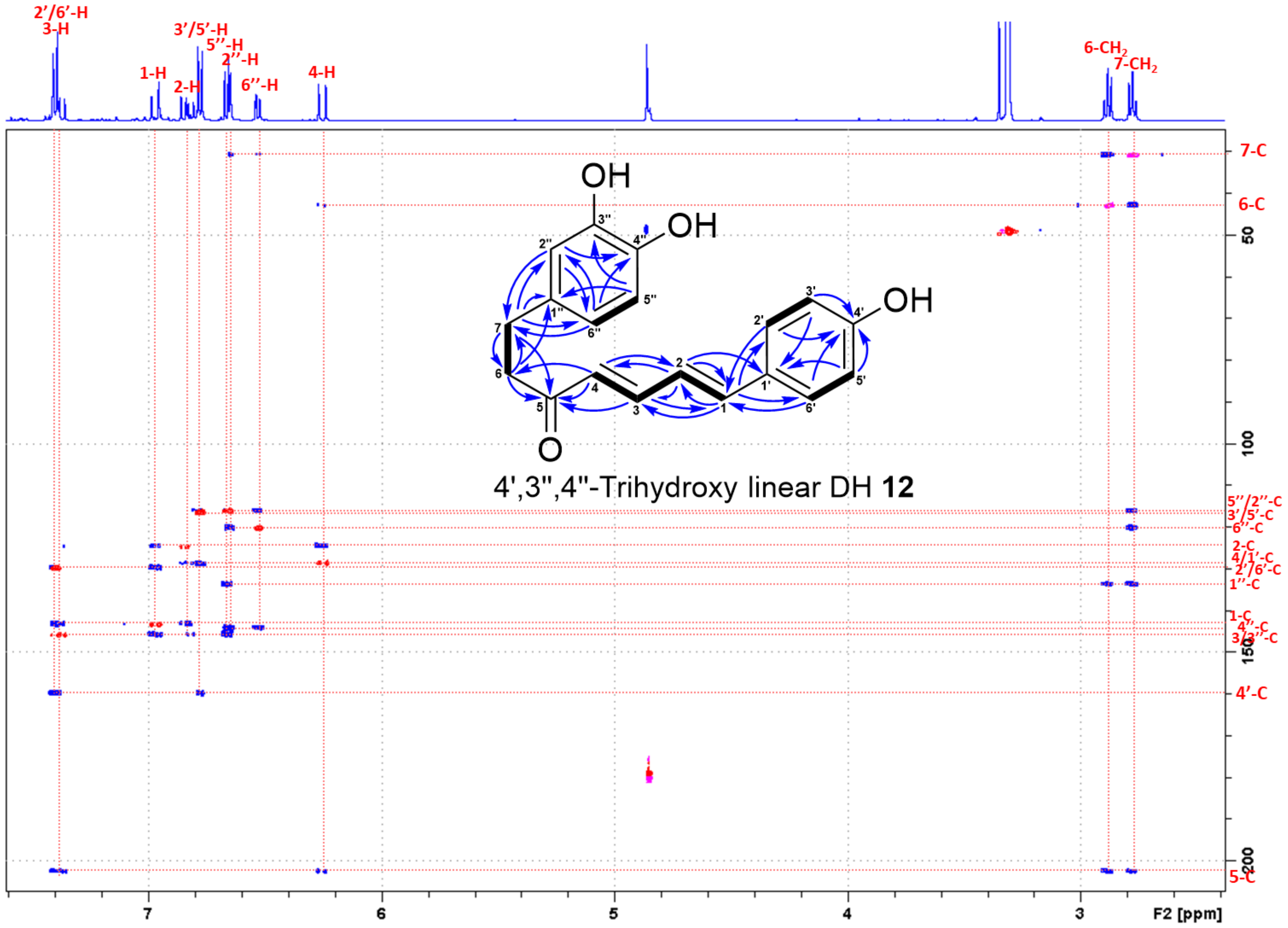


**Figure S52.** Superimposed HSQC and HMBC spectra of 4',3",4"-trihydroxy linear DH **12** in $CD_3OD$.

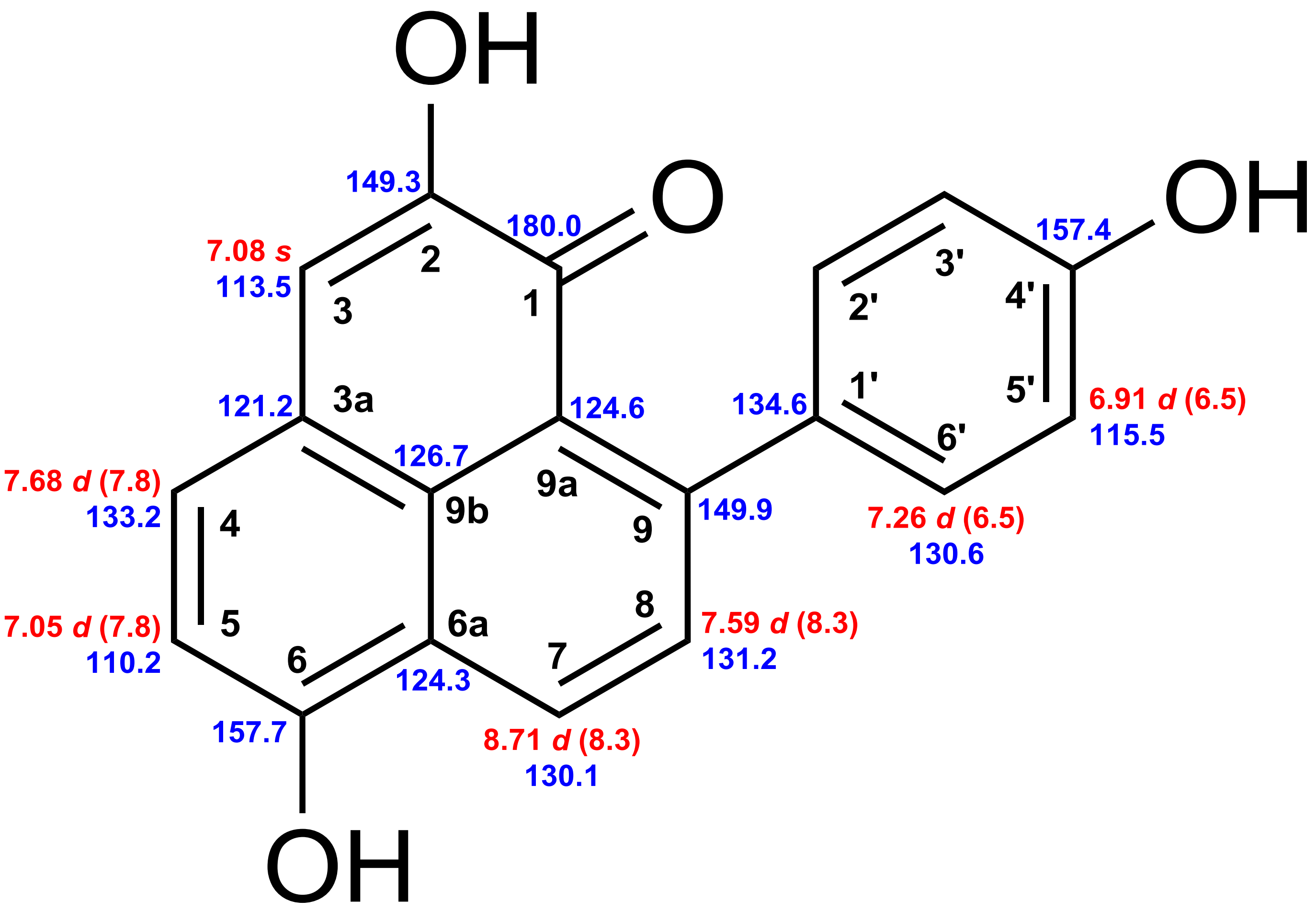


4'-Hydroxylachnanthocarpone **14**

**Figure S53.** Chemical shifts of 4'-hydroxylachnanthocarpone **14**. Red: $^{1}$H chemical shifts (δ ppm, *mult.*, $^{3}J_{HH}$ in Hz). Blue: $^{13}$C chemical shifts (δ ppm).

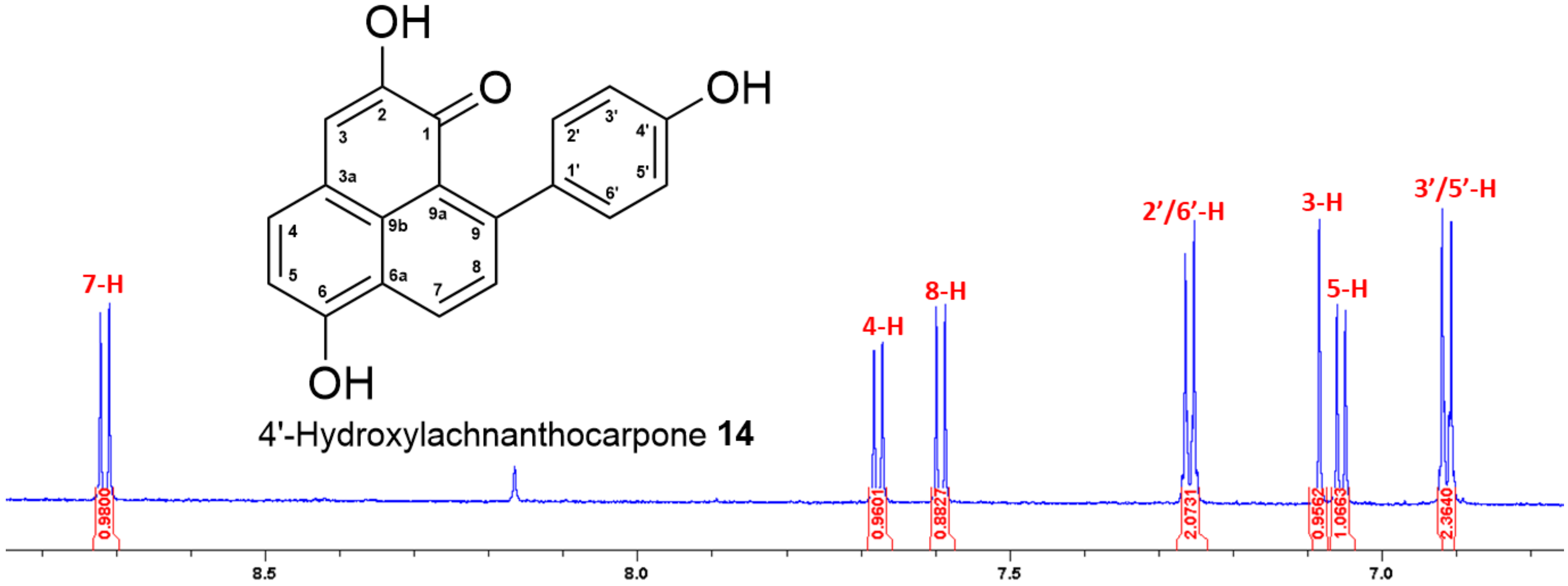


**Figure S54.** $^{1}$H NMR spectrum (700 MHz, acetone-$d_6$) of 4'-hydroxylachnanthocarpone **14**.

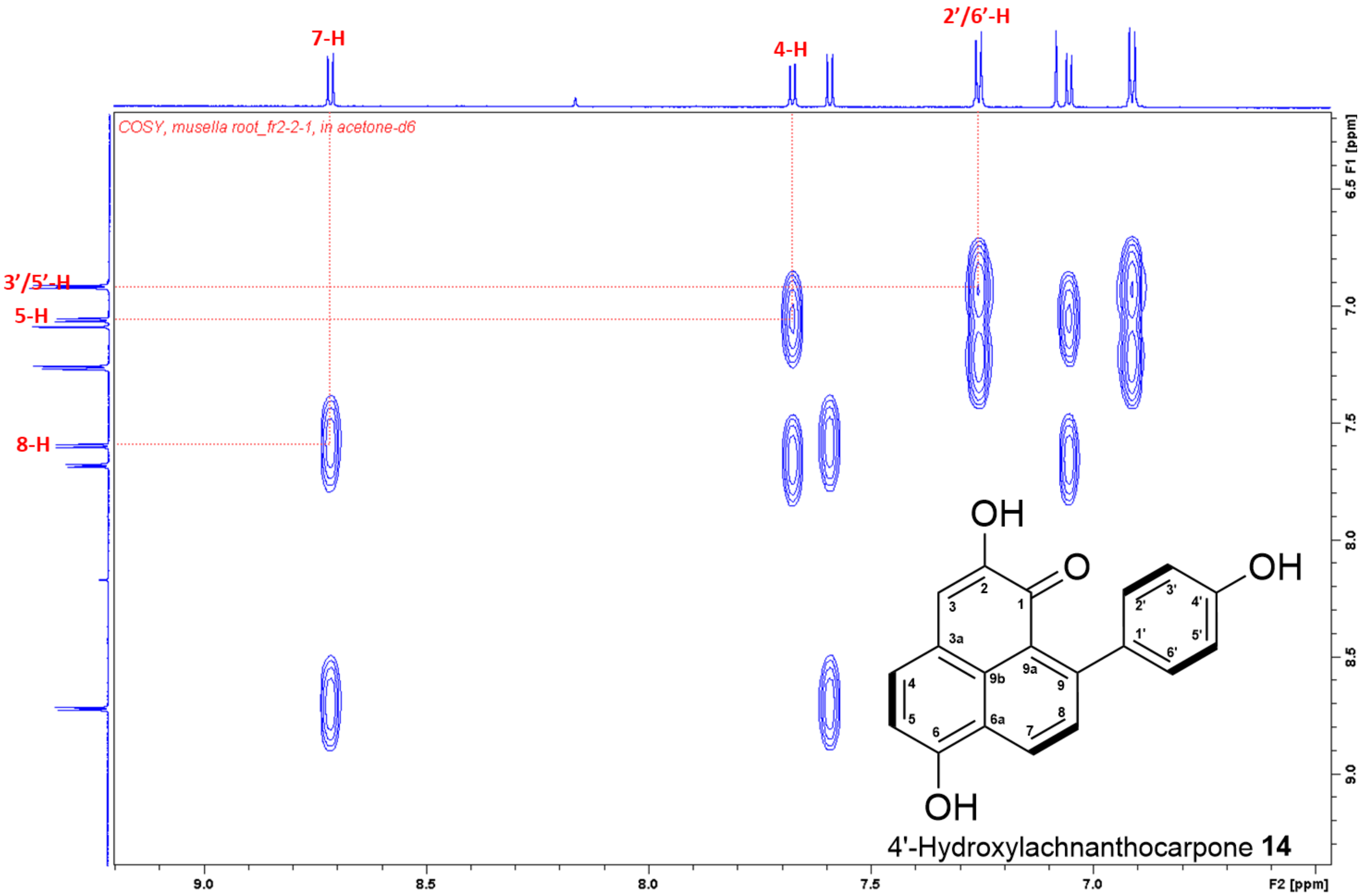


**Figure S55.** $^{1}H$–$^{1}H$ COSY spectrum of 4'-hydroxylachnanthocarpone **14** in acetone-$d_6$.

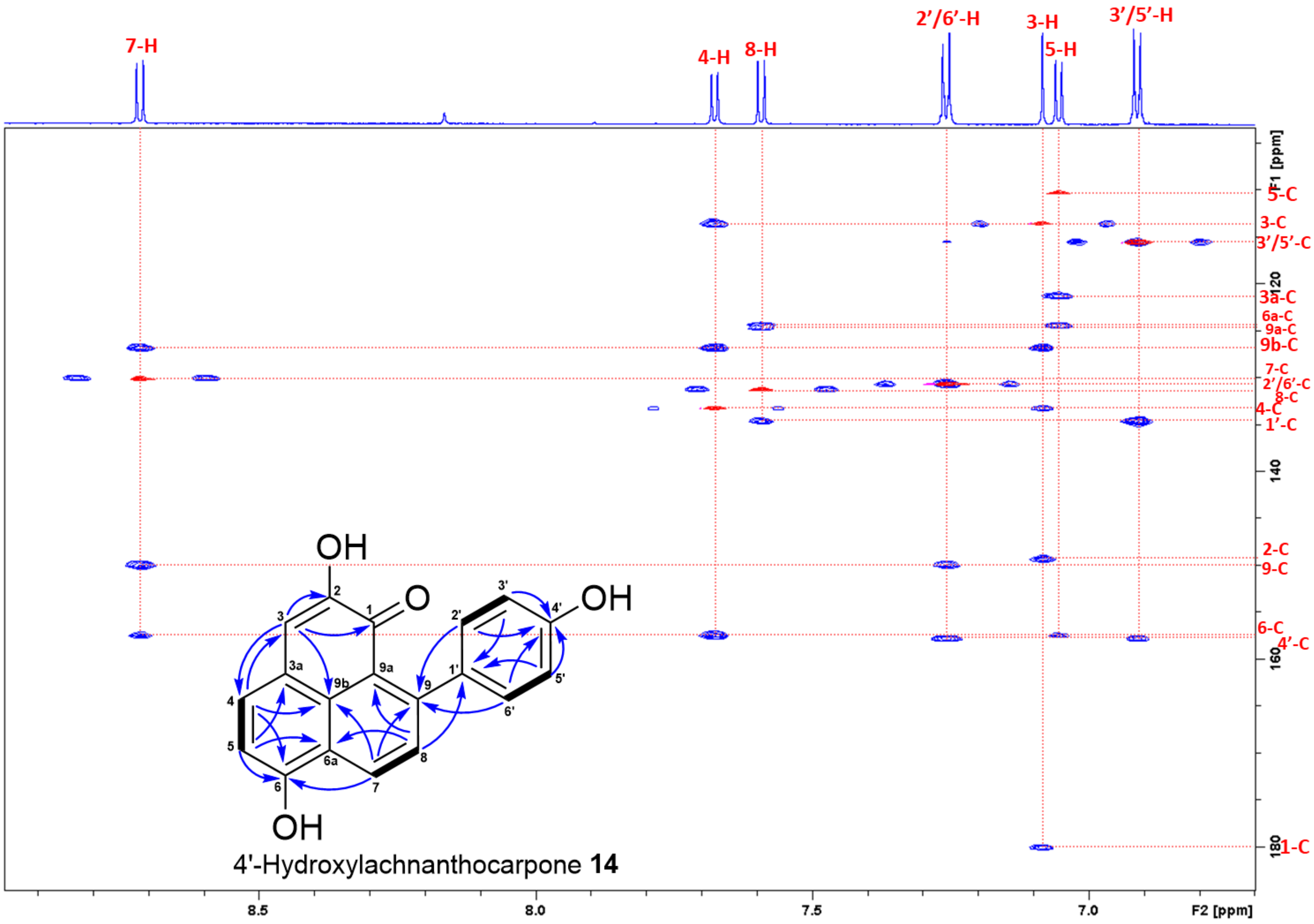


**Figure S56.** Superimposed HSQC and HMBC spectra of 4'-hydroxylachnanthocarpone **14** in acetone-d6.

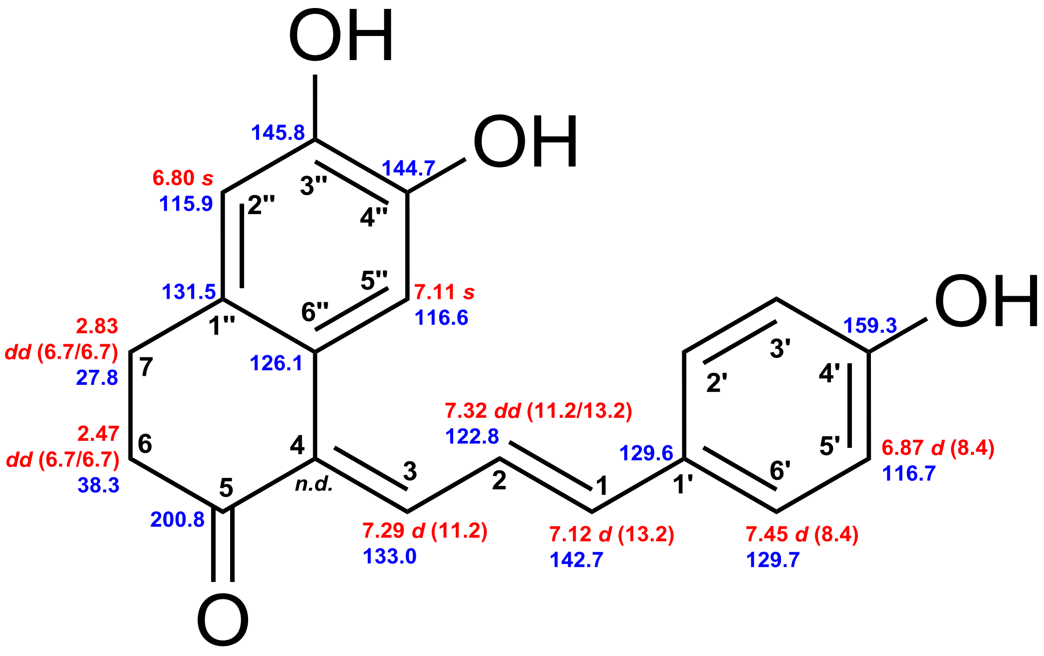


Monocyclic DH **13**

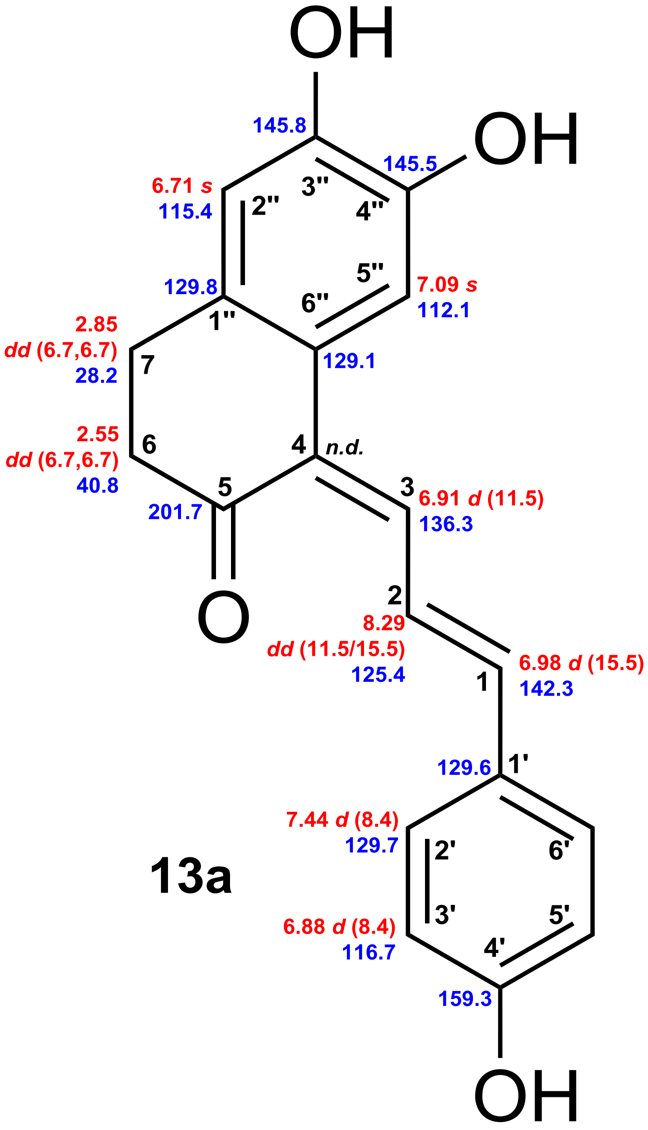


**13a**

**Figure S57.** Chemical shifts of monocyclic DH **13** and **13a** (*n.d.* not detected). Red: $^{1}$H chemical shifts (δ ppm, *mult*., $^{3}J_{HH}$ in Hz). Blue: $^{13}$C chemical shifts (δ ppm).

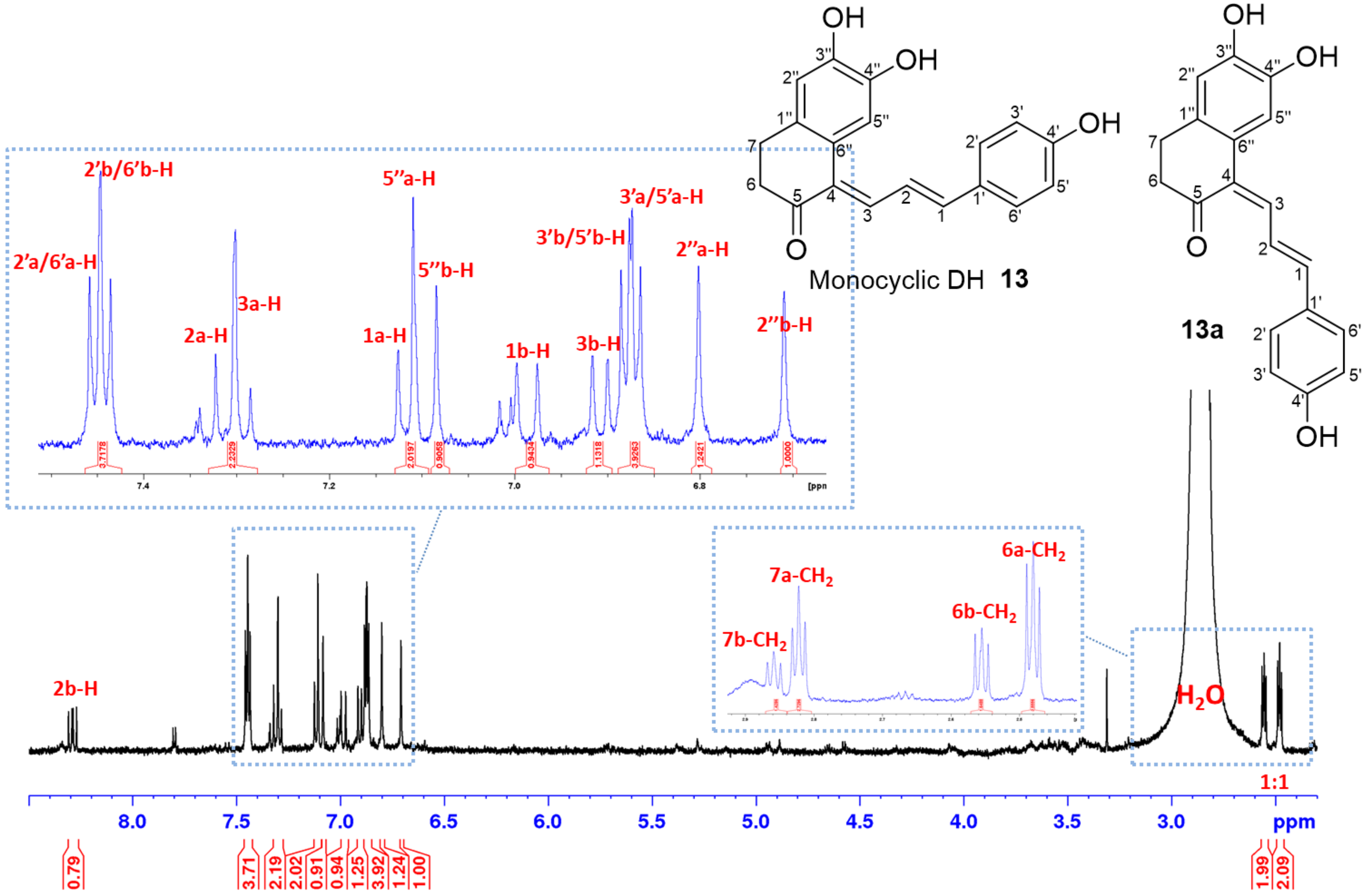


**Figure S58.** [1]H NMR spectrum (700 MHz, acetone-$d_6$) of monocyclic DH **13** (assigned as a) and **13a** (assigned as b).

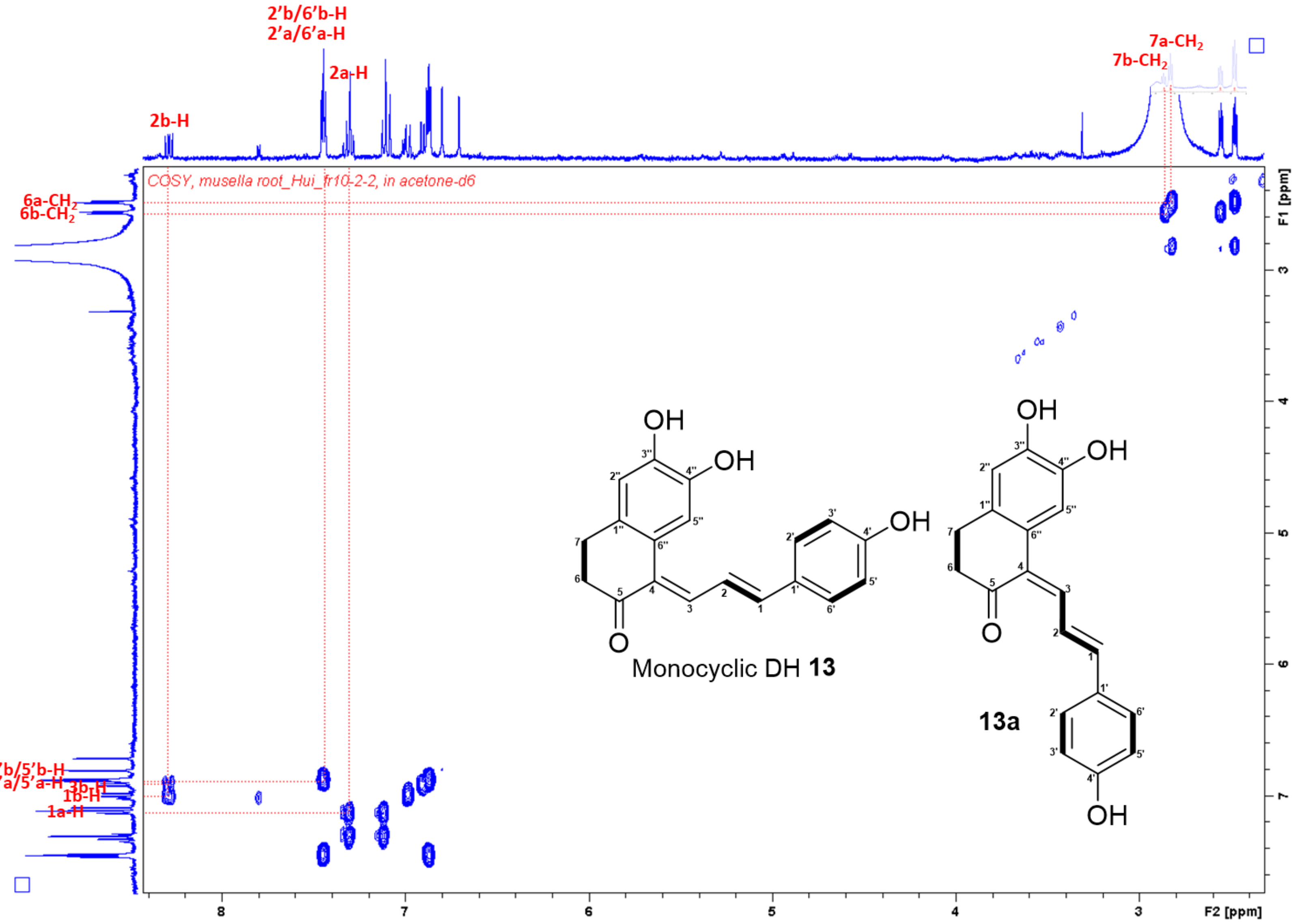


**Figure S59.** $^1$H–$^1$H COSY spectrum of monocyclic DH **13** (assigned as a) and **13a** (assigned as b) in acetone-$d_6$.

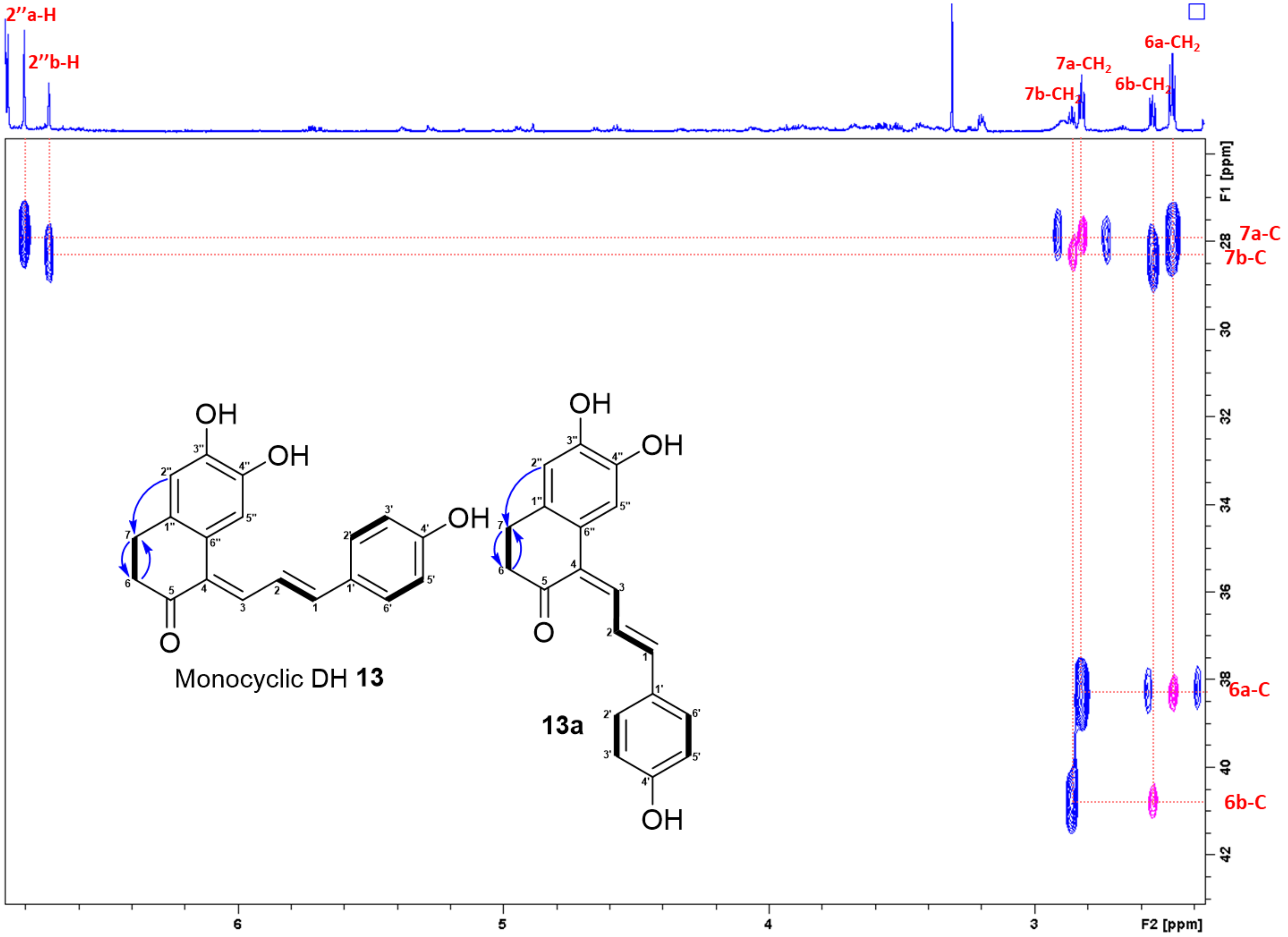


**Figure S60.** Superimposed HSQC and HMBC spectra of monocyclic DH **13** (assigned as a) and **13a** (assigned as b) in acetone-$d_6$ (part-1).

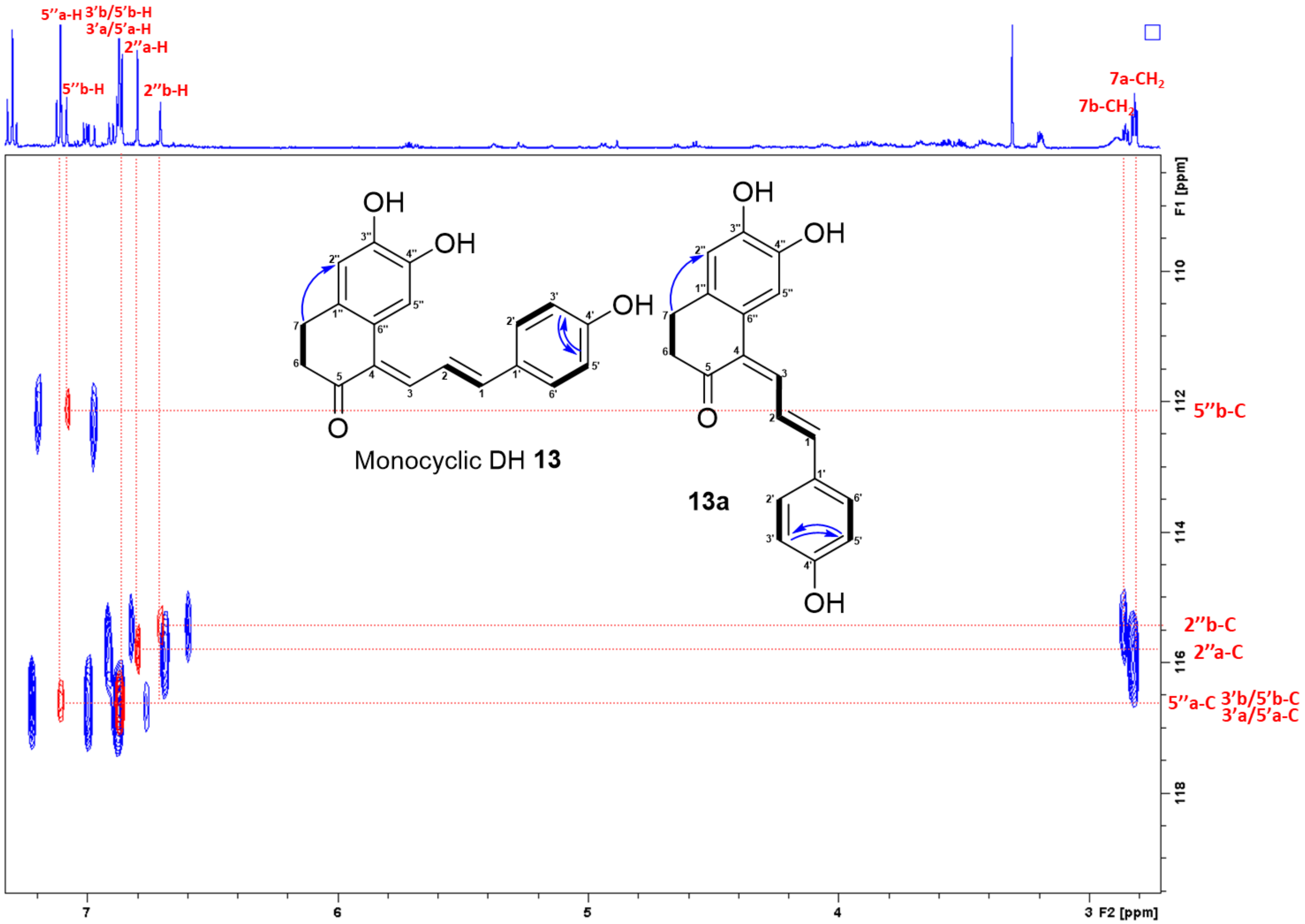


**Figure S61.** Superimposed HSQC and HMBC spectra of monocyclic DH **13** (assigned as a) and **13a** (assigned as b) in acetone-$d_6$ (part-2).

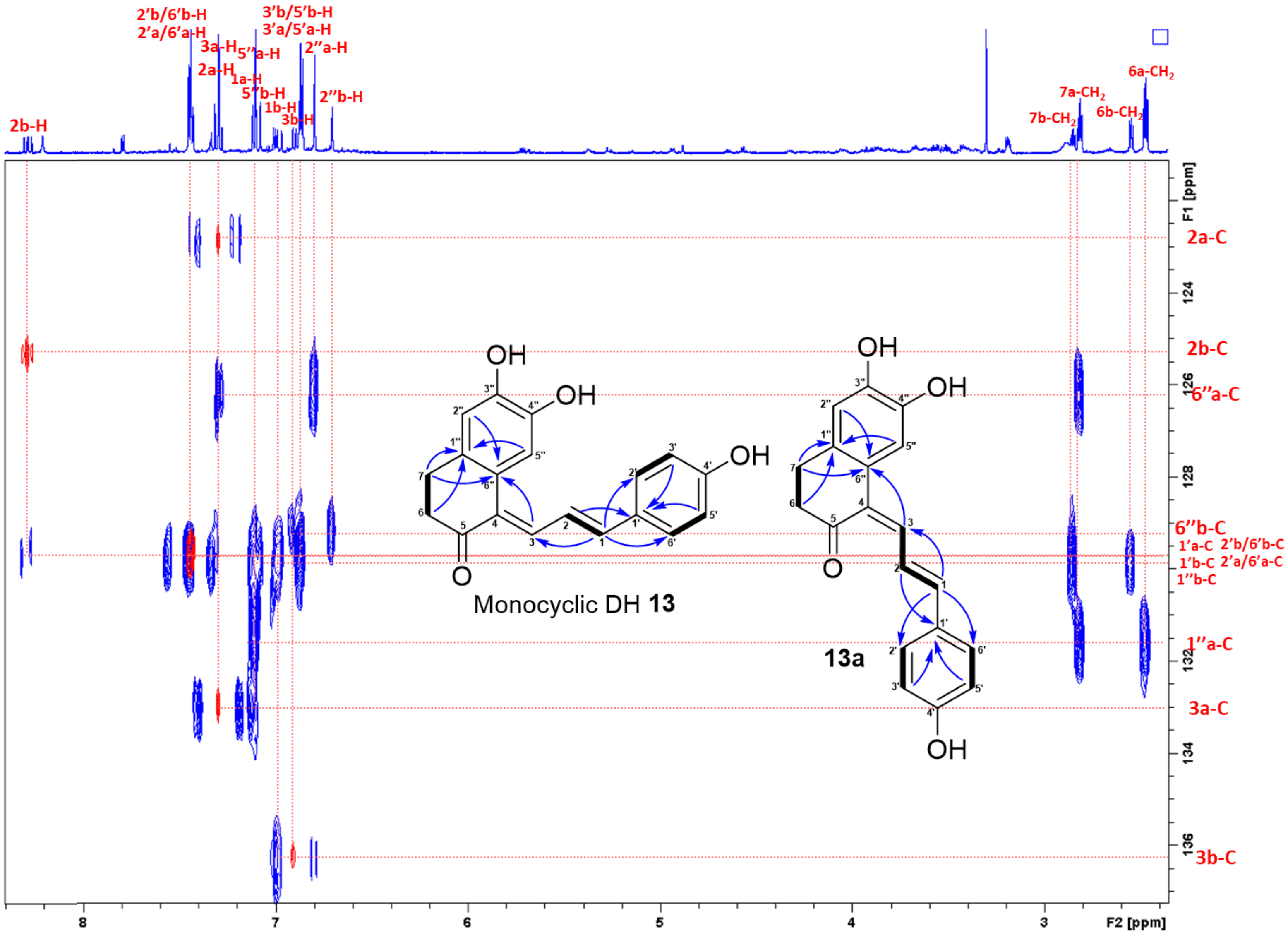


**Figure S62.** Superimposed HSQC and HMBC spectra of monocyclic DH **13** (assigned as a) and **13a** (assigned as b) in acetone-$d_6$ (part-3).

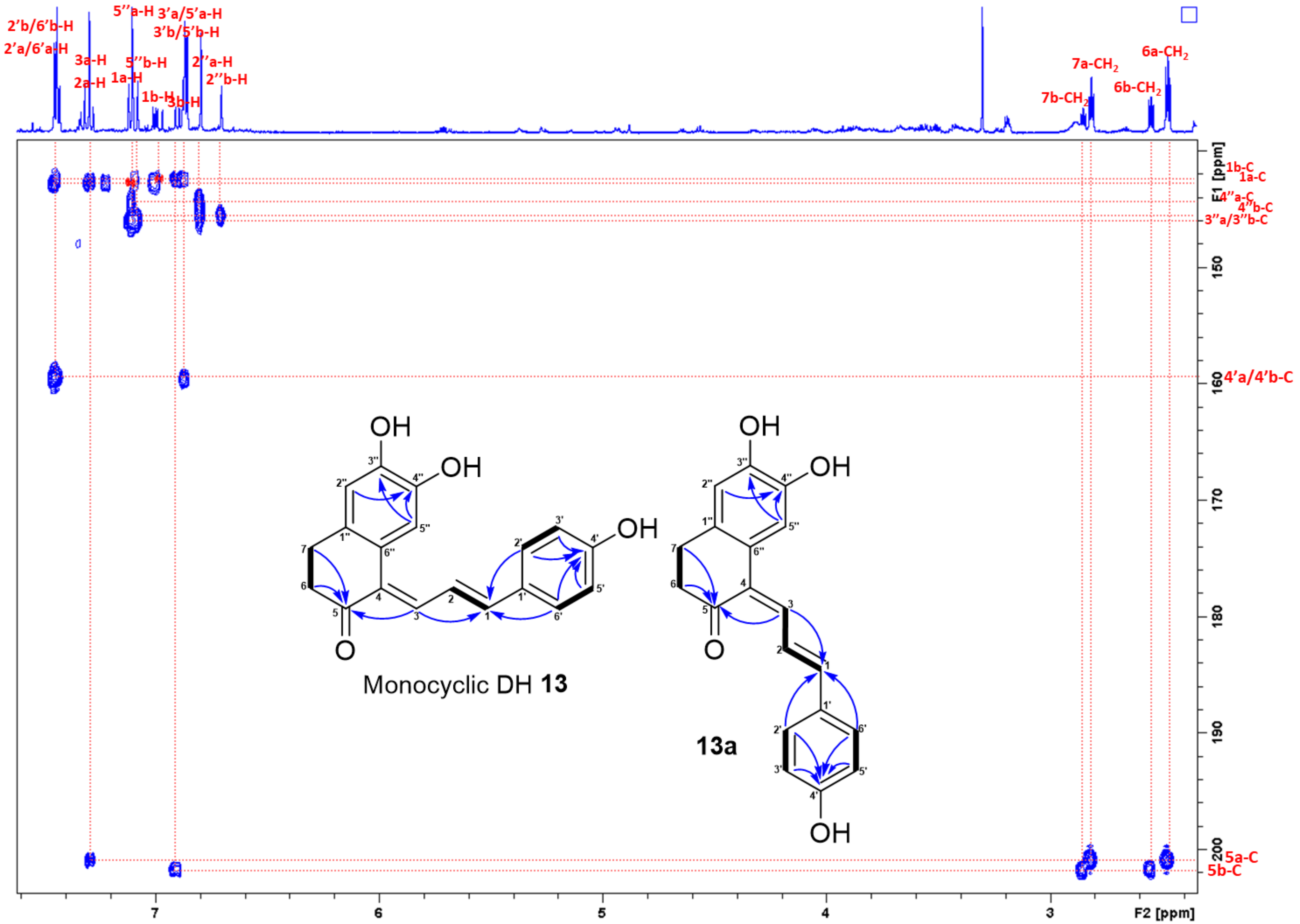


**Figure S63.** Superimposed HSQC and HMBC spectra of monocyclic DH **13** (assigned as a) and **13a** (assigned as b) in acetone-$d_6$ (part-4).

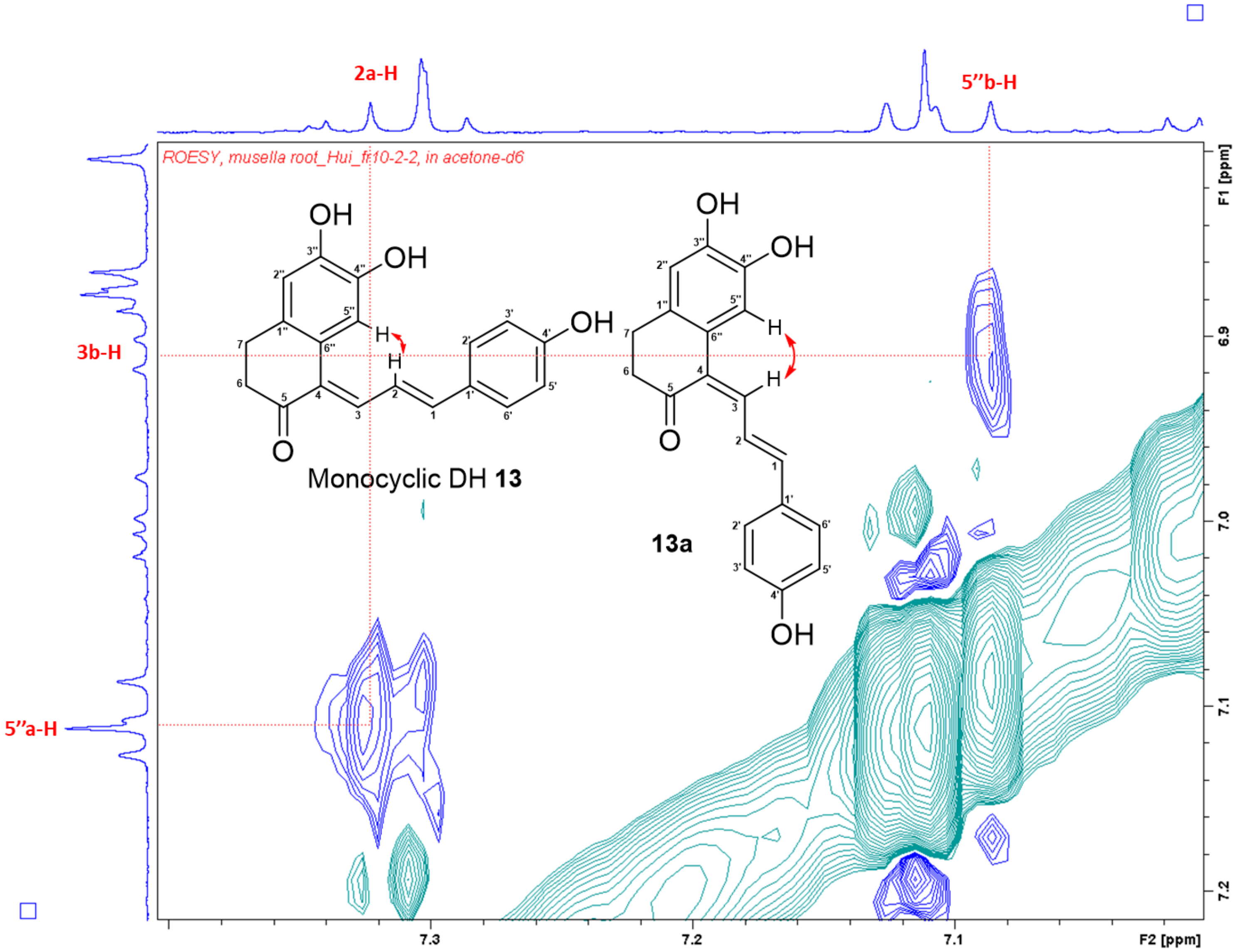


**Figure S64.** ROESY spectrum of monocyclic DH **13** (assigned as a) and **13a** (assigned as b) in acetone-$d_6$.

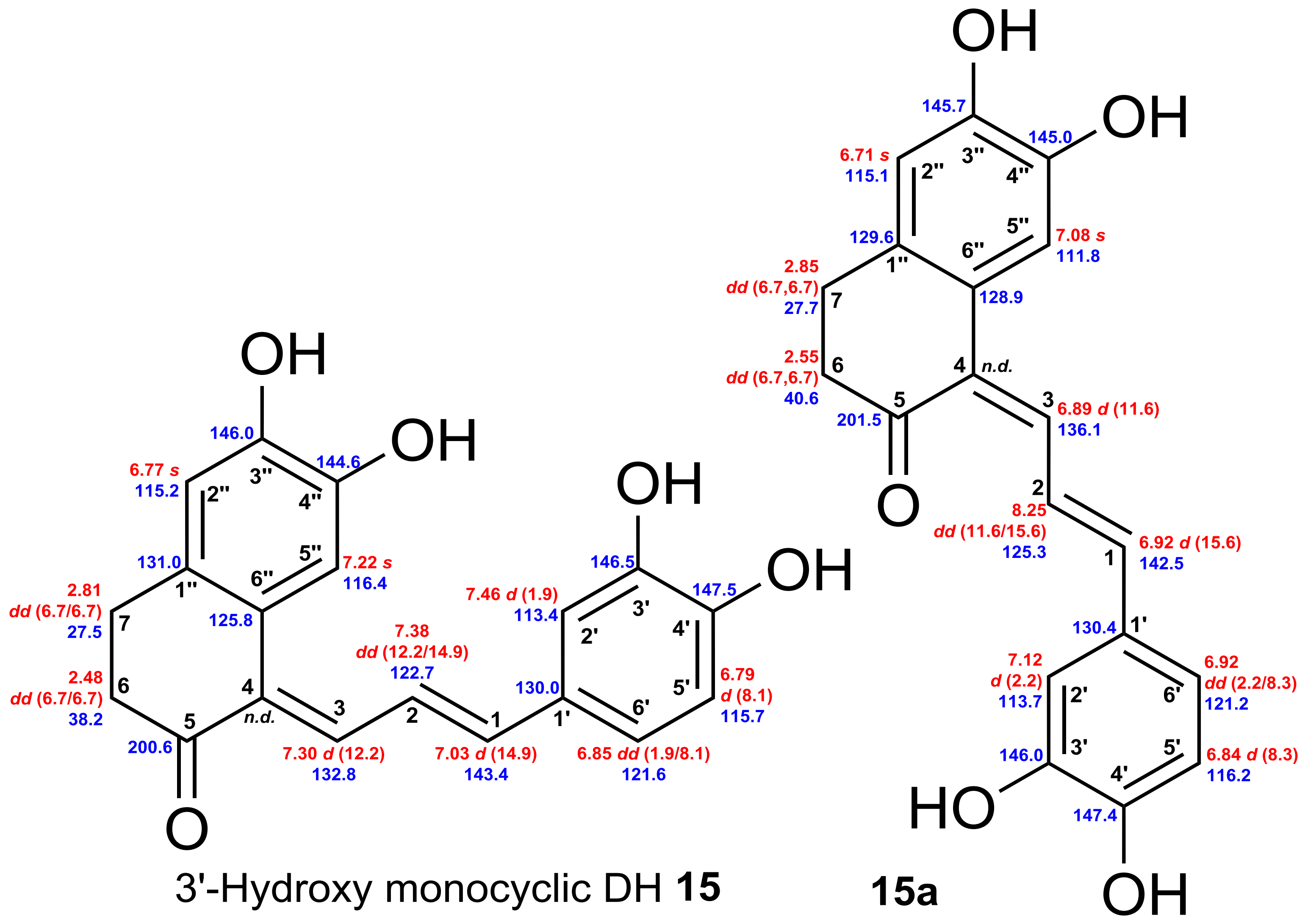


**Figure S65.** Chemical shifts of 3'-hydroxy monocyclic DH **15** and **15a** (*n.d.* not detected). Red: $^{1}H$ chemical shifts (δ ppm, *mult.*, $^{3}J_{HH}$ in Hz). Blue: $^{13}C$ chemical shifts (δ ppm).

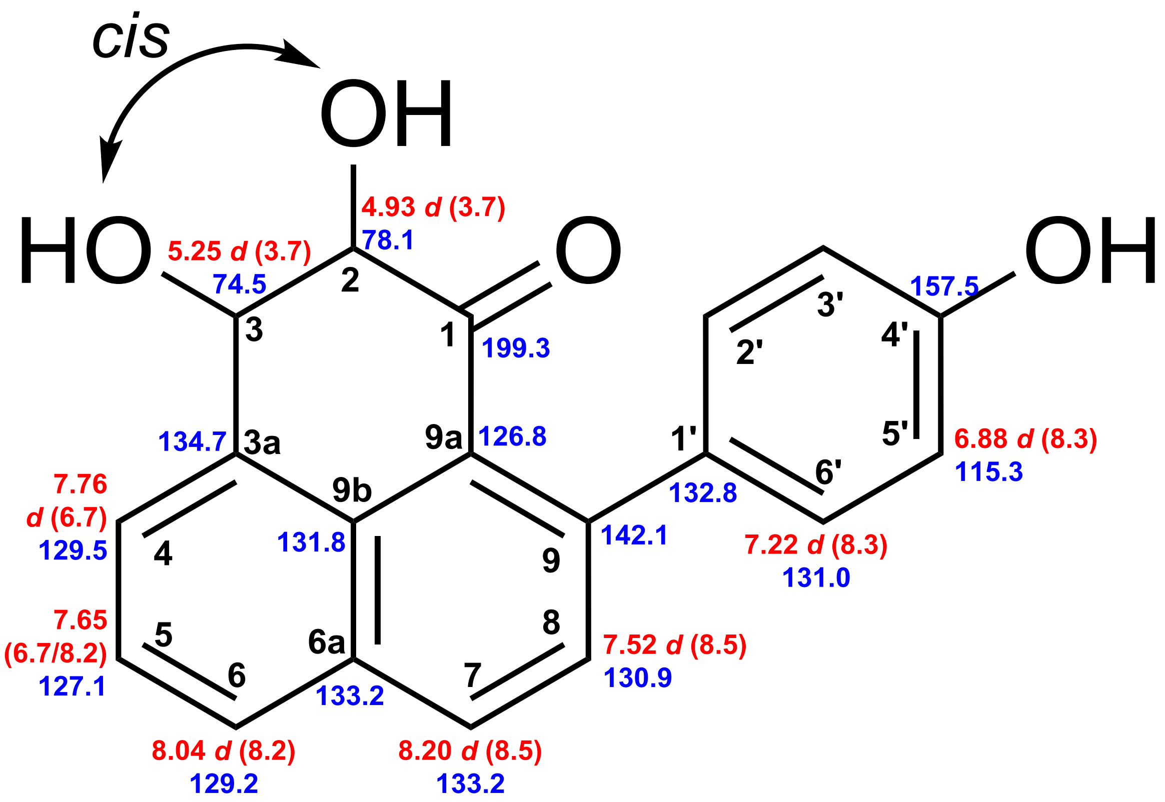


(−)-*cis*-2,3-dihydro-2,3-dihydroxy-9-(4′-hydroxyphenyl)phenalen-1-one

**Figure S66.** Chemical shifts of (−)-*cis*-2,3-dihydro-2,3-dihydroxy-9-(4'-hydroxyphenyl) phenalen-1-one. Red: $^{1}H$ chemical shifts (δ ppm, *mult*., $^{3}J_{HH}$ in Hz). Blue: $^{13}C$ chemical shifts (δ ppm).

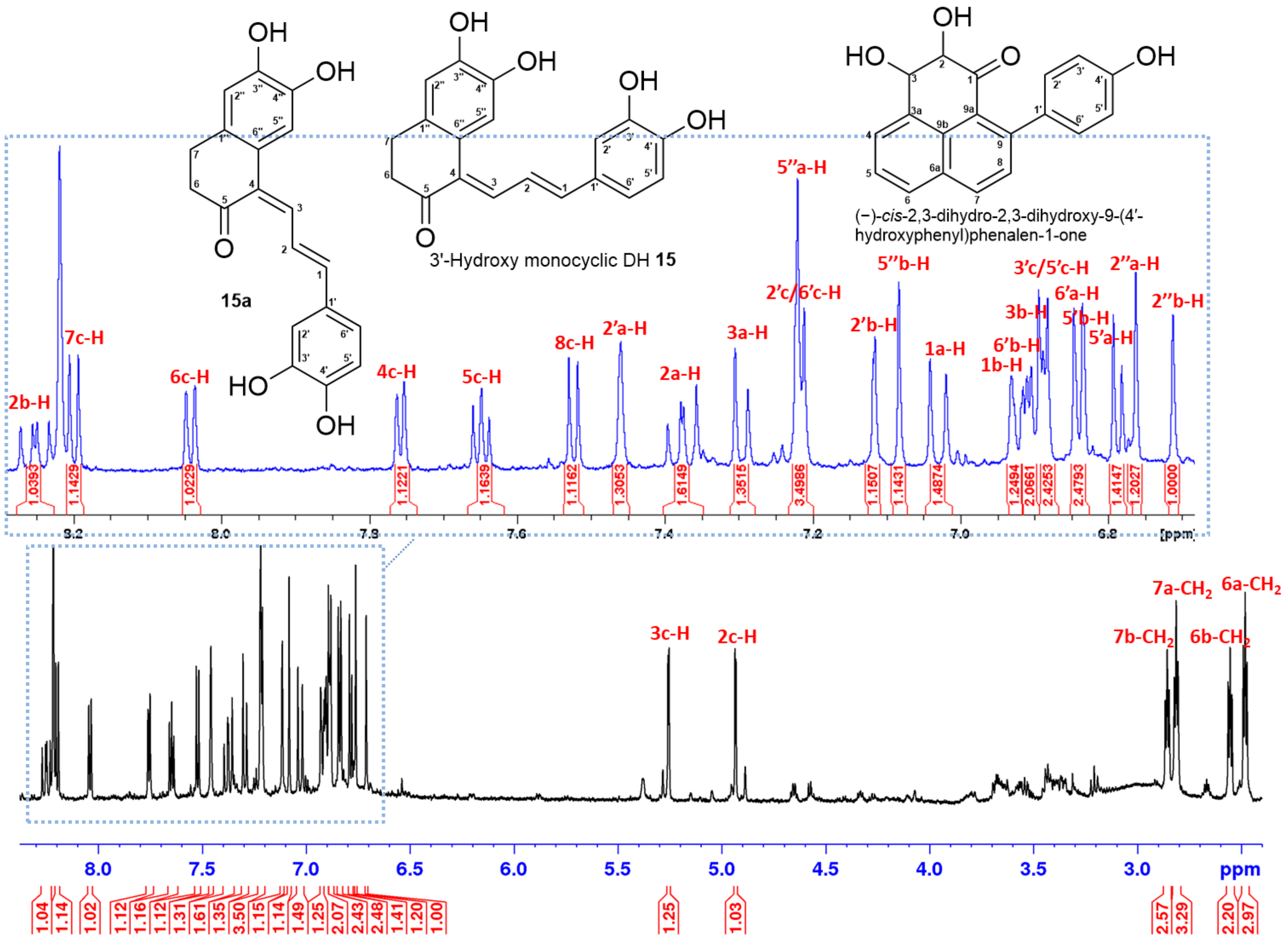


**Figure S67.** [1]H NMR spectrum (700 MHz, acetone-$d_6$) of 3'-hydroxy monocyclic DH **15** (assigned as a), **15a** (assigned as b), and (−)-*cis*-2,3-dihydro-2,3-dihydroxy-9-(4′-hydroxyphenyl)phenalen-1-one (assigned as c).

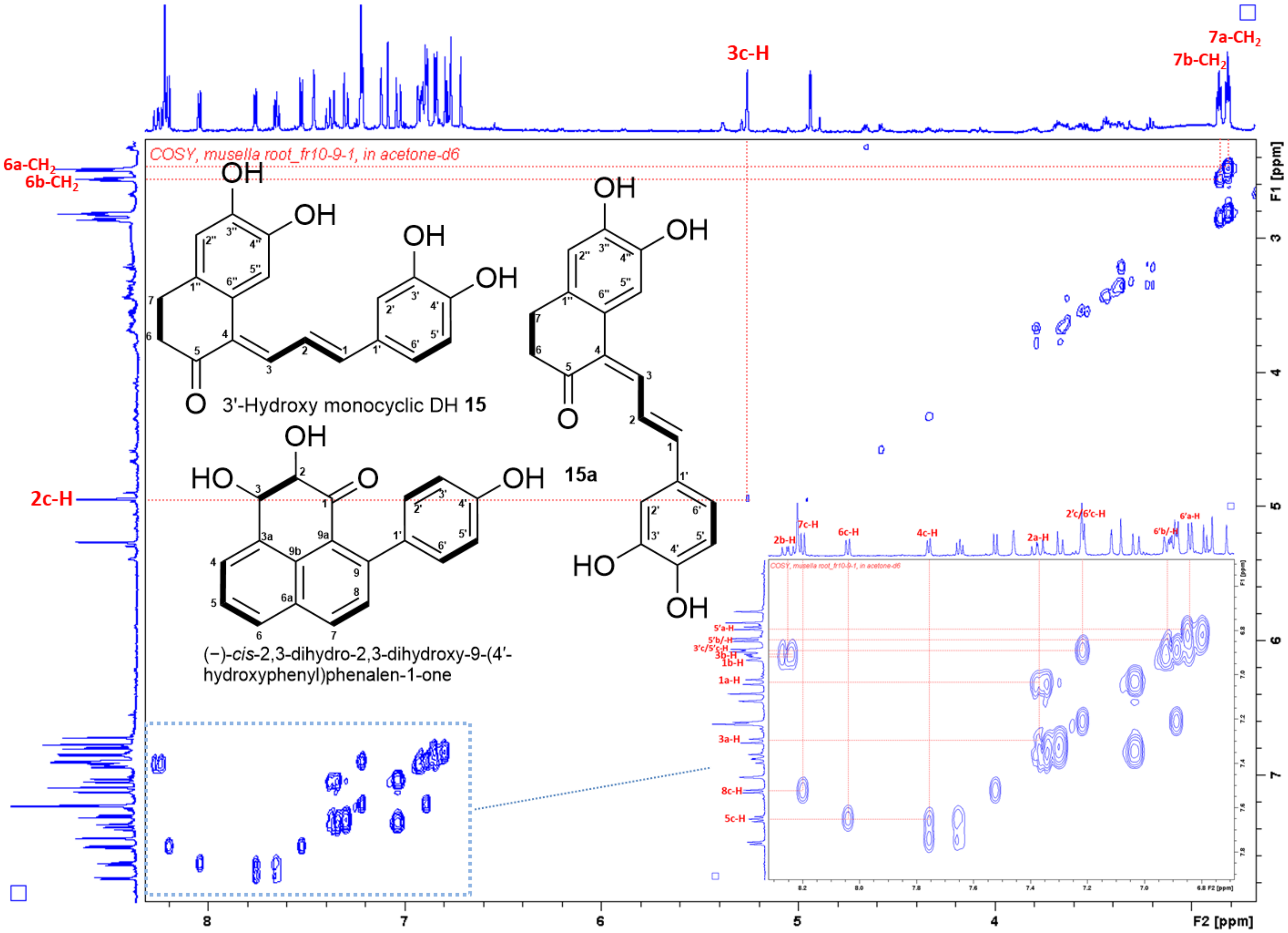


**Figure S68.** $^{1}H$–$^{1}H$ COSY spectrum of 3'-hydroxy monocyclic DH **15** (assigned as a), **15a** (assigned as b) and (−)-*cis*-2,3-dihydro-2,3-dihydroxy-9-(4′-hydroxyphenyl)phenalen-1-one (assigned as c) in acetone-$d_6$.

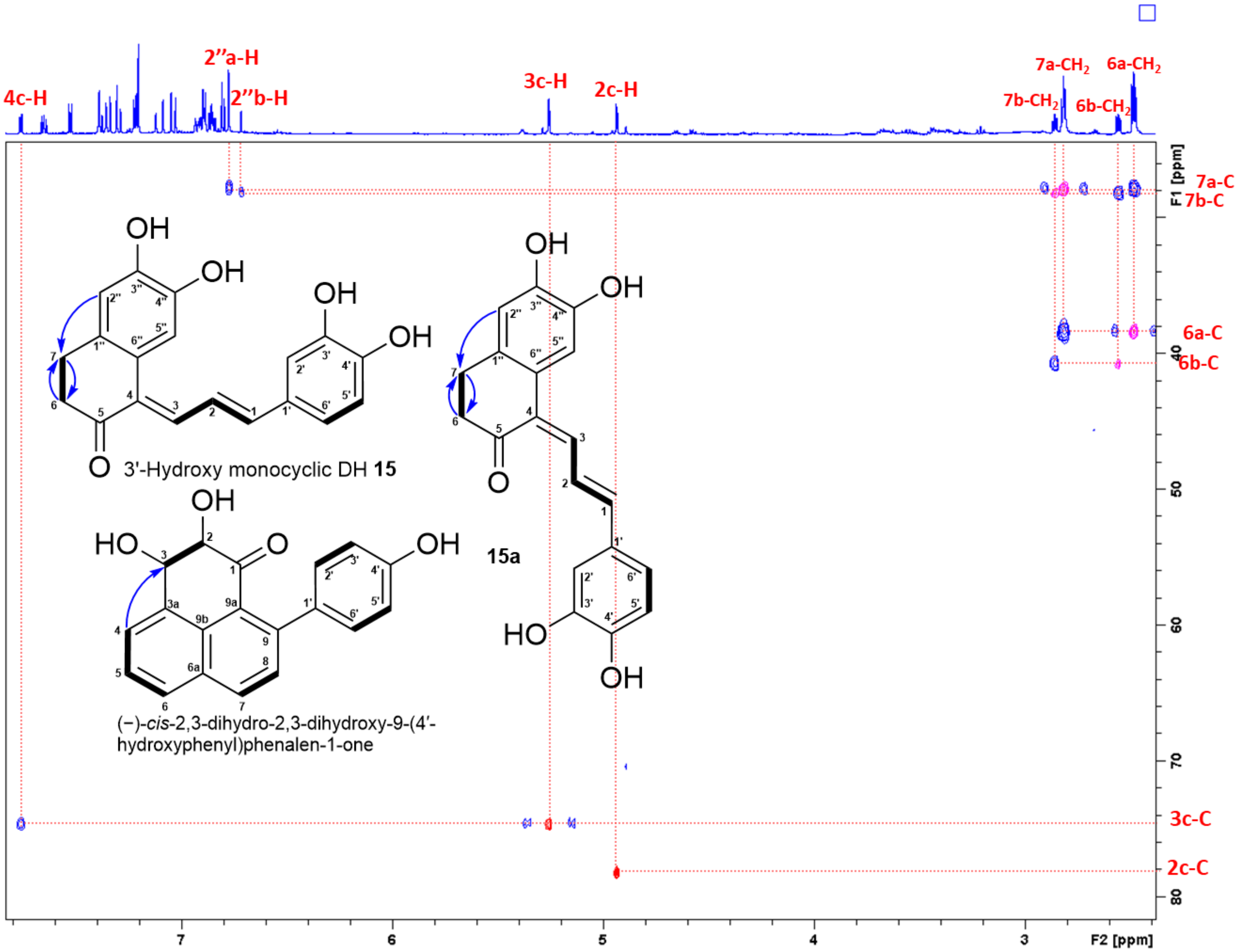


**Figure S69.** Superimposed HSQC and HMBC spectra of 3'-hydroxy monocyclic DH **15** (assigned as a)**, 15a** (assigned as b) and (−)-*cis*-2,3-dihydro-2,3-dihydroxy-9-(4′-hydroxyphenyl)phenalen-1-one (assigned as c) in acetone-$d_6$ (part-1).

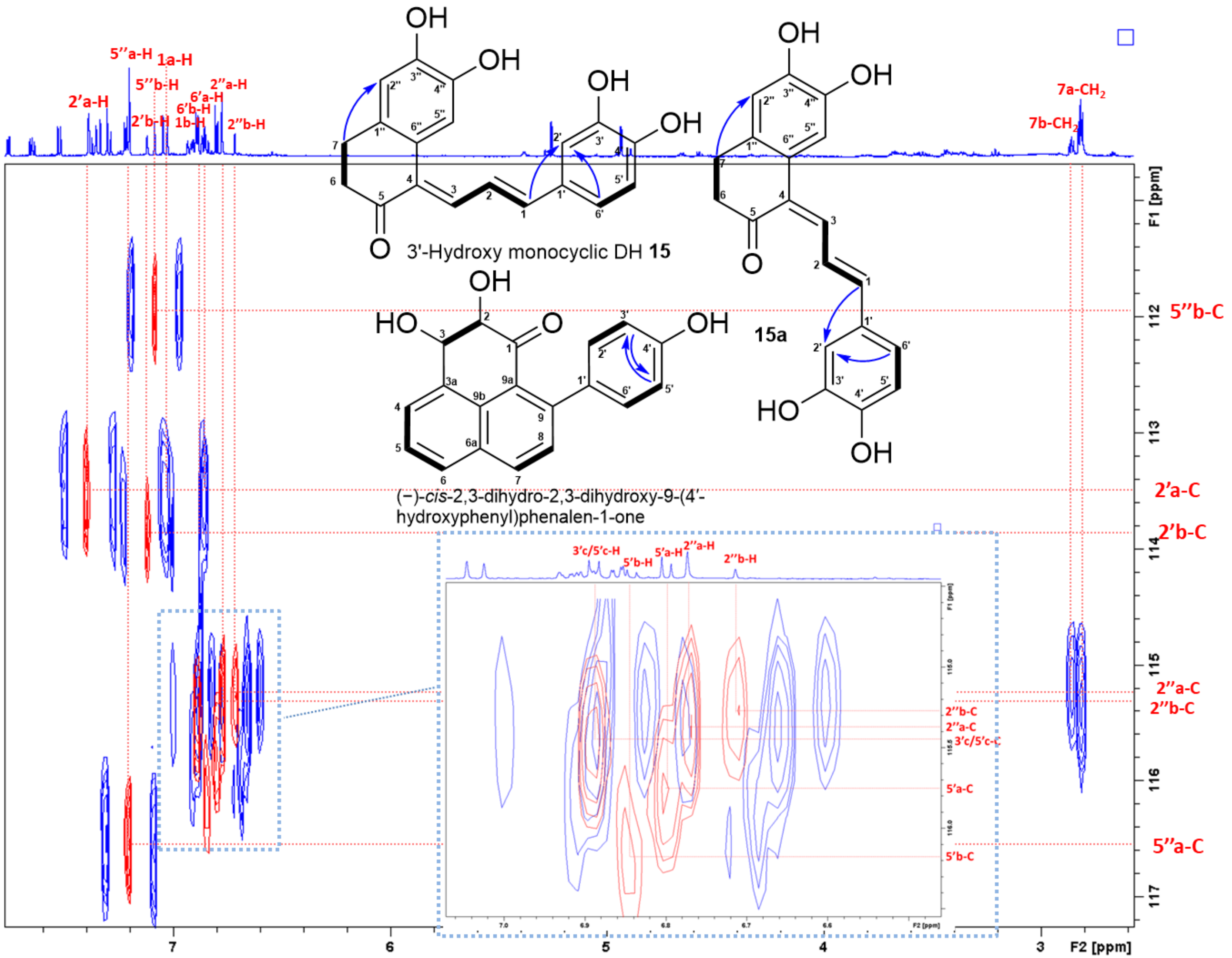


**Figure S70.** Superimposed HSQC and HMBC spectra of 3'-hydroxy monocyclic DH **15** (assigned as a)**, 15a** (assigned as b) and (−)-*cis*-2,3-dihydro-2,3-dihydroxy-9-(4′-hydroxyphenyl)phenalen-1-one (assigned as c) in acetone-$d_6$ (part-2).

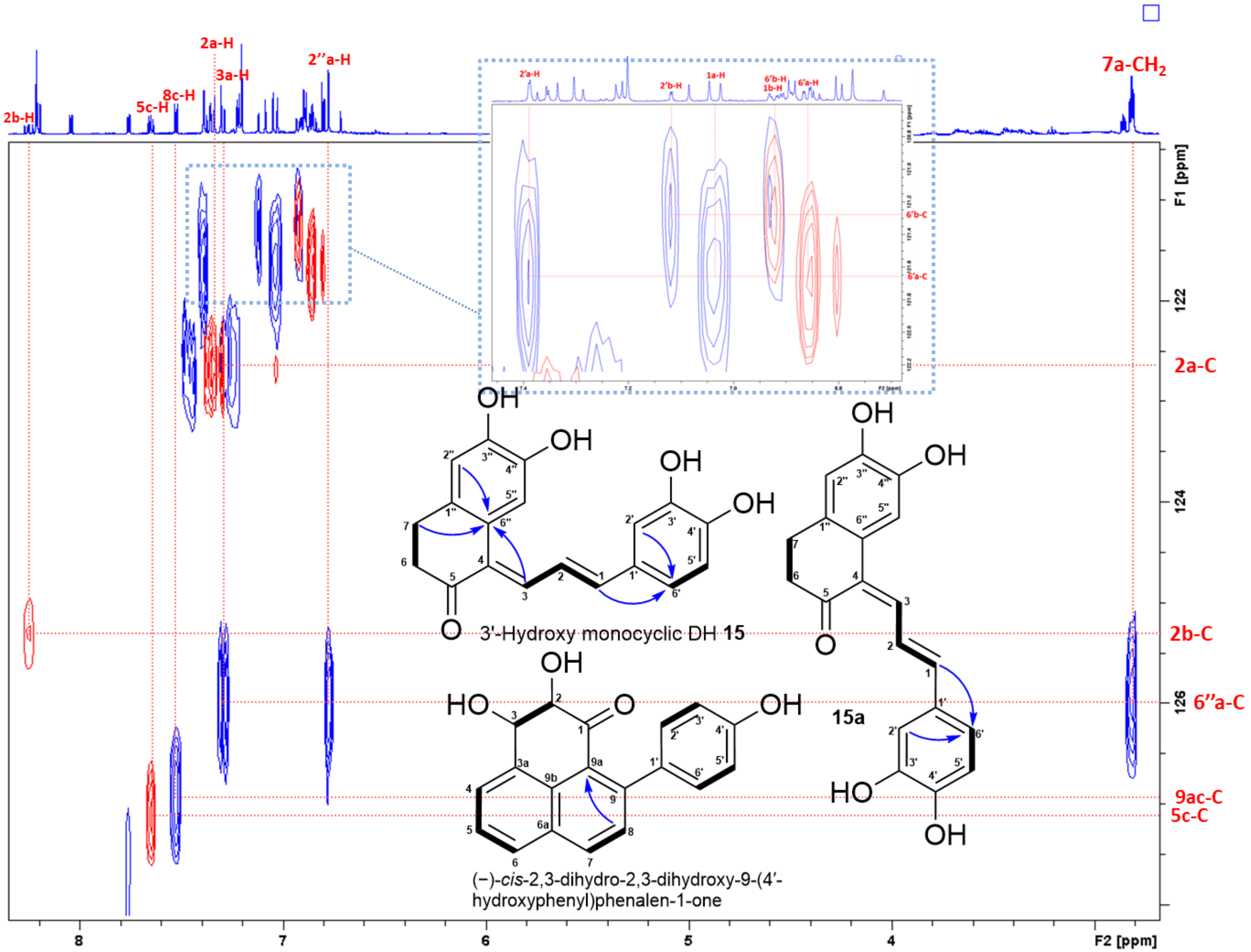


**Figure S71.** Superimposed HSQC and HMBC spectra of 3'-hydroxy monocyclic DH **15** (assigned as a)**, 15a** (assigned as b) and (−)-*cis*-2,3-dihydro-2,3-dihydroxy-9-(4′-hydroxyphenyl)phenalen-1-one (assigned as c) in acetone-$d_6$ (part-3).

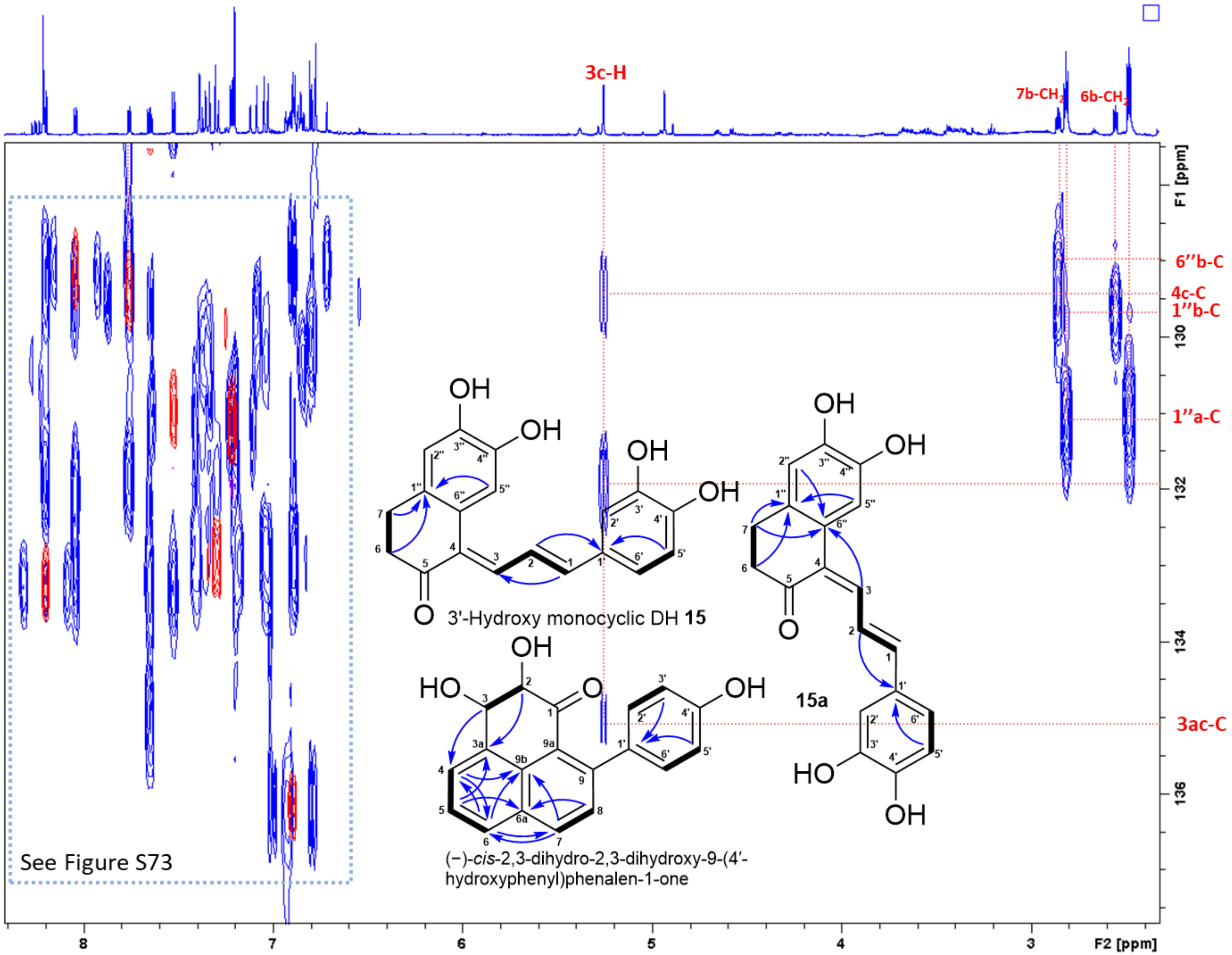


**Figure S72.** Superimposed HSQC and HMBC spectra of 3'-hydroxy monocyclic DH **15** (assigned as a)**, 15a** (assigned as b) and (−)-*cis*-2,3-dihydro-2,3-dihydroxy-9-(4′-hydroxyphenyl)phenalen-1-one (assigned as c) in acetone-$d_6$ (part-4).

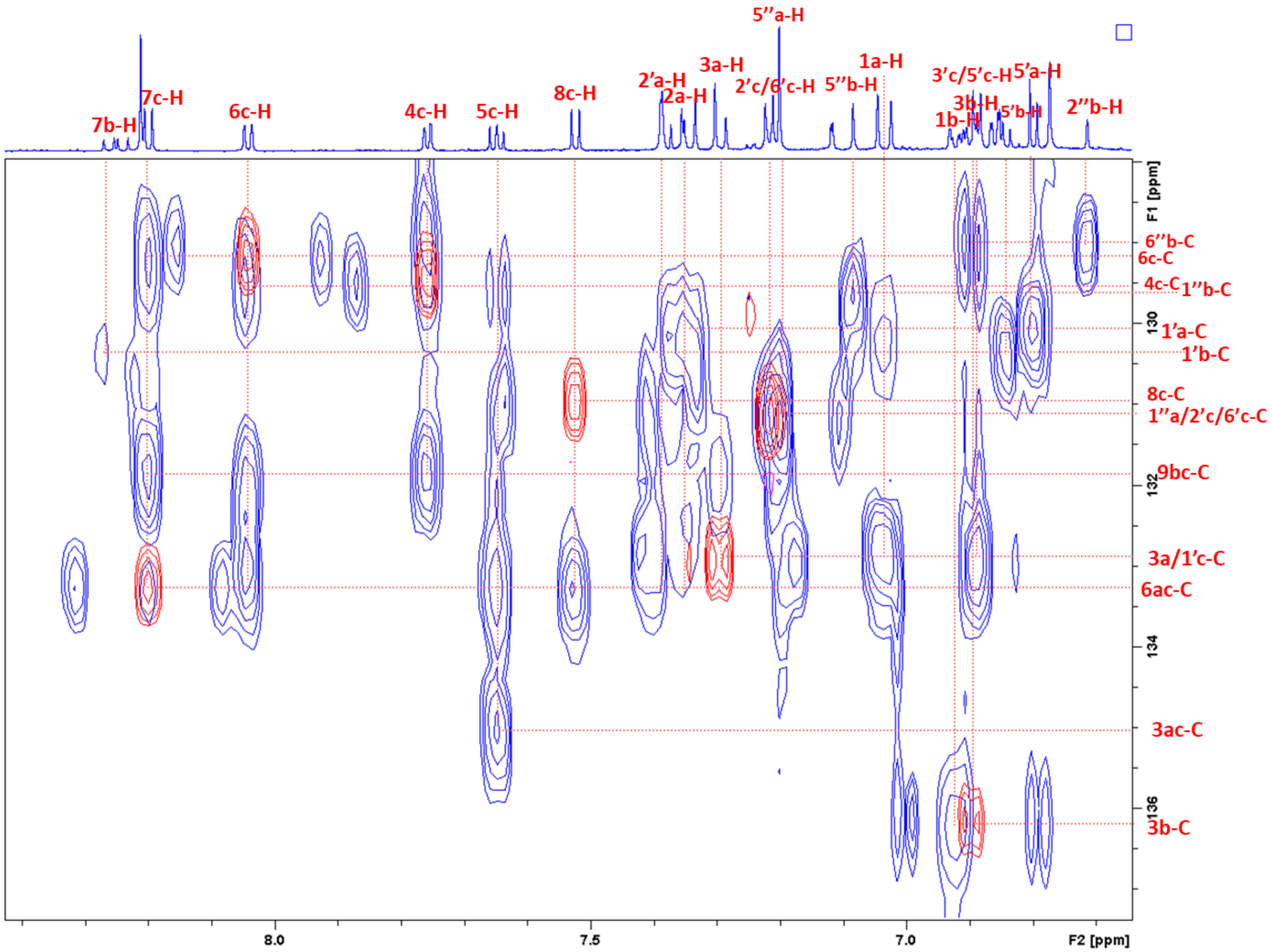


**Figure S73.** Superimposed HSQC and HMBC spectra of 3'-hydroxy monocyclic DH **15** (assigned as a)**, 15a** (assigned as b) and (−)-*cis*-2,3-dihydro-2,3-dihydroxy-9-(4′-hydroxyphenyl)phenalen-1-one (assigned as c) in acetone-$d_6$ (part-5).

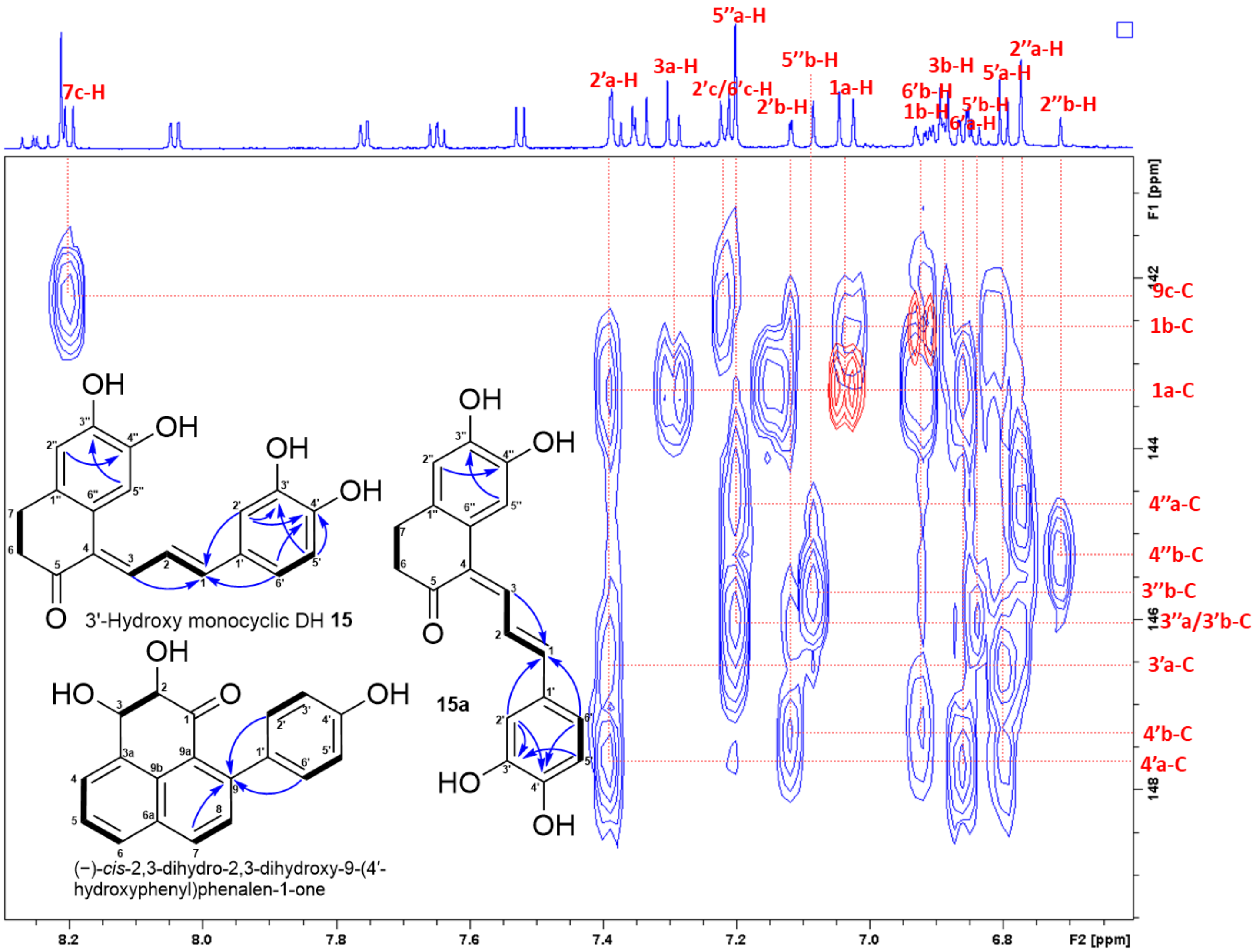


**Figure S74.** Superimposed HSQC and HMBC spectra of 3'-hydroxy monocyclic DH **15** (assigned as a)**, 15a** (assigned as b) and (−)-*cis*-2,3-dihydro-2,3-dihydroxy-9-(4′-hydroxyphenyl)phenalen-1-one (assigned as c) in acetone-$d_6$ (part-6).

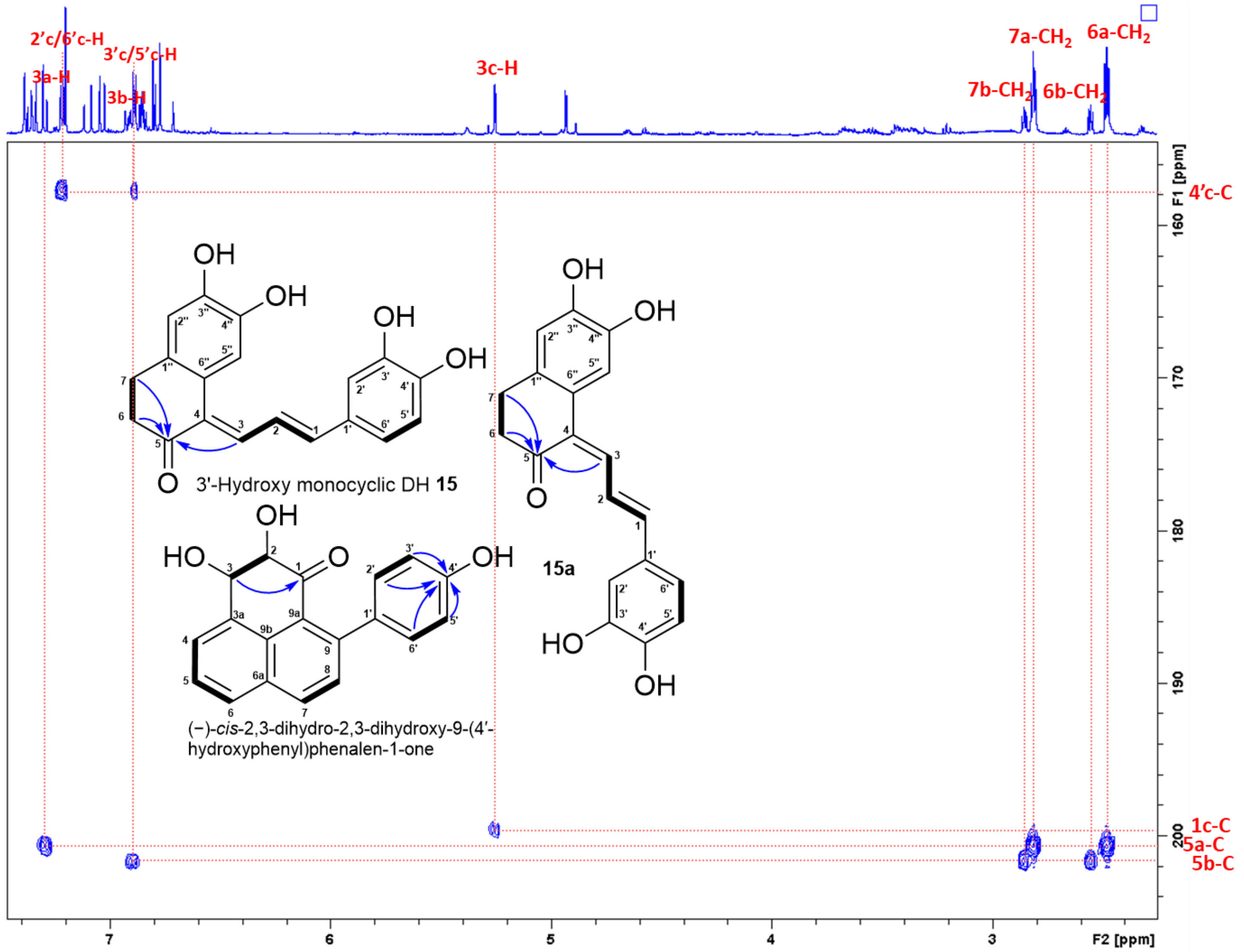


**Figure S75.** Superimposed HSQC and HMBC spectra of 3'-hydroxy monocyclic DH **15** (assigned as a)**, 15a** (assigned as b) and (−)-*cis*-2,3-dihydro-2,3-dihydroxy-9-(4′-hydroxyphenyl)phenalen-1-one (assigned as c) in acetone-$d_6$ (part-7).

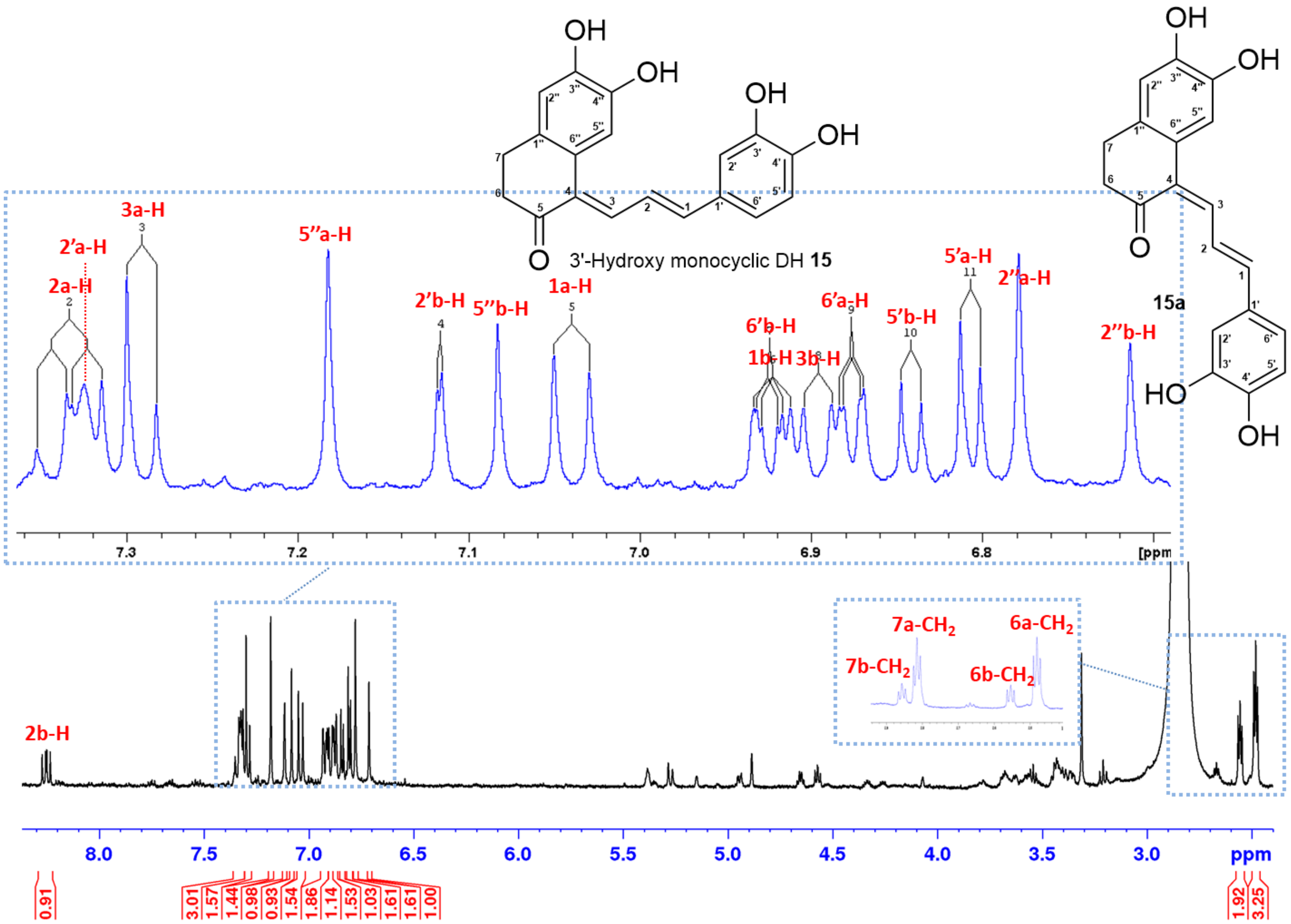


**Figure S76.** $^1$H NMR spectrum (700 MHz, acetone-$d_6$) of 3'-hydroxy monocyclic DH **15** (assigned as a) and **15a** (assigned as b).

#### Supplementary Tables

**Table S1.** $^1$H (700 MHz) and $^{13}$C (175 MHz) NMR Data for dihydroferuloyl-β-keto acid **6a** in $CD_3CN$.

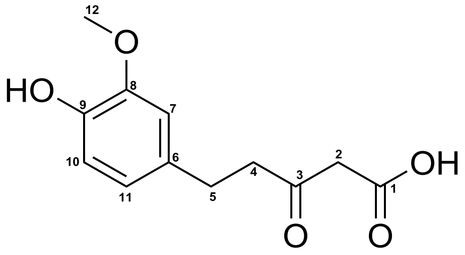


Dihydroferuloyl-β-keto acid **6a**

| No. | $\delta_H$, mult. (J in Hz) | $\delta_C$, type |
|---|---|---|
| 1 | – | 169.1, C |
| 2 | 3.42, bd | 49.3, $CH_2$ |
| 3 | – | 204.9, C |
| 4 | 2.76, dd (7.6/7.6) | 45.1, $CH_2$ |
| 5 | 2.82, dd (7.6/7.6) | 29.7, $CH_2$ |
| 6 | – | 134.1, C |
| 7 | 6.79, d (1.7) | 112.9, CH |
| 8 | – | 147.8, C |
| 9 | – | 145.1, C |
| 10 | 6.70, d (8.1) | 115.4, CH |
| 11 | 6.63, dd (1.7/8.1) | 121.4, CH |
| 12 | 3.81, s | 56.4, $CH_3$ |

**Table S2.** $^{1}$H (700 MHz) and $^{13}$C (175 MHz) NMR Data for monocyclic DH **13** and **13a** and 3'-hydroxy monocyclic DH **15** and **15a** in acetone-$d_6$ (*n.d.* not detected).

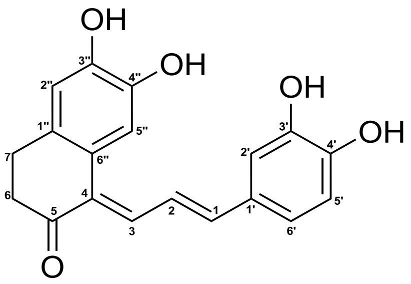


Monocyclic DH **13** R = H
3'-Hydroxy monocyclic DH **15** R = OH

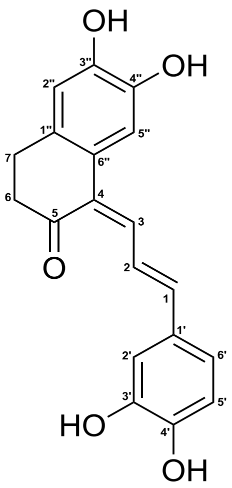


**13a** R = H
**15a** R = OH

| No. | **15** | | **15a** | | **13** | | **13a** | |
|---|---|---|---|---|---|---|---|---|
| | **$\delta_H$, mult. (*J* in Hz)** | **$\delta_C$, type** | **$\delta_H$, mult. (*J* in Hz)** | **$\delta_C$, type** | **$\delta_H$, mult. (*J* in Hz)** | **$\delta_C$, type** | **$\delta_H$, mult. (*J* in Hz)** | **$\delta_C$, type** |
| 1 | 7.12, *d* (13.2) | 142.7, CH | 6.98, *d* (15.5) | 142.3, CH | 7.03, *d* (14.9) | 143.4, CH | 6.92, *d* (15.6) | 142.5, CH |
| 2 | 7.32, *dd* (11.2/13.2) | 122.8, CH | 8.29, *dd* (11.5/15.5) | 125.4, CH | 7.38, *dd* (12.2/14.9) | 122.7, CH | 8.25, *dd* (11.6/15.6) | 125.3, CH |
| 3 | 7.29, *d* (11.2) | 133.0, CH | 6.91, *d* (11.5) | 136.3, CH | 7.30, *d* (11.2) | 132.8, CH | 6.89, *d* (11.6) | 136.1, CH |
| 4 | | *n.d.* | | *n.d.* | | *n.d.* | | *n.d.* |
| 5 | | 200.8, C | | 201.7, C | | 200.6, C | | 201.5, C |
| 6 | 2.47, *dd* (6.7/6.7) | 38.3, $CH_2$ | 2.55, *dd* (6.7/6.7) | 40.8, $CH_2$ | 2.48, *dd* (6.7/6.7) | 38.2, $CH_2$ | 2.55, *dd* (6.7/6.7) | 40.6, $CH_2$ |
| 7 | 2.83, *dd* (6.7/6.7) | 27.8, $CH_2$ | 2.85, *dd* (6.7/6.7) | 28.2, $CH_2$ | 2.81, *dd* (6.7/6.7) | 27.5, $CH_2$ | 2.85, *dd* (6.7/6.7) | 27.7, $CH_2$ |
| 1' | | 129.6, C | | 129.6, C | | 130.0, C | | 130.4, C |
| 2' | 7.45, *d* (8.4) | 129.7, CH | 7.44, *d* (8.4) | 129.7, CH | 7.46, *d* (1.9) | 113.4, CH | 7.12, *d* (2.2) | 113.7, CH |
| 3' | 6.87, *d* (8.4) | 116.7, CH | 6.88, *d* (8.4) | 116.7, CH | | 146.5, C | | 146.0, C |
| 4' | | 159.3 C | | 159.3, C | | 147.5 C | | 147.4 C |
| 5' | 6.87, *d* (8.4) | 116.7, CH | 6.88, *d* (8.4) | 116.7, CH | 6.79, *d* (8.1) | 115.7, CH | 6.84, *d* (8.3) | 116.2, CH |
| 6' | 7.45, *d* (8.4) | 129.7, CH | 7.44, *d* (8.4) | 129.7, CH | 6.85, *dd* (1.9/8.1) | 121.6, CH | 6.92, *dd* (2.2/8.3) | 121.2, CH |

**Table S2.** continued

| No. | 15 | | 15a | | 13 | | 13a | |
|---|---|---|---|---|---|---|---|---|
| | $\delta_H$, mult. (*J* in Hz) | $\delta_C$, type | $\delta_H$, mult. (*J* in Hz) | $\delta_C$, type | $\delta_H$, mult. (*J* in Hz) | $\delta_C$, type | $\delta_H$, mult. (*J* in Hz) | $\delta_C$, type |
| 1'' | | 131.5, C | | 129.8, C | | 131.0, C | | 129.6, C |
| 2'' | 6.80, *s* | 115.9, CH | 6.71, *s* | 115.4, CH | 6.77, *s* | 115.2, CH | 6.71, *s* | 115.1, CH |
| 3'' | | 145.8, C | | 145.8, C | | 146.0, C | | 145.7, C |
| 4'' | | 144.7, C | | 145.5, C | | 144.6, C | | 145.0, C |
| 5'' | 7.11, *s* | 116.6, CH | 7.09, *s* | 112.1, CH | 7.22, *s* | 116.4, CH | 7.08, *s* | 111.8, CH |
| 6'' | | 126.1, C | | 129.1, C | | 125.8, C | | 128.9, C |

**Table S3.** $^{1}H$ (700 MHz) and $^{13}C$ (175 MHz) NMR Data for 4'-hydroxylachnanthocarpone **14** in acetone-$d_6$.

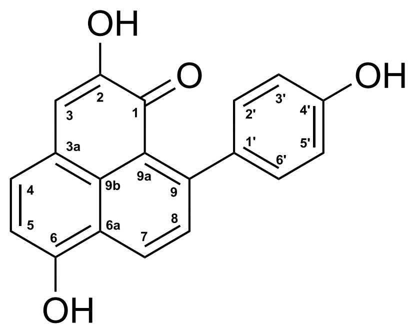


4'-Hydroxylachnanthocarpone **14**

| No. | $\delta_H$, mult. (J in Hz) | $\delta_C$, type |
|---|---|---|
| 1 | – | 180.0, C |
| 2 | – | 149.3, C |
| 3 | 7.08, s | 113.5, CH |
| 3a | – | 121.2, C |
| 4 | 7.68, d (7.8) | 133.2, CH |
| 5 | 7.05, d (7.8) | 110.2, CH |
| 6 | – | 157.7, C |
| 6a | – | 124.3, C |
| 7 | 8.71, d (8.3) | 130.1, CH |
| 8 | 7.59, d (8.3) | 131.2, CH |
| 9 | – | 149.9, C |
| 9a | – | 124.6, C |
| 9b | – | 126.7, C |
| 1’ | – | 134.6, C |
| 2’/6’ | 7.26, d (6.5) | 130.6, CH |
| 3’/5’ | 6.91, d (6.5) | 115.5, CH |
| 4’ | – | 157.4, C |

**Table S6.** Composition of *in vitro* enzyme assays.

| Enzyme conbinations (each 1 µM) | Starter substrates | Extender substrate (100 µM) | Cofactor (1 mM) | Workup |
|---|---|---|---|---|
| *Ml*DCS1 | *p*-coumaroyl-CoA **1**, feruloyl-CoA **2**, cinnamoyl-CoA, caffeoyl-CoA, *p*-dihydrocoumaroyl-CoA (100 µM) | malonyl-CoA | – | 5 µL of 10 M NaOH, then incubated at 65 °C for 10 min |
| *Ml*DCS2 | *p*-coumaroyl-CoA **1**, feruloyl-CoA **2**, cinnamoyl-CoA, caffeoyl-CoA, p-dihydrocoumaroyl-CoA (100 µM) | malonyl-CoA | – | 5 µL of 10 M NaOH, then incubated at 65 °C for 10 min |
| *Ml*DCS1, *Ml*CURS1 | *p*-coumaroyl-CoA **1** (200 µM) | malonyl-CoA | – | – |
| *Ml*DCS1, *Ml*CURS2 | *p*-coumaroyl-CoA **1** (200 µM) | malonyl-CoA | – | – |
| *Ml*DCS2, *Ml*CURS1 | feruloyl-CoA **2** (200 µM) | malonyl-CoA | – | – |
| *Ml*DCS2, *Ml*CURS2 | feruloyl-CoA **2** (200 µM) | malonyl-CoA | – | – |
| *Ml*DBR | *p*-coumaroyl-CoA **1**, feruloyl-CoA **2** (100 µM) | – | NADPH /NADH | 5 µL of 10 M NaOH, then incubated at 65 °C for 10 min |
| *Ml*DBR | bisdemethoxycurcumin **9**, curcumin **10** (100 µM) | – | NADPH /NADH | – |
| *Ml*DCS1, *Ml*DBR | *p*-coumaroyl-CoA **1** (100 µM) | malonyl-CoA | NADPH /NADH | – |
| *Ml*DCS2, *Ml*DBR | feruloyl-CoA **2** (100 µM) | malonyl-CoA | NADPH /NADH | incubated for 1, 1.5, 2, 3, and 10 hours |
| *Ml*DCS1, *Ml*DBR, *Ml*CURS1 | *p*-coumaroyl-CoA **1** (200 µM) | malonyl-CoA | NADPH | – |
| *Ml*DCS2, *Ml*DBR, *Ml*CURS2 | feruloyl-CoA **2** (200 µM) | malonyl-CoA | NADPH | – |
